# BioMedGraphica: An All-in-One Platform for Joint Textual Biomedical Prior Knowledge and Numeric Graph Generation

**DOI:** 10.1101/2024.12.05.627020

**Authors:** Heming Zhang, Shunning Liang, Tim Xu, Wenyu Li, Di Huang, Yuhan Dong, Guangfu Li, J. Philip Miller, S. Peter Goedegebuure, Marco Sardiello, Jonathan Cooper, William Buchser, Patricia Dickson, Ryan C. Fields, Carlos Cruchaga, Yixin Chen, Michael Province, Philip Payne, Fuhai Li

**Author notes:** Co-first authors (Heming Zhang, Shunning Liang, Tim Xu).

## Abstract

Multi-omic data analysis is essential for scientific discovery in precision medicine. However, translating statistical results of omic data analysis into novel scientific hypothesis remains a significant challenge. Human experts must manually review analysis results and generate new hypothesis based on extensive and inter-connected biomedical prior knowledge, which is subjective and not scalable. While large language models (LLMs) can accelerate the discovery, their reasoning improves when grounded in structured, auditable and comprehensive biomedical prior knowledge. Biomedical knowledge, however, is scattered across heterogeneous databases that use diverse and inconsistent nomenclature systems, making it difficult to integrate resources into a unified format for scalable analysis. This fragmentation limits the ability of AI systems to fully leverage biomedical data for scientific discovery. To address these challenges, we developed ***BioMedGraphica***, an all-in-one platform that harmonizes fragmented biomedical resources by integrating 11 entity types and 30 relation types from 43 databases into a unified knowledge graph containing **2,306,921** entities and **27,232,091** relations. In addition, to the best of our knowledge, this is the first work to propose a novel **Textual-Numeric Graph (TNG)** data-structure for multi-omics data analysis. In TNG, textual information captures prior biological knowledge (e.g., transcription start sites, functions, mechanisms), while numeric values represent quantitative biomedical features, and the integrated relations can help uncover mechanisms. By bridging prior knowledge with user-specific data, TNG is a novel and ideal data-structure for the development of graph foundation models, with the potential to improve prediction performance and interpretability, while also augmenting LLMs by supplying graph-structured mechanistic context to strengthen reasoning. The details for BioMedGraphica code can be accessed by github link: https://github.com/FuhaiLiAiLab/BioMedGraphica and BioMedGraphica knowledge graph data can be downloaded from huggingface dataset: https://huggingface.co/datasets/FuhaiLiAiLab/BioMedGraphica

## 1 Introduction

In recent years, the exponential growth of omic datasets^1–10^ has created unprecedented opportunities to advance biomedical research, precision medicine, improve clinical decision-making, and accelerate drug discovery. While the convergence of artificial intelligence (AI) models with massive omic datasets is revolutionizing the paradigm of scientific discovery in precision medicine, this transformation is still in its infancy^9,11^. Especially, in omic data analysis, the first step is to identify a set of differentially expressed targets and enriched signaling pathways or biological functions; and then followed by the human manual review with the support of online search of extensive and inter-connected prior knowledge to generate expert-specific scientific hypotheses to be further evaluated, which is subjective and not scalable. Though Large language models (LLMs) and agentic AI models^12–16^ are transforming scientific discovery due to their ability to interpret and reason with human-readable textual information, their reasoning improves when grounded in structured, auditable evidence^17–19^. However, the landscape of biomedical knowledge remains highly fragmented, with essential information dispersed across a multitude of publications, databases, and proprietary datasets. This fragmentation presents significant challenges as different sources often employ inconsistent nomenclature and terminologies, hindering effective data integration^20^. The vast scope of biomedical data, from genes and proteins to clinical phenotypes and diseases, complicates the development of unified solutions, particularly in terms of entity matching and data harmonization. As a result, several knowledge graph systems have been developed to integrate this extensive array of data resources, aiming to merge information across the spectrum of biomedical domains^21–26^. However, existed knowledge graphs struggle to reconcile entities across these diverse, heterogeneous datasets, which are often noisy, inconsistent, and formatted in various ways. Without harmonizing nomenclature systems over multiple resources in biomedical domains, most current systems lack comprehensive coverage of biomedical resources, which leads to failure on implementing efficient matching algorithms, and rely heavily on manual work, making them less efficient for large-scale data integration. As a result, converting unstructured data into formats suitable for graph-based artificial intelligence (AI) models^27–34^ becomes an onerous task (e.g., mosGraphGen^35^, IntergAO^36^) require extensive manual curation.

To address these challenges, ***BioMedGraphica*** was developed as an advanced platform that transforms the integration and utilization of biomedical data. By integrating data from **43** high-quality biomedical databases, we unify **11** key biomedical entity types—ranging from promoters, genes, transcripts, proteins, signaling pathways, metabolites and microbiotas to clinical phenotypes, diseases, and drugs—and **30** relations / edge types into a cohesive knowledge graph, resulting in **2,306,921** entities and **27,232,091** relations. With harmonizing over multiple knowledge bases, this study provides one of the most comprehensive biomedical knowledge graphs available today, enabling large-scale exploration of biological and clinical relationships. Meanwhile, for real life scenarios of biomedical application, we removed the isolated entities and reconstructed this database with 834,809 entities and 27,232,091 relations, called ***BioMedGraphica-Conn***. Built on the harmonized nomenclature system, a core innovation of BioMedGraphica is its entity-matching framework, which combines hard matching, based on standardized identifiers across resources, with soft matching, powered by language models such as BioBERT^37^, to generate embeddings that rank potential matches across heterogeneous datasets. This approach enhances entity recognition by providing more accurate and flexible integration compared to traditional rule-based methods, enabling robust, scalable alignment that reduces manual curation while maintaining precision. These matching capabilities are implemented in a user-friendly web interface with an intuitive GUI, accessible at https://app.biomedgraphica.org/. This allows researchers, clinicians, and data scientists to input heterogeneous biomedical datasets and receive well-integrated, structured outputs that are AI-ready, including a novel data format we introduce: **textual-numeric graph (TNG)**. In TNG, textual information (biomedical prior knowledge) conveys known biological functions, while numeric values capture the quantitative features of biomedical entities across multiple levels, from molecular to clinical, along with their corresponding relations in the knowledge graph. By bridging prior knowledge with user-specific data, TNG provides a robust foundation for advancing graph foundation models. Prior work such as GraphSeqLM^38^ has demonstrated that integrating textual features with omic graphs improves prediction accuracy, and BioMedGraphica extends this by offering a systematic and interpretable framework for promising foundation of better model performance and mechanistic interpretability (facilitating discoveries in biomarker identification, drug target exploration, disease etiology, and personalized therapeutic strategies). Moreover, TNG can augment large language models by supplying graph-structured mechanistic context, thereby strengthening reasoning and supporting the development of next-generation agentic AI systems^17–19,39^. Designed with low coupling in entity space and a streamlined pipeline, the platform supports continuous updates and integration of new resources, ensuring its relevance as biomedical knowledge expands. Together, these advances position BioMedGraphica as both a practical tool of translational science and a cornerstone for the era of foundation models and agentic AI in biomedicine, creating an ***all-in-one*** platform for graph foundation models that pave the way for improving prediction performance and interpretability while augmenting LLMs with mechanistic context to drive scalable, evidence-based scientific discoveries in biomedicine.

## 2 Methods

### 2.1 Overview of Data Resources Used in *BioMedGraphica*

To enable robust biomedical entity recognition and relation extraction, BioMedGraphica integrates a diverse and well-curated collection of biomedical data resources. These resources span both entity databases, which catalog structured identifiers and metadata for various biological and chemical entities, and relation databases, which capture known associations and interactions across biological systems. Together, these datasets form the foundation for harmonizing multi-scale biomedical knowledge, supporting the accurate construction of structured graphs for downstream applications in biomedical discovery and precision medicine. The following subsections describe the data collection strategies, integration methods, and scope of both entity and relation datasets utilized in BioMedGraphica.

#### 2.1.1 Entity Databases Introduction and Collection

A wide range of reputable biomedical databases were utilized to gather and integrate various types of data related to genes, transcripts, proteins, and other biomedical entities (see **Figure 2**). This comprehensive integration ensured data consistency and accuracy, creating a unified framework essential for research. As shown in **Table 1**, the total number of entries in the original data file from each respective database were listed. For the ChEBI (Chemical Entities of Biological Interest) database, we utilized two primary datasets: one provided the mapping between ChEBI IDs and their corresponding InChI, while the other contained the mapping between ChEBI IDs and another database. For UNII (Unique Ingredient Identifier) database, our source data was obtained from two sources: one from PubChem, and the other provided by the FDA. For SILVA, we selected the LSU and SSU datasets. In **Table 1**, we not only present the total number of entries after merging the two files but also indicate the total number of rows for each dataset in parentheses. Similarly, for GTDB (Genome Taxonomy Database), we selected the data for both Archaea and Bacteria. The total number of entries after merging is indicated, with the individual row counts for each dataset provided in parentheses. Below is an expanded description of the databases used and the extracted data (check data overall details in **Table 1** and **Table S1**).

**Table 1.** Overview of Entity Databases.

| Database Names | Full Names | Entity Types | Total Number of Rows |
| --- | --- | --- | --- |
| Ensembl<sup>40</sup> | Ensembl | Gene | 86,406 |
|  |  | Transcript | 451,959 |
|  |  | Protein | 157,628 |
| OMIM<sup>41</sup> | Online Mendelian Inheritance in Man | Gene | 29,021 |
| HGNC<sup>42</sup> | HUGO Gene Nomenclature Committee | Gene | 43,916 |
| NCBI<sup>43</sup> | National Center for Biotechnology Information | Gene | 193,340 |
|  |  | Microbiota | 2,631,459 |
| RefSeq<sup>44</sup> | NCBI - Reference Sequence Database | Gene | 845,230 |
|  |  | Transcript | 19,404 |
|  |  | Protein | 376,561 |
| RNACentral<sup>45</sup> | RNACentral | Transcript | 66,789 |
| UniProt<sup>46</sup> | Universal Protein Resource | Protein | 20,417 |
| Reactome<sup>47</sup> | Reactome | Pathway | 2,751 |
| KEGG<sup>48</sup> | Kyoto Encyclopedia of Genes and Genomes | Pathway | 365 |
| WikiPathways<sup>49</sup> | WikiPathways | Pathway | 1,534 |
| Pathway Ontology<sup>50</sup> | Pathway Ontology | Pathway | 2,677 |
| ComPath<sup>51</sup> | Comparative Pathology Platform of the University of Bern | Pathway | 1,592 |
| HMDB<sup>52</sup> | Human Metabolome Database | Metabolite | 217,920 |
| ChEBI<sup>53</sup> | Chemical Entities of Biological Interest | Metabolite | 5,757 |
|  |  |  | 392,733 |
| SILVA<sup>54</sup> | SILVA | Microbiota | 2,214,227(227,318; 2,224,690) |
| Greengenes<sup>55</sup> | Greengenes | Microbiota | 1,144,866 |
| RDP<sup>56</sup> | Ribosomal Database Project | Microbiota | 10,302 |
| GTDB<sup>57</sup> | Genome Taxonomy Database | Microbiota | 596,859(12,477; 584,382) |
| CTD<sup>58</sup> | The Comparative Toxicogenomics Database | Exposure | 3,539 / 224,304 |
| HPO<sup>59</sup> | Human Phenotype Ontology | Phenotype | 19,533 |
| UMLS<sup>60</sup> | Unified Medical Language System | Phenotype | 14,036,386 |
|  |  | Disease | 16,704,679 |
| ICD10 / ICD11<sup>61,62</sup> | International Classification of Diseases | Disease | 12,597 / 36,044 |
| DO<sup>63</sup> | Disease Ontology | Disease | 38,212 |
| MeSH<sup>64</sup> | Medical Subject Headings | Disease | 5,056 |
| SNOMED-CT<sup>65</sup> | Systematized Nomenclature of Medicine Clinical Terms | Disease | 1,679,595 |
| Mondo<sup>66</sup> | Mondo | Disease | 134,286 |
| PubChem<sup>67</sup> | Public Chemical Databases | Drug | 123,357 |
| NDC<sup>68</sup> | National Drug Code | Drug | 107,980 |
| UNII<sup>69</sup> | Unique Ingredient Identifier | Drug | 159,376 |
|  |  |  | 152,870 |
| DrugBank<sup>70</sup> | DrugBank | Drug | 17,430 |

#### 2.1.2 Relation Database Introduction and Collection

This study integrates not only entity datasets but also a comprehensive range of relational datasets, facilitating the exploration of various biological and chemical interactions (see **Figure 2**). These relational datasets capture complex relationships between genes, transcripts, proteins, drugs, diseases, phenotypes, pathways, metabolites, and microbiota, supporting advanced analyses in precision health (check data overall details in **Table 2** and **Table S2** for data collection details in supplementary section).

**Table 2.** General Information about Relation Databases.

| Database Names | Full Names | From | To | Edge Types | Number of Rows |
| --- | --- | --- | --- | --- | --- |
| Ensembl<sup>40</sup> | Ensembl | Gene | Transcript | Gene-Transcript | 412,034 |
|  |  | Transcript | Protein | Transcript-Protein | 412,034 |
| RefSeq<sup>44</sup> | Reference Sequence Database | Gene | Transcript | Gene-Transcript | 33,421 |
|  |  | Transcript | Protein | Transcript-Protein | 30,207 |
| UniProt<sup>46</sup> | Universal Protein Database | Transcript | Protein | Transcript-Protein | 20,417 |
|  |  | Protein | Disease | Protein-Disease | 20,417 |
| BioGrid<sup>71,72</sup> | Biological General Repository for Interaction Datasets | Protein | Protein | Protein-Protein | 942,241 |
| STRING<sup>73,74</sup> | Search Tool for the Retrieval of Interacting Genes/Proteins | Protein | Protein | Protein-Protein | 13,715,404 |
| KEGG<sup>48</sup> | Kyoto Encyclopedia of Genes and Genomes | Protein | Protein | Protein-Protein | 52,973 |
|  |  | Protein | Pathway | Protein-Pathway | 21,051 |
|  |  | Metabolite | Pathway | Metabolite-Pathway | 19,125 |
|  |  | Pathway | Protein | Pathway- Protein | 24,475 |
|  |  | Pathway | Drug | Pathway-Drug | 2,334 |
|  |  | Drug | Pathway | Drug-Pathway | 3,922 |
| HPO<sup>59</sup> | Human Phenotype Ontology | Protein | Phenotype | Gene-Phenotype | 316,718 |
|  |  | Protein | Disease | Gene-Disease | 15,593 |
|  |  | Phenotype | Phenotype | Phenotype-Phenotype | 19,533 |
|  |  | Phenotype | Disease | Phenotype-Disease | 271,776 |
|  |  | Disease | Phenotype | Disease-Phenotype | 271,776 |
| DisGeNet<sup>75</sup> | DisGeNet | Protein | Disease | Protein-Disease | 91,484 |
| DISEASES<sup>76</sup> | DISEASES | Protein | Disease | Protein-Disease | 96,763 |
| HMDB<sup>52</sup> | Human Metabolome Database | Metabolite | Protein | Metabolite-Protein | 863,759 |
|  |  | Metabolite | Disease | Metabolite-Disease | 27,670 |
|  |  | Drug | Metabolite | Drug-Metabolite | 3,258 |
| STITCH | STITCH ('search tool for interactions of chemicals') | Metabolite | Protein | Metabolite-Protein | 15,473,939 |
| MetaNetX<sup>77</sup> | MetaNetX | Metabolite | Metabolite | Metabolite-Metabolite | 11,723 |
| DisBiome<sup>78</sup> | DisBiome | Microbiota | Disease | Microbiota-Disease | 10,866 |
| HMDAD | Human Microbe-Disease Association Database | Microbiota | Disease | Microbiota-Disease | 483 |
| MDAD<sup>79</sup> | Microbe-Drug Association Database | Microbiota | Drug | Microbiota-Drug | 5,055 |
|  |  | Drug | Microbiota | Drug-Microbiota | 5,055 |
| PharmacoMicrobiomics<sup>80</sup> | PharmacoMicrobiomics | Microbiota | Drug | Microbiota-Drug | 69 |
|  |  | Drug | Microbiota | Drug-Microbiota | 69 |
| CTD<sup>58</sup> | The Comparative Toxicogenomics Database | Pathway | Exposure | Pathway-Exposure | 1,624,470 |
|  |  | Exposure | Gene | Exposure-Gene | 2,892,325 |
|  |  | Exposure | Pathway | Exposure-Pathway | 1,624,470 |
|  |  | Exposure | Disease | Exposure-Disease | 9,329,083 |
| DO<sup>63</sup> | Disease Ontology | Disease | Disease | Disease-Disease | 14,339 |
| DrugBank<sup>70</sup> | DrugBank | Drug | Protein | Drug-Protein | 26,245 |
|  |  | Drug | Drug | Drug-Drug | 2,855,848 |
| BindingDB<sup>81</sup> | Binding Database | Drug | Protein | Drug-Protein | 1,610,889 |
| DrugCentral<sup>82</sup> | DrugCentral | Drug | Protein | Drug-Protein | 14,301 |
|  |  | Drug | Disease | Drug-Disease | 42,307 |
| SIDER<sup>83</sup> | Side Effect Resource | Drug | Phenotype | Drug-Phenotype | 309,849 |

The last column represents the total number of rows in the original dataset from each database.

### 2.2 Harmonizing Resources

As shown in **Figure 1**, the integrated biomedical knowledge graph system, ***BioMedGraphica***, has been proposed. With collected datasets from various sources, the knowledge graph will integrate 11 different types of entities from 30 databases into a universal knowledge database. And the promoter entities directly copied the entities from gene entities, since the ***BioMedGraphica*** defaultly set the promoters have the impacts on the genes. In addition, the relationship between those entities were included by harmonizing the 22 relational databases with 30 edge types. The details of merging and harmonizing can check following descriptions.

**Figure 1.**
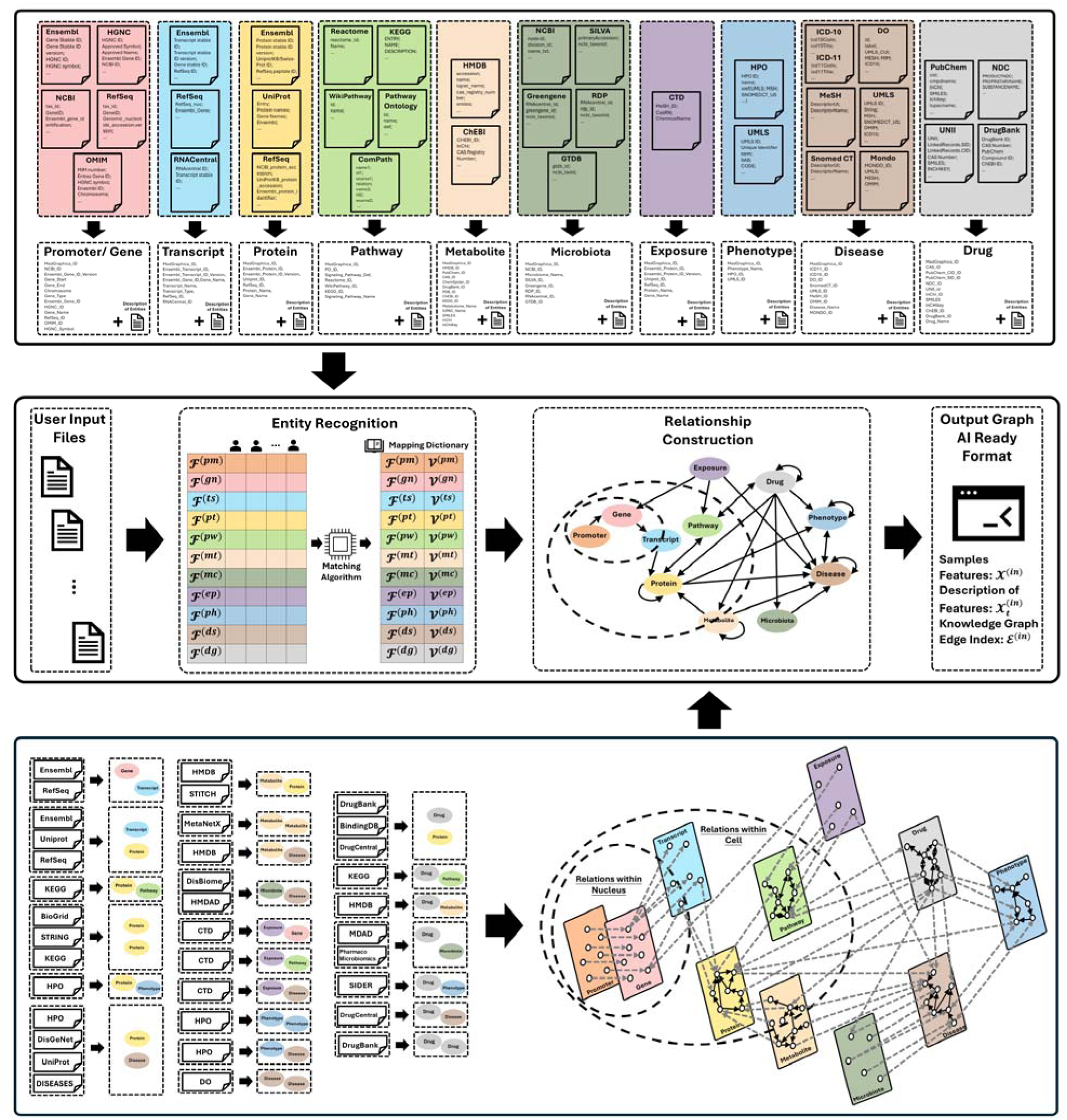
Overview of ***BioMedGraphica.*** Upper panel shows integration of the entities from various databases. Lower panel demonstrates the relation harmonization process and constructed knowledge graph. Mid panel displays the general procedures of BioMedGraphica, with entity recognition and relationship construction based on the user input files, outputting the graph AI ready format files.

**Figure 2.**
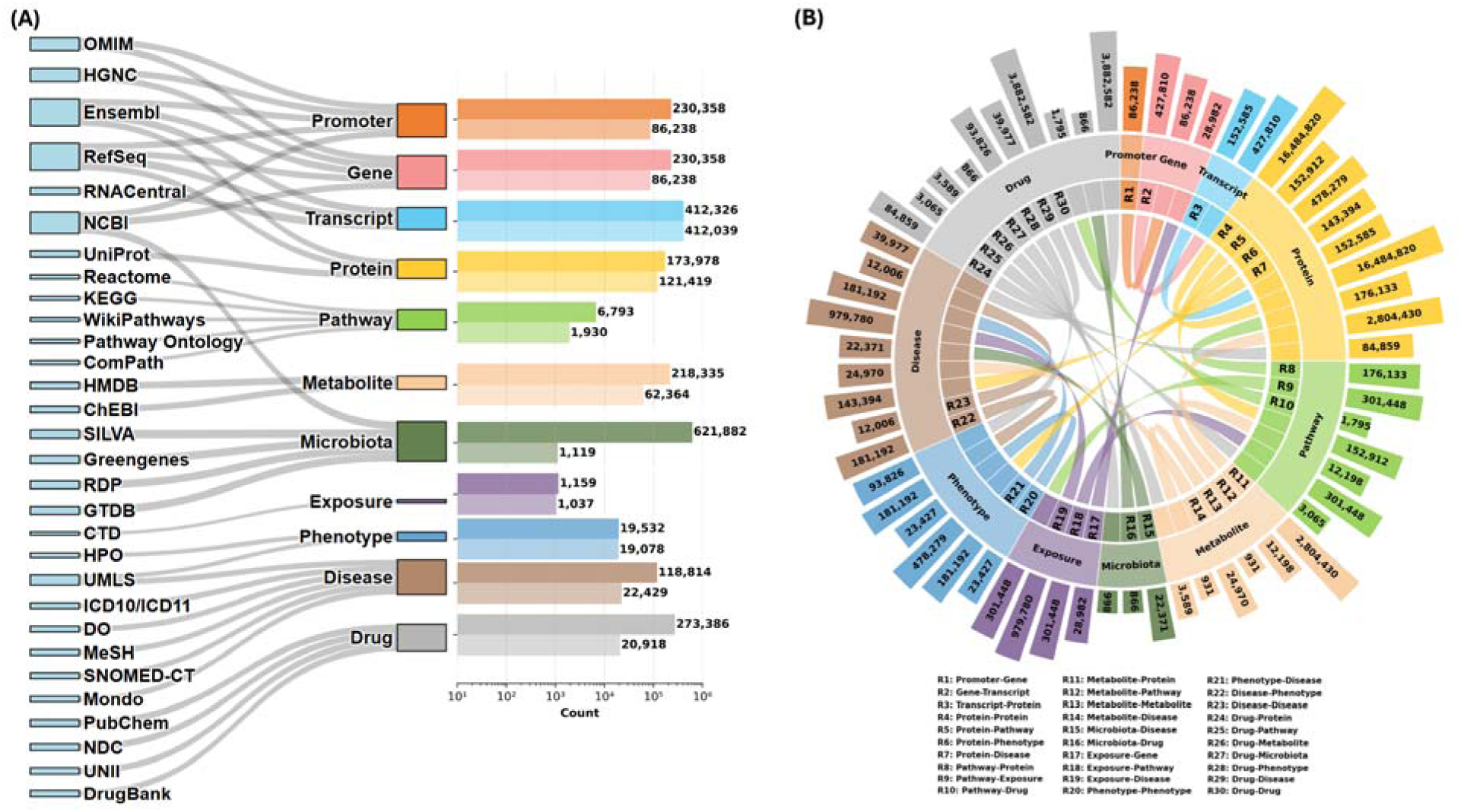
Overview of integrated biomedical entities and their relations in ***BioMedGraphica***. **(A)** Data sources and entity distributions. The left panel shows the data sources (e.g., OMIM, HGNC, Ensembl, UniProt, KEGG, SILVA, DrugBank, etc.) used to define and harmonize 11 biomedical entity types: Promoter, Gene, Transcript, Protein, Pathway, Metabolite, Microbiota, Exposure, Phenotype, Disease, and Drug. The right panel presents the logarithmic-scaled bar plot showing the total number of unique entities in full BioMedGraphica (upper bar) and BioMedGraphica-Conn (lower opacity bar). **(B)** Circular chord diagram of pairwise relationships between biomedical entities. Each colored segment represents a specific entity type, and outer arcs quantify the total number of cross-entity edges for each type. The inner chords indicate the direction and volume of entity-to-entity relationships (e.g., Gene–Transcript, Protein– Pathway, Microbiota–Disease). Each relationship type is labeled (e.g., R1: Promoter–Gene, R2: Gene–Transcript, R10: Pathway–Drug, R23: Disease–Drug and total edge counts for selected relationships are annotated.)

#### 2.2.1 Entity Integration

To construct a unified biomedical knowledge base, comprehensive entity integration was performed across multiple biological domains. For gene entities, datasets from Ensembl, HGNC, and NCBI were first merged based on Ensembl IDs, followed by incorporation of RefSeq and OMIM data using NCBI Gene IDs as the primary unifying identifier. Transcript entities were integrated by adopting the Ensembl Transcript Stable ID as the standard reference, with descriptions retrieved through the Ensembl BioMart API. Protein entities were merged by aligning Ensembl and UniProt records via Protein Stable ID Versions, subsequently incorporating RefSeq mappings, with annotations obtained from UniProt. Pathway entities were integrated using Pathway Ontology (PO) as the foundational framework, supplemented by KEGG, Reactome, and WikiPathway datasets through equivalent mappings facilitated by ComPath. For metabolite entities, ChEBI IDs were used initially for alignment, with HMDB IDs ultimately established as the minimal granularity unit. Microbiota data were harmonized using NCBI Taxon IDs to ensure consistency across datasets. Exposure entities were unified based on CAS numbers, leveraging their broad availability across relevant databases. Phenotype integration was initiated from HPO terms, with systematic cleaning of labels to remove generic descriptors and consolidation based on refined labels linked to HPO identifiers. Disease entities were integrated through a multi-step mapping strategy involving UMLS, MeSH, SNOMED-CT, ICD-10, ICD-11, Disease Ontology, and Mondo, with UMLS IDs designated as the minimal unit of granularity. Finally, drug entities were merged by aligning NDC and UNII datasets using substance names, followed by incorporation of PubChem, CAS, ChEBI, and DrugBank information through mapped identifiers, establishing CAS numbers as the primary reference. Throughout the integration process, database merging was anchored by bolded columns in the supplementary tables, ensuring the uniqueness of key identifiers, with detailed workflows and results documented in Supplementary Figures **S1**–**S10** and Tables **S3**–**S12**.

#### 2.2.2 Relation Integration

The construction of edges utilized data from **22** distinct databases, mapping raw database IDs to their corresponding BioMedGraphica IDs to form relationships. A notable challenge arose from one-to-many mappings, where a single database ID, such as 614807 (OMIM ID), corresponds to multiple BioMedGraphica IDs (BMG_DS065861 and BMG_DS080589), due to the OMIM databases having one-to-many relationships with other databases. Aside from this, all relationships were directional and presented in a From-To format. To address bidirectional relationships, two distinct methodologies were employed. The first involved reversing the direction of the relationship. For instance, while protein-protein interactions are intrinsically bidirectional, the original dataset lacked explicit directionality. To resolve this, a reversed copy of the data was generated, merged with the original dataset, and duplicates were subsequently eliminated. The second approach entailed establishing new relationships where reversal was inappropriate. For example, in disease-phenotype associations, reversing the data alone was insufficient; instead, a complementary phenotype-to-disease relationship was created to accurately represent the connection. The edge structure was meticulously designed to conform to a one-to-one mapping framework, ensuring that each instance of a database ID mapping to multiple BioMedGraphica IDs resulted in the generation of distinct edges. This strategy significantly amplified the total number of edges, exceeding a straightforward summation of inter-database relationships due to the one-to-many nature of the mappings, see **Table 3** for details.

**Table 3.** Harmonized Relations Information.

| Interaction Type | Database | Initial Edge Number | Matching |  | Total |
| --- | --- | --- | --- | --- | --- |
|  |  |  | Unique | Total |  |
| Gene-Transcript | Ensembl | 412,034 | 412,034 | 427,393 | 427,810 |
|  | RefSeq | 33,421 | 6,625 | 7,001 |  |
| Transcript-Protein | Ensembl | 412,034 | 123,845 | 123,858 | 152,585 |
|  | Uniprot | 50,765 | 50,765 | 50,765 |  |
|  | RefSeq | 30,207 | 6,868 | 35,575 |  |
| Protein-Protein | BioGrid | 1,690,122 | 1,689,574 | 1,689,574 | 16,484,820 |
|  | STRING | 13,715,404 | 13,285,010 | 13,193,859 |  |
|  | KEGG | 52,973 | 51,971 | 1,967,514 |  |
| Protein-Pathway | KEGG | 21,051 | 20,867 | 152,912 | 152,912 |
| Protein-Phenotype | HPO | 259,118 | 255,171 | 478,279 | 478,279 |
| Protein-Disease | UniProt | 4,821 | 4,784 | 5,573 | 143,394 |
|  | DISEASES | 69,896 | 11,108 | 12,387 |  |
|  | HPO | 7,482 | 7,184 | 15,444 |  |
|  | DisGeNet | 91,484 | 70,968 | 129,750 |  |
| Pathway-Protein | KEGG | 24,475 | 24,368 | 176,133 | 176,133 |
| Pathway-Drug | KEGG | 2,334 | 1,331 | 1,795 | 1,795 |
| Pathway-Exposure | CTD | 1,624,470 | 305,427 | 301,448 | 301,448 |
| Metabolite-Protein | HMDB | 863,759 | 849,980 | 849,993 | 2,804,430 |
|  | STITCH | 74,576,305 | 1,973,337 | 1,975,413 |  |
| Metabolite-Pathway | KEGG | 19,622 | 7,208 | 12,198 | 12,198 |
| Metabolite-Metabolite | MetaNetX | 23,711 | 886 | 931 | 931 |
| Metabolite-Disease | HMDB | 24,755 | 24,669 | 24,970 | 24,970 |
| Microbiota-Disease | DisBiome | 8,438 | 3,521 | 4,362 | 22,371 |
|  | HMDAD | 450 | 21,663 | 18,014 |  |
| Microbiota-Drug | MDAD | 5,055 | 2,078 | 816 | 866 |
|  | PharmacoMicrobiomics | 69 | 68 | 67 |  |
| Exposure-Gene | CTD | 42,261 | 28,982 | 323,906 | 323,906 |
| Exposure-Pathway | CTD | 1,624,470 | 305,427 | 301,448 | 301,448 |
| Exposure-Disease | CTD | 9,329,083 | 679,195 | 979,780 | 979,780 |
| Phenotype-Phenotype | HPO | 23,461 | 23,427 | 23,427 | 23,427 |
| Phenotype-Disease | HPO | 155,989 | 155,063 | 181,192 | 181,192 |
| Disease-Phenotype | HPO | 155,989 | 155,063 | 181,192 | 181,192 |
| Disease-Disease | DO | 11,836 | 9,747 | 12,006 | 12,006 |
| Drug-Protein | DrugBank | 25,670 | 20,865 | 23,100 | 84,859 |
|  | BindingDB | 1,161,440 | 58,858 | 58,858 |  |
|  | DrugCentral | 14,301 | 13,427 | 15,107 |  |
| Drug-Pathway | KEGG | 3,922 | 2,380 | 6,235 | 6,235 |
| Drug-Metabolite | HMDB | 3,258 | 3,171 | 3,065 | 3,065 |
| Drug-Microbiota | MDAD | 5,055 | 2,078 | 805 | 866 |
|  | PharmacoMicrobiomics | 69 | 68 | 67 |  |
| Drug-Phenotype | SIDER | 152,759 | 91,692 | 93,826 | 93,826 |
| Drug-Disease | DrugCentral | 50,011 | 35,940 | 39,977 | 39,977 |
| Drug-Drug | DrugBank | 2,855,848 | 2,845,794 | 3,882,582 | 3,882,582 |

## 3 Results

### 3.1 BioMedGraphica: an integrative textual biomedical prior knowledge graph

For entity integration, the promoter entity was added to the entity by copying gene entity due to the intuitively setting each gene was influenced by a corresponding promoter. Therefore, the database for ***BioMedGraphica*** includes **11** entity types and **30** edge types, contains **2,306,921** entities and **27,232,091** relations, composing the knowledge graph *g= (V, ε)* and **834,809** entities and **27,087,971** relations, composing the connected knowledge graph *g_c_* = (*V*_c_, ε_c_) (check number of each entity and edge type in **Table 4** and **Table 5**). Beyond structural knowledge integration, each entity is further enriched with comprehensive textual annotations, denoted as *T* and *T_c_* for the full graph f and its connected component *g_c_*, respectively. These include nomenclature records harmonized across multiple biomedical resources, capturing alternative identifiers, synonyms, and cross-references that resolve inconsistencies across databases. In addition, descriptive metadata provides functional insights, mechanistic roles, and biological contexts, such as molecular activities, pathway involvement, disease associations, and therapeutic relevance (details are documented in Supplementary Figures **S1**–**S10**). By combining nomenclature harmonization with descriptive annotations, the textual knowledge graph not only ensures consistent entity recognition but also provides a semantically rich layer of biomedical knowledge that facilitates graph–text fusion, interpretability, and downstream AI applications.

**Table 4.** Summarized Entity Information.

| Entity Type | Math Annotation | BMG Count | BMG Percentage | BMGC Count | BMGC Percentage | BMGC in BMG |
| --- | --- | --- | --- | --- | --- | --- |
| Promoter | $\mathcal{V}^{(pm)}$ | 230,358 | 9.9855% | 86,238 | 10.3303% | 37.4365% |
| Gene | $\mathcal{V}^{(gn)}$ | 230,358 | 9.9855% | 86,238 | 10.3303% | 37.4365% |
| Transcript | $\mathcal{V}^{(ts)}$ | 412,326 | 17.8734% | 412,039 | 49.3573% | 99.9304% |
| Protein | $\mathcal{V}^{(pt)}$ | 173,978 | 7.5416% | 121,419 | 14.5445% | 69.7899% |
| Pathway | $\mathcal{V}^{(pw)}$ | 6,793 | 0.2945% | 1,930 | 0.2312% | 28.4116% |
| Metabolite | $\mathcal{V}^{(mt)}$ | 218,335 | 9.4643% | 62,364 | 7.4705% | 28.5634% |
| Microbiota | $\mathcal{V}^{(mc)}$ | 621,882 | 26.9572% | 1,119 | 0.1340% | 0.1799% |
| Exposure | $\mathcal{V}^{(ep)}$ | 1,159 | 0.0502% | 1,037 | 0.1242% | 89.4737% |
| Phenotype | $\mathcal{V}^{(ph)}$ | 19,532 | 0.8467% | 19,078 | 2.2853% | 97.6756% |
| Disease | $\mathcal{V}^{(ds)}$ | 118,814 | 5.1503% | 22,429 | 2.6867% | 18.8774% |
| Drug | $\mathcal{V}^{(dg)}$ | 273,386 | 11.8507% | 20,918 | 2.5057% | 7.6515% |
| Total | $\mathcal{V}$ | 2,306,921 | 100% | 834,809 | 100% | 36.1872% |

\* BMG shorts for *BioMedGraphica*, and BMGC shorts for *BioMedGraphica-Conn*, which stands for the connected knowledge graph by removing the isolated nodes in *BioMedGraphica*.

**Table 5.** Summarized Information of Relation Types.

| Relation Type | Math Annotation | Count | Percentage |
| --- | --- | --- | --- |
| Promoter-Gene | $\mathcal{E}^{(pm-gn)}$ | 230,358 / 86,238 | 0.8459% / 0.3184% |
| Gene-Transcript | $\mathcal{E}^{(gn-ts)}$ | 427,810 | 1.5710% |
| Transcript-Protein | $\mathcal{E}^{(ts-pt)}$ | 152,585 | 0.5603% |
| Protein-Protein | $\mathcal{E}^{(pt-pt)}$ | 16,484,820 | 60.5345% |
| Protein-Pathway | $\mathcal{E}^{(pt-pw)}$ | 152,912 | 0.5615% |
| Protein-Phenotype | $\mathcal{E}^{(pt-ph)}$ | 478,279 | 1.7563% |
| Protein-Disease | $\mathcal{E}^{(pt-ds)}$ | 143,394 | 0.5266% |
| Pathway- Protein | $\mathcal{E}^{(pw-pt)}$ | 176,133 | 0.6468% |
| Pathway-Exposure | $\mathcal{E}^{(pw-ep)}$ | 301,448 | 1.1070% |
| Pathway-Drug | $\mathcal{E}^{(pw-dg)}$ | 1,795 | 0.0066% |
| Metabolite-Protein | $\mathcal{E}^{(mt-pt)}$ | 2,804,430 | 10.2982% |
| Metabolite-Pathway | $\mathcal{E}^{(mt-pw)}$ | 12,198 | 0.0447% |
| Metabolite-Metabolite | $\mathcal{E}^{(mt-mt)}$ | 931 | 0.0034% |
| Metabolite-Disease | $\mathcal{E}^{(mt-ds)}$ | 24,970 | 0.0916% |
| Microbiota-Disease | $\mathcal{E}^{(mc-ds)}$ | 22,371 | 0.0821% |
| Microbiota-Drug | $\mathcal{E}^{(mc-dg)}$ | 866 | 0.0032% |
| Exposure-Gene | $\mathcal{E}^{(ep-gn)}$ | 28,982 | 0.1064% |
| Exposure-Pathway | $\mathcal{E}^{(ep-pw)}$ | 301,448 | 1.1070% |
| Exposure-Disease | $\mathcal{E}^{(ep-ds)}$ | 979,780 | 3.5979% |
| Phenotype-Phenotype | $\mathcal{E}^{(ph-ph)}$ | 23,427 | 0.0860% |
| Phenotype-Disease | $\mathcal{E}^{(ph-ds)}$ | 181,192 | 0.6654% |
| Disease-Phenotype | $\mathcal{E}^{(ds-ph)}$ | 181,192 | 0.6654% |
| Disease-Disease | $\mathcal{E}^{(ds-ds)}$ | 12,006 | 0.0441% |
| Drug-Protein | $\mathcal{E}^{(dg-pt)}$ | 84,859 | 0.3116% |
| Drug-Pathway | $\mathcal{E}^{(dg-pw)}$ | 3,065 | 0.0113% |
| Drug-Metabolite | $\mathcal{E}^{(dg-mt)}$ | 3,589 | 0.0132% |
| Drug-Microbiota | $\mathcal{E}^{(dg-mc)}$ | 866 | 0.0032% |
| Drug-Phenotype | $\mathcal{E}^{(dg-ph)}$ | 93,826 | 0.3445% |
| Drug-Disease | $\mathcal{E}^{(dg-ds)}$ | 39,977 | 0.1468% |
| Drug-Drug | $\mathcal{E}^{(dg-dg)}$ | 3,882,582 | 14.2574% |
| Total | $\mathcal{E}$ | 27,232,091 / 27,087,971 | 100% |

\*For promoter-gene relation, the column count demonstrates the number of relations in BMG and BMGC respectively, due to that promoter-gene relation are virtual relation generated automatically.

### 3.2 A tool for Textual-Numeric Graph (TNG) data generation

BioMedGraphica is a unified software platform designed to integrate heterogeneous biomedical datasets with a structured knowledge graph and affiliated textual annotations, enabling coherent representation for both graph foundation models and agentic AI systems. By bridging data ranging from omic data to clinical data with curated graph-based knowledge, the tool provides a scalable framework for constructing TNG subsets tailored to user inputs. As shown in **Figure 3**, user can input the files into the software, which are 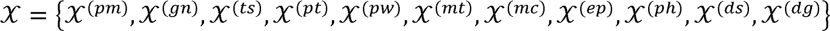, where 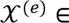 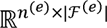 and e denotes one of the 11 entity types mentioned above, *n_e_* stands for the number of samples, ‘.*F*(e)represents features set of the entity type e. Once the files are imported, sample sizes across entity types are aligned to *n^(in)^*, defined as the intersection of all inputs. This alignment yields a unified matrix 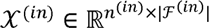, and 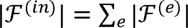, which can be viewed as a consolidated representation of the user-provided data. To ensure semantic coherence and enable effective inference, BioMedGraphica employs the BioMedGraphica-Conn, *g_c_*, as a solid knowledge graph foundation, which removes isolated nodes and retains only the largest connected subgraph. This strategy is consistent with graph theory principles that emphasize the importance of connectivity for traversal and reasoning and aligns with prior work showing that connected knowledge graphs enhance AI-driven interpretability and mechanistic discovery^84–86^. Hence, by matching the features ’.*F^(in)^* with entities V existing in knowledge graph *g_c_*, the entities will be formed with *V^(in)^* and mapping function *M: ’.F^(in)^ ➔ V^(in)^*, which is curated in python dictionary format.

**Figure 3.**
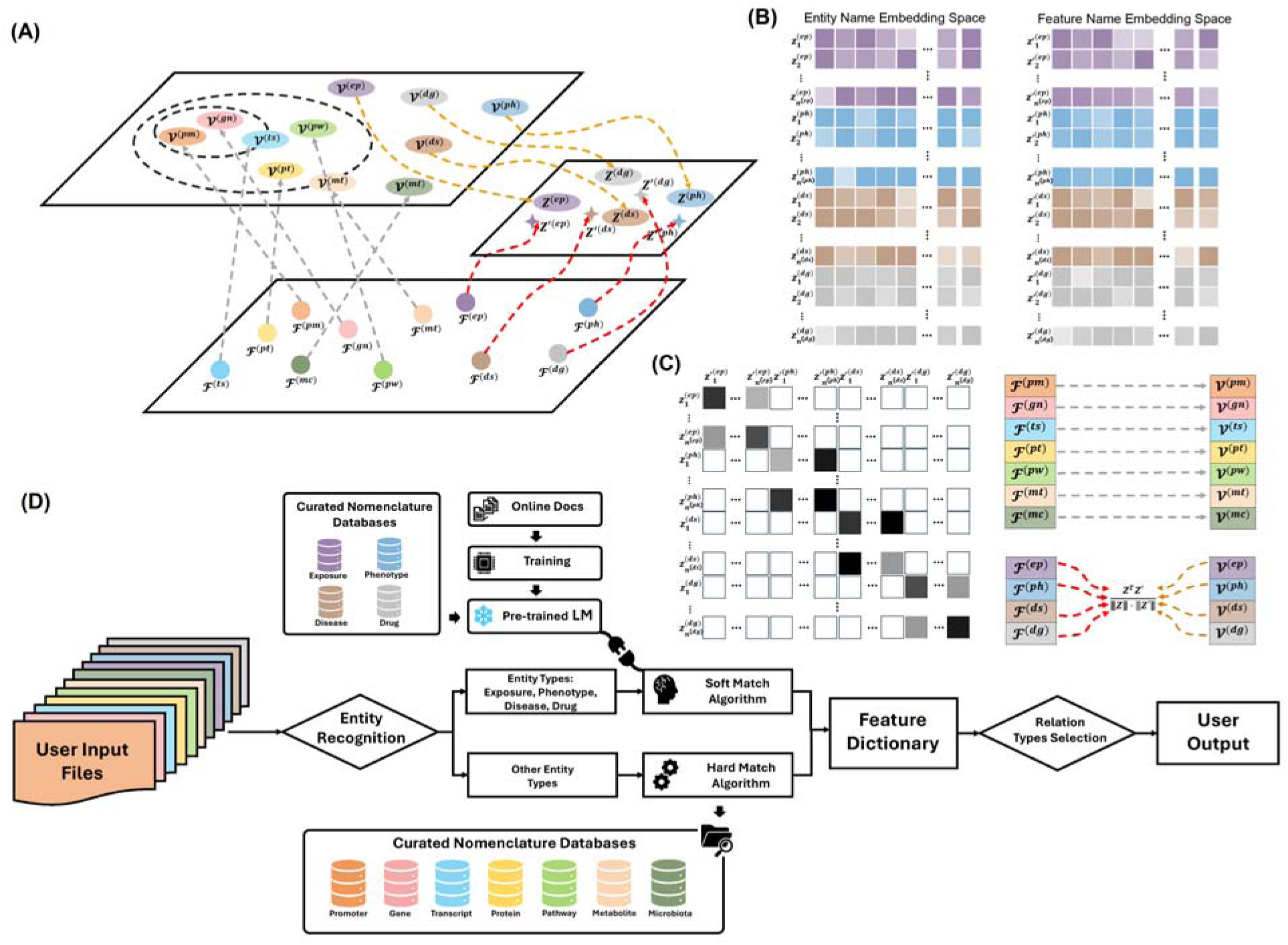
Pipeline of software ***BioMedGraphica***. **(A)** Entity matching. Two strategies are used: a hard match that retrieves entities via standardized identifiers from curated nomenclatures, and a soft match that embeds both entity names and user-provided feature names into a shared representation space with a pre-trained LM, selecting the highest-similarity candidate. **(B)** Schematic of embedding spaces for entity names and feature names, illustrating their vectorized representations by pretrained LM. **(C)** Cosine similarities yield a top *k* candidate set, and user confirmation finalizes a one-to-one mapping for each feature to generate mapping dictionary *D*. **(D)** BioMedGraphica pipeline begins with user input files (with sample integration). For standardized-ID entities—promoter, gene, transcript, protein, pathway, metabolite, microbiota—BioMedGraphica performs hard matching against curated nomenclatures. For free-text entities—exposure, phenotype, disease, drug—it applies soft matching with a pre-trained LM for semantic alignment. Matched items are consolidated into a feature dictionary with contextual attributes; relation selection and auto-completion then produce structured outputs (TNG subsets). The workflow combines curated-ID precision with LM-based semantics to generate AI-ready biomedical graphs enriched with textual annotations.

In addition, users may incorporate virtual entities that function as essential intermediates in biological processes (e.g., transcripts in the gene–transcript–protein chain), thereby producing a refined entity set *V^(sub)^*. Users may also specify the relation types to include, yielding a subgraph 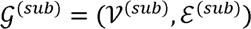. Together with the matched and auto-completed (virtual) entities, a descriptive feature matrix *T*^(sub)^ 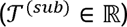, is generated, representing the associated textual annotations. This process results in an aligned feature space of size|*V*^(sub)^|, forming a new feature matrix 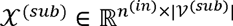. Finally, the complete TNG user-specific subset is produced as 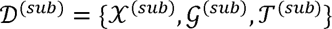. The subsequent section elaborates on the procedures for entity matching and relation construction.

#### 3.2.1 Entity Recognition

The entities from the input files will be recognized by either hard match algorithm or soft match algorithm. For most of entity types including promoter, gene, transcript, protein, pathway, metabolite and microbiota, having uniformed IDs to be served as the matching symbols, they can be applied with the hard match algorithm by fitting them into the IDs collected from various resources included in the BioMedGraphica. However, for the entity types with flexibility to name them with self-definition, including exposure, phenotype, disease and drug, they should be applied with the soft match algorithm.

When matching the features input by users to the existing entities in the BioMedGraphica knowledge base, the specialized designed algorithm using a pre-trained BioBERT model was leveraged for disease, phenotype, drug and exposure entities, which allows for the comparison of disease, phenotype, drug and exposure terms based on their semantic similarity for building the mapping dictionary *D*. Then, the similarity score will be calculated between a given query feature name, *f_name_* (*f* is the corresponded entity), and precomputed entity embeddings by scoring function S with

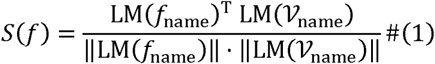

where 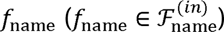 is the queried feature name from the unified user input file, 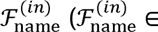 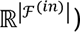 and 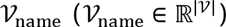 is the corresponding entity names of *F^(in)^* and *V*, and pre-trained BioBERT language model is denoted as LM. In detail, the model will process phenotype, drug and disease entities in ***BioMedGraphica*** by

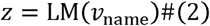

where *v*_name_ (*v*_name_ ∈ *V*_name_) is entity name and 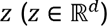 denotes the transformed embedding space for *f*_name_. Similarly, the queried feature name will be embedded by

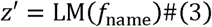

where 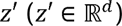 denotes the transformed embedding space for *f*_name_. Afterwards, the top *k* most similar entities will be extracted by

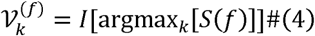

where argmax*_k_* can identify top *k* most similar entity names 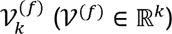 and l(•) is the one-to-one mapping function which will map the entity names to entities in ***BioMedGraphica***. In these top *k* most similar entity, the user will define only one entity, *V^(f)^*, to be matched for the queried feature name *f*_name_. For other entity types, the hard match method was leveraged to search for exact entity name for the queried feature name *f*_name_ with *V^(f)^*. With this, the dictionary function M will be generated.

#### 3.2.2 Relation / Knowledge Graph Construction

By extracting the corresponding entities *V^(in)^* of the input features ’.*F^(in)^* from the connected knowledge graph *<u>g_c_,</u>* users can select the edge types annotated in **Table 5** to construct the *ε^(in)^*. To ensure the connectivity of the constructed subgraph, we designed a shortest-path–based connectivity assessment and auto-completion strategy. Specifically, if certain downstream nodes are missing, they are labeled as candidate entities for supplementation and corresponding virtual node suggestions are generated. The core connection, defined as 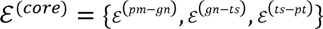, which serves as the backbone for connectivity analysis and the set of nodes is denoted as 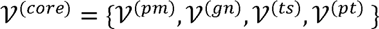.

To alleviate computational complexity, entity and relation types are generalized into coarse-grained categories within the BioMedGraphica abstract knowledge graph (Figure 1). This abstraction yields an abstract graph 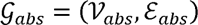, consisting of 11 nodes and 30 edges, each corresponding to the entity and relation types of the underlying concrete knowledge graph. Based on the abstract knowledge graph *g_abs_*, we constructed an undirected knowledge graph 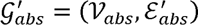 for algorithm development. Within abs abs this framework, the abstract core connection set is denoted as 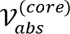 with its corresponding node set 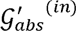, while the abstract input connection set and its associated nodes are represented as 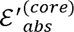 and 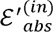, respectively, defining the input-specific graph as 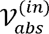. Based on the proposed strategy, we abs abs devised Algorithm 1 to implement the verification and supplementation process in the abstract knowledge graph. The procedure begins by identifying a core connection chain whenever input entities intersect with protein, gene, or transcript nodes. Non-core entities are then attached to this chain through direct edges where possible, with missing nodes along these paths recorded as candidates for supplementation. For pairs that cannot be directly connected or when no core node is present, shortest paths are computed under a hop constraint, and the minimal set of missing nodes is proposed. Connectivity is ultimately assessed across all entity pairs: if no supplementation is required, the subgraph is deemed connected; otherwise, candidate nodes are iteratively suggested until connectivity is achieved.

##### Algorithm 1

Connectivity Verification of Abstract Knowledge Graph 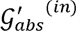

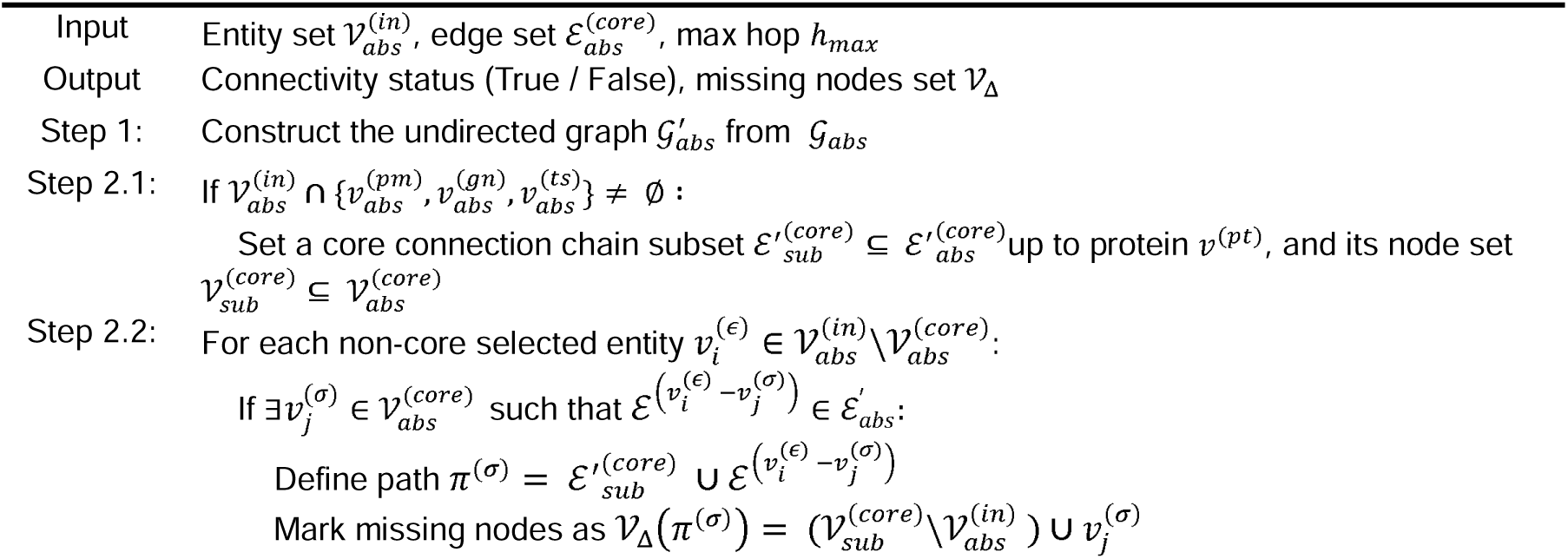
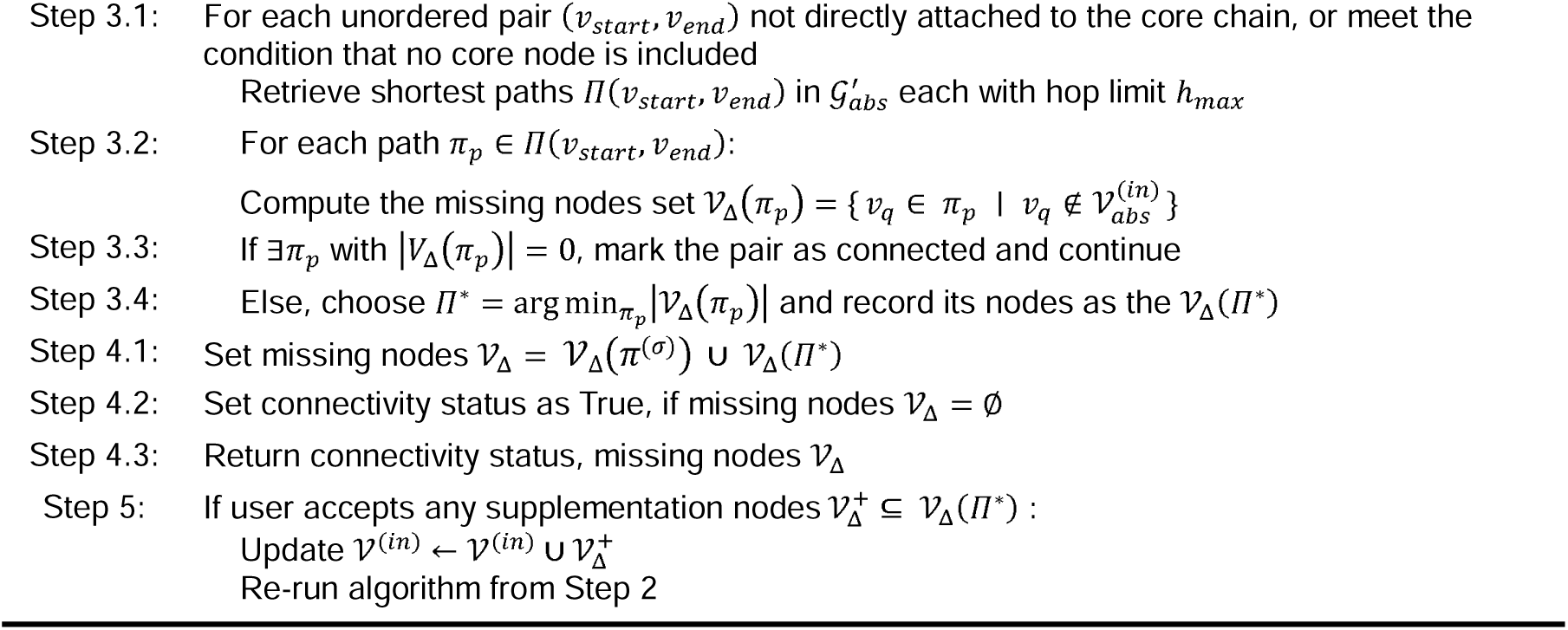

#### 3.2.3 Entities / Relations Matching Accuracy

The evaluation of entity and relation matching demonstrates a clear dichotomy between deterministic hard matching and probabilistic soft matching. Hard matching yields near-perfect accuracy for entities standardized under controlled vocabularies (e.g., promoter, gene, transcript, protein, etc.), confirming the robustness of BioMedGraphica’s curated nomenclature integration. In contrast, soft matching powered by pre-trained language models significantly enhances recognition for less standardized entities (e.g., phenotype, disease, exposure, drug), which enables automated semantic alignment and substantially improves efficiency over manual mapping, serving as a practical complement to hard matching for broader entity coverage. Nonetheless, current LLMs exhibit non-trivial error rates in multiple scenarios, including critical misassignments such as mapping *NC_000019.10* (RefSeq ID) to an incorrect Ensembl ID (*ENSG00000272512*). More concerning is their instability in relation extraction: for example, ChatGPT-5 erroneously linked the phenotype *Leukocytosis* to a misidentified drug with CAS number *106-60-5*, introducing fabricated relations inconsistent with biomedical ground truth. These results underscore both the strengths and limitations of generic language models, highlighting the need for BioMedGraphica’s hybrid strategy, which combines deterministic hard matches for structured entities with carefully constrained LM-driven soft matches, thereby ensuring reproducibility, accuracy, and trustworthiness in knowledge graph construction and downstream AI tasks.

#### 3.2.4 Graphical User Interface (GUI) Design

The GUI was developed to enhance accessibility and usability, enabling researchers without extensive programming expertise to efficiently construct TNG subsets through an intuitive, guided workflow. By lowering the technical barrier, the interface facilitates broader adoption of BioMedGraphica across interdisciplinary biomedical communities (see **Figure 4** for overview of BioMedGraphica GUI). To achieve this, the GUI implements a stepwise design that systematically guides users from data input to final output, ensuring both transparency and reproducibility in the processing pipeline. In detail, the GUI was developed to streamline the workflow of data input, recognition, filtering, and output generation. The interface begins with a user input module that supports both file upload and manual entry. Upon submission, the system performs automated data recognition and displays the inferred format in a preview pane for user validation. Users are then provided with options to refine the recognition type via dropdown menus or radio buttons (e.g., Entity Type A, Entity Type B), thereby enabling precise specification when necessary. Once confirmed, the workflow transitions to the entity-matching stage, where input records are aligned with the BioMedGraphica ID system. Only validated entities are retained for subsequent procedures. Users may further refine their datasets by selecting relational entities from curated databases (e.g., Relation Database 1, Relation Database 2), with the system reminding the inclusion of essential intermediate entities and relations to support automated completion. After filters are applied, the system generates the processed TNG, which can be downloaded or visualized in a structured format. Overall, the GUI ensures usability and accessibility by guiding users through each stage of processing with intuitive controls, real-time validation, and clear instructions. The application is publicly accessible at https://biomedgraphica.app.

**Figure 4.**
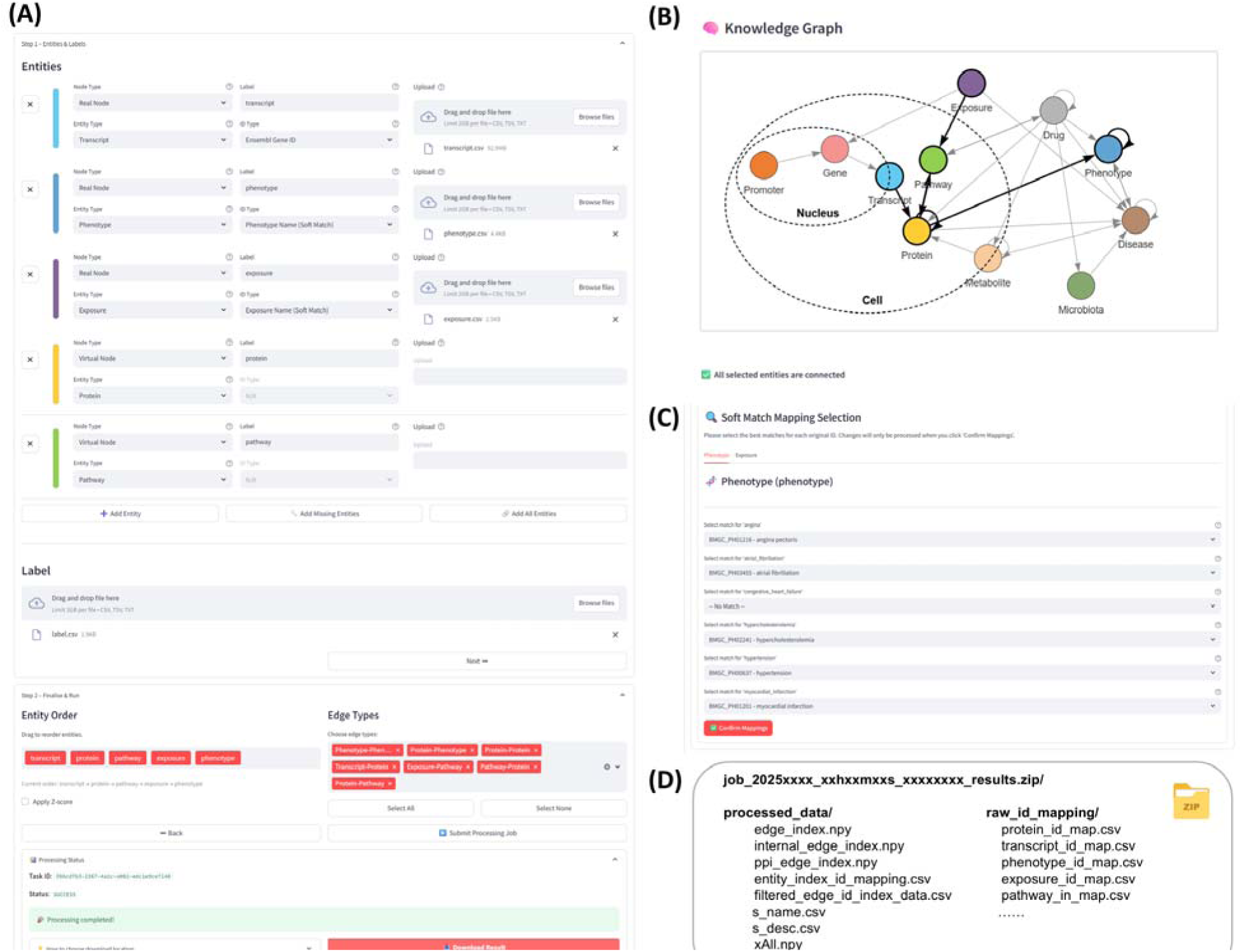
***BioMedGraphica*** GUI and usage demonstration of the Emory_Vascular dataset. **(A)** Overview of the web-based user interface, showing the two-step workflow for converting four input biomedical files into TNG formats. **(B)** The view of BioMedGraphica knowledge graph will highlight the selected entity types and relations based on the entity types integrated in Step 1. This panel also evaluates graph connectivity and marks missing entity types, which can be automatically completed via virtual entities if required. **(C)** The soft matching results interface, where candidate matches BioMedGraphica IDs are displayed, requiring user confirmation before proceeding. **(D)** Structure of the compressed output directory generated upon workflow completion, containing graph-ready feature matrices and entity-to-ID mapping files. For further details and instructions, please see the video demo from link: https://github.com/FuhaiLiAiLab/BioMedGraphica/blob/main/README.md

## 4 Data Access and Usage Demonstration

### 4.1 BioMedGraphica Knowledge Graph Data Access

We provide two levels of data accessibility to support diverse user needs. First, the raw datasets are made available with download links, accompanied by detailed processing instructions in supplementary section and open-source code hosted in our GitHub repository (https://github.com/FuhaiLiAiLab/BioMedGraphica/). These resources allow researchers to fully reproduce the data harmonization pipeline and customize entity and relation extraction according to their own requirements. Second, for users seeking ready-to-use resources, we also provide the processed and integrated BioMedGraphica knowledge graph through the Huggingface dataset link (https://huggingface.co/datasets/FuhaiLiAiLab/BioMedGraphica), enabling seamless adoption in downstream computational pipelines. To facilitate adoption across communities with varying technical expertise, we also released step-by-step tutorials in the above GitHub repository that guide users through the process of transforming raw data into harmonized entities and relations, ensuring semantic consistency across heterogeneous datasets. Upon completing the tutorial procedures, the resulting BioMedGraphica database functions as a comprehensive knowledge base that can directly interface with the BioMedGraphica software for knowledge graph construction and analysis. In parallel, a dedicated software tutorial is also provided, demonstrating how to initiate the platform, configure workflows, and generate tailored TNG subsets. Together, these resources ensure that BioMedGraphica is not only transparent and reproducible but also accessible and scalable, thereby empowering researchers across biomedical, computational, and translational domains to leverage the system effectively.

### 4.2 Usage Notes

#### Data Preparation: Entities and Labels

Prior to using the platform, users are required to prepare and standardize input files, consisting of feature files and a sample label file. For proper integration, all sample IDs across files should be harmonized to the same identifier system and listed in the first column of each file, thereby avoiding mismatches during downstream processing. Feature columns are expected to use either standardized database identifiers (e.g., Ensembl stable Gene IDs, HGNC symbols) or well-defined textual names (e.g., drug names, HPO terms), with any abbreviations expanded to facilitate reliable soft matching (see GitHub repository for detailed formatting guidelines). During upload, the platform performs real-time analysis of entity connectivity to ensure that the constructed knowledge graph remains fully connected. In cases where connectivity gaps are detected, the system automatically identifies missing components (see **Figure 4B**) and remind users to supplement them as virtual entities, based on connectivity checks against the core signaling graph, *E*(core), described in Section 3.2.2.

#### Configuration Finalization and Data Integration

Once all entities have been defined in Step 1, the workflow advances to Step 2, where the system automatically generates an entity ordering that reflects the canonical flow of biological signaling processes. This ordering, presented through the feature names assigned in the previous step, provides users with an intuitive representation of the signaling hierarchy. To accommodate specific experimental contexts or analytical preferences, users are given the option to manually refine this ordering through an interactive reordering panel (see **Figure 4A** step2). At this stage, additional configuration options are available, including the ability to enable z-score normalization of feature values and to specify which relation or edge types should be retained for downstream graph construction, thereby offering flexibility in tailoring the knowledge graph to the intended application.

After configuration is finalized, the system executes the data integration pipeline. This involves aligning common sample identifiers across all input files to ensure consistency, performing hard identifier matching for standardized nomenclatures, and applying embedding-based soft matching for entities expressed in natural-language terms. The soft matching process is implemented with user-in-the-loop confirmation, balancing automation with human oversight to improve reliability. The result of this pipeline is a set of AI-ready outputs, including graph-structured feature matrices and entity-to-identifier mapping files. These outputs are systematically packaged and made available for download through the platform interface, enabling immediate use in graph-based AI models and downstream biomedical analyses.

### 4.3 A Case Study: generating TNG using *BioMedGraphica*

To demonstrate the practical utility and technical robustness of BioMedGraphica, we make a case study from a real multi-omic dataset of Alzheimer’s disease (AD). About 6.5 million people are living with Alzheimer’s disease (AD) in the United States, and the estimated healthcare cost is about 321 billion, which will increase to 1 trillion by 2050^87^. There is no effective treatment for AD^88,89,90^, which is partially due to the unknown signaling pathways^91,92,93,94,95,96,97,98^ that lead to neurodegeneration, though more than 50 genes/loci have been associated with AD^91,99,100^. The data is accessible from the from AMP-AD portal (https://adknowledgeportal.synapse.org/Explore/Studies/DetailsPage/StudyData?Study=syn18909507) illustrates the complete workflow from entity configuration to graph-ready output (see **Figure 4**). In details, users are required to preprocess entity feature matrices so that the first column contains sample identifiers and the remaining columns store feature values. The sample label file should similarly use the first column for sample IDs and the second for class labels, with categorical labels pre-encoded into numeric values (e.g., via ordinal encoding). All input files must be supplied in CSV, TXT, or TSV format, and a consistent sample ID scheme must be applied across files.

To enable automatic recognition of entity types in Step 1 shown in **Figure 4**, it is recommended that filenames of processed feature files include the corresponding entity type keywords (e.g., promoter, transcript, protein). No specific naming is required for the sample label file. After uploading the entity files, the system automatically assigns entity types and labels, which users may adjust if necessary. Users then specify the identifier system (e.g., Ensembl gene ID, HGNC symbol) or textual names (e.g., drug names, HPO terms) associated with each file. The system subsequently evaluates knowledge graph connectivity using the shortest-path algorithm (shown in Section 3.2.2), highlights any missing nodes, and allows supplementation either automatically (via the *Add Missing Node* function) or manually (via user-defined virtual nodes). Connectivity status is updated in real time. Once the label file is uploaded and validated, users may proceed to Step 2 shown in **Figure 4**. In Step 2, BioMedGraphica automatically generates the entity file order and selects the relevant edge types based on predefined biological hierarchies and relation types. Users may review and optionally adjust these settings, including enabling z-score normalization of feature matrices. Upon final confirmation, the job is submitted to the server, and the system outputs a compressed archive containing graph-ready feature matrices and entity-to-ID mapping files. This concludes the data integration process.

## 5 Summary and Conclusion

Omic data analysis plays a crucial role in precision medicine for identifying novel disease-related targets and pathways. However, translating numeric and statistical omic analysis results into new scientific discoveries remains a major challenge, as human experts must manually review predicted targets, assess their statistical significance, and search extensive, inter-connected prior knowledge to generate hypotheses—a process that is subjective and not scalable. Large laboratories with rich resources can more easily test and validate discoveries, but smaller laboratories often struggle to generate the right testable hypotheses due to limited infrastructure and knowledge networks. This creates barriers to equitable discovery and slows the broader impact of omic data. Recently, large language models (LLMs) have begun to transform scientific discovery through their ability to interpret and reason with human-readable biomedical knowledge at scale. By integrating LLMs with multi-omic data analysis, it becomes possible to automate hypothesis generation, improve scalability, and accelerate precision medicine research.

Our unique contributions of this study are clear. First, we introduce a novel data format, TNG, integrating biomedical prior knowledge with quantitative data. In this format, entity names, descriptions, and biological functions are represented as textual information, while multi-omic profiles and other biomedical measurements are encoded as numeric values. To the best of our knowledge, this is the first systematic framework to propose TNG as a representation for integrating textual biomedical knowledge with biomedical data. Second, to facilitate the construction of TNGs, we developed ***BioMedGraphica***, an all-in-one platform that integrates 11 entity types and 30 relation types from 43 biomedical databases into a harmonized knowledge graph containing over 2.3 million entities and 27 million relations. Beyond its scope and harmonization, BioMedGraphica also provides a graphical user interface (GUI) that enables researchers to generate customized TNG datasets, bridging fragmented biomedical data with curated prior knowledge through an accessible and reproducible workflow. Third, the resulting TNGs are directly applicable for graph foundation models and large language model augmentation, providing both predictive power and mechanistic interpretability. Together, these advances establish BioMedGraphica as not only one of the most comprehensive biomedical knowledge graph resources to date but also as a cornerstone for the development of next-generation graph foundation models and agentic AI in biomedicine, paving the way for scalable, interpretable, and evidence-based discoveries that empower both large and small laboratories in precision medicine.

## Author contribution

FL conceptualized the project. Methodology and project design were developed by HZ, TX, WL, DH, GL, YC, MP, PP, FL. TX and HZ were responsible for software implementation. The manuscript was written by HZ, SL, TX, WL, FL. SL, HZ, WL, YD contributed to data collection and analysis.

## Supplementary Materials

### Section A. Details of Data Resources

#### A.1. Data Resources for Entities

##### Ensembl^40^

Ensembl is a widely used resource for genome annotation and provides access to a wide variety of genomic data across numerous species, with a strong focus on vertebrate genomes. The Ensembl project integrates gene, transcript, and protein data, offering detailed genomic features. This database was accessed through the BioMart API, which allows flexible retrieval of large datasets based on specific criteria. For gene entities, Gene Stable IDs were selected, including versioned identifiers that ensure traceability across different releases. Important genomic features such as gene start and end positions, biotypes, and chromosomal coordinates were extracted. To maintain consistency in gene nomenclature, the mapping relationships between Ensembl and the HUGO Gene Nomenclature Committee (HGNC) were preserved. This consistency is crucial for ensuring that gene annotations align across various databases. Additionally, transcript entities were obtained using Transcript Stable IDs and corresponding Gene IDs, while protein entities were extracted with Protein Stable IDs. Mapping relationships between Ensembl, UniProt, and RefSeq were also preserved to ensure accurate and cross-compatible dataset integration.

##### OMIM^41^ (Online Mendelian Inheritance in Man)

OMIM is an essential resource for understanding the genetic basis of human diseases and provides detailed information on gene-phenotype relationships. The database integrates clinical features with genetic data, offering insights into the hereditary nature of various conditions. Data from OMIM was retrieved to capture gene-related records, particularly focusing on mapping relationships between OMIM IDs and NCBI gene IDs. This facilitated the standardization of gene-related data across different resources. Furthermore, HGNC symbols were retained to align OMIM gene identifiers with other databases used in this study. Chromosomal information was also supplemented, which aids in genomic localization and contextual understanding of the data. Ensuring the uniqueness of gene records was a priority, and merging was performed based on gene IDs to guarantee that each entry in the dataset remained unique and free from redundancy.

##### HGNC^42^ (HUGO Gene Nomenclature Committee)

The HGNC is the authoritative resource for assigning unique symbols and names to human genes. As gene nomenclature can vary across different databases, the HGNC serves as a standard for the human genome, providing approved gene symbols and names. Data was accessed via BioMart to extract HGNC-approved gene information, including attributes such as HGNC ID, gene symbol, gene name, and chromosomal location. The inclusion of HGNC data ensures that gene-related information in the dataset is standardized and consistent with official naming conventions. Mapping relationships between HGNC, Ensembl, and NCBI IDs were retained to facilitate cross-referencing across these major databases. To ensure accuracy and prevent redundancy, the uniqueness of Ensembl IDs was verified during the merging process.

##### NCBI^43^ (National Center for Biotechnology Information) – Gene

The NCBI Gene database provides extensive information on genes and their functions, supporting a wide range of research in genetics, genomics, and bioinformatics. For this study, human gene data was extracted, retaining key attributes such as NCBI gene IDs, gene symbols, descriptive gene names, and chromosomal positions. The NCBI Gene database is an important resource for identifying gene sequences, gene structure, and gene functions, making it essential for the construction of a comprehensive gene dataset. Mapping relationships between NCBI gene IDs and Ensembl IDs were preserved to ensure consistency across datasets, facilitating the integration of data from different sources. For microbiome-related data, entries from the NCBI Taxonomy Database were also included. This database provides authoritative taxonomic classifications, focusing on bacterial taxa, and corresponding NCBI Taxon IDs were retained to ensure accurate classification and integration with other microbiome datasets.

##### NCBI - RefSeq^44^ (Reference Sequence Database)

RefSeq is a curated collection of publicly available nucleotide sequences and their corresponding protein translations, which provides a critical reference standard for the annotation of genes, transcripts, and proteins. RefSeq data was retrieved for both gene and transcript entities in this study, focusing on entries with the status of either “REVIEWED” or “MODEL” to ensure high data quality. Essential attributes such as gene ID, RefSeq ID, and chromosomal information were retained to provide accurate gene annotations. Additionally, the MANE project, which provides a set of transcript alignments between RefSeq and Ensembl, was utilized to ensure that transcript mapping between these databases was consistent and high-quality. Protein entities were also integrated, with mapping relationships between RefSeq, UniProt, and Ensembl retained to ensure cross-database compatibility. Uniqueness of the Ensembl IDs was verified throughout the data processing stages to ensure data integrity.

##### RNAcentral^45^

RNAcentral is a comprehensive resource for non-coding RNA sequences, integrating data from over 40 specialist databases. RNAcentral provides access to a wide variety of RNA sequence information, including microRNAs, tRNAs, and other functional RNA molecules that play critical roles in gene regulation. Human-specific RNAcentral IDs and corresponding Ensembl IDs were retrieved for this study, ensuring that non-coding RNA entities could be accurately integrated with gene and protein data from other databases. The uniqueness of each Ensembl ID was verified to ensure the integrity of the dataset and to avoid duplications during the integration process.

##### UniProt^46^ (Universal Protein Resource)

UniProt is a globally recognized repository of protein sequences and functional information. It provides detailed annotations on protein sequences, structure, function, and interactions. Data from UniProt was accessed via its API, and UniProt IDs, along with protein names and their corresponding Ensembl IDs, were retrieved. This enabled the integration of protein-specific data with the broader dataset, ensuring that protein information was accurately cross-referenced with gene and transcript data from Ensembl. The uniqueness of each Ensembl ID was verified during the data integration process to ensure consistency and to prevent errors in protein-related data.

##### Reactome^47^

Reactome is a curated knowledgebase of biological pathways, and it is a key resource for understanding the molecular mechanisms underlying cellular processes. Human-specific pathway data was extracted from Reactome for this study, enabling the integration of pathway-related information with gene and protein data. The inclusion of Reactome pathways facilitates research into functional genomics and systems biology, where pathway analysis is critical for understanding complex biological processes.

##### KEGG^48^ (Kyoto Encyclopedia of Genes and Genomes)

KEGG is a comprehensive resource for understanding high-level functions and utilities of biological systems, such as cells, organisms, and ecosystems. Human pathway data was retrieved using the bioservices package, with a focus on integrating KEGG pathways with other biological pathways from Reactome and WikiPathways. The inclusion of KEGG enables the dataset to support metabolic and signaling pathway analysis, providing valuable insights into cellular functions and disease mechanisms.

##### WikiPathways^49^

WikiPathways is an open, collaborative platform for the curation of biological pathways. Data from WikiPathways was converted to CSV format, and human-specific pathways were filtered for inclusion in this study. Mapping relationships between WikiPathways, KEGG, and Reactome were maintained to ensure consistent integration of pathway-related data. The inclusion of WikiPathways supports research into a wide variety of biological pathways, complementing the curated data from Reactome and KEGG.

##### Pathway Ontology^50^

Pathway Ontology provides a standardized framework for the classification of biological pathways and their relationships. Preprocessing of the OBO-formatted file enabled the extraction of PO IDs, and mapping relationships with KEGG and Reactome were preserved. This integration allows for comprehensive pathway analysis, ensuring that biological pathways from multiple sources can be consistently linked.

##### ComPath^51^

ComPath is a database that integrates pathway mapping relationships across KEGG, Reactome, and WikiPathways. All equivalent mappings were selected for this study, ensuring that pathway data from different sources could be cross-referenced. This comprehensive approach to pathway integration enables in-depth biological pathway analysis and facilitates the exploration of molecular mechanisms underlying diseases.

##### HMDB^52^ (Human Metabolome Database)

HMDB is the most comprehensive, freely accessible database of small molecule metabolites found in the human body. It provides extensive mapping relationships related to metabolomics data. For this study, HMDB data was parsed from XML files, retaining key attributes such as CAS number, SMILES, InChI, and mapping relationships with other databases. The inclusion of HMDB supports research into human metabolism, drug interactions, and disease mechanisms, enabling detailed metabolomics analysis.

##### ChEBI^53^ (Chemical Entities of Biological Interest)

ChEBI is a database focused on ’small’ chemical compounds and is used extensively for research in chemistry and biology. ChEBI provides manually annotated information about the structure, formula, and biological roles of chemical entities. In this study, ChEBI data was selected for drug entities, particularly those with a 3-star rating to ensure the highest data quality. Important attributes such as ChEBI ID, InChI, and the mapping relationship to CAS Registry Numbers were retained. In addition to drug entities, metabolome data from ChEBI was included, focusing on human metabolites. Mapping relationships with other databases, such as the Human Metabolome Database (HMDB), were preserved to enable cross-referencing of metabolite information.

##### SILVA^54^

SILVA is a high-quality, curated database of ribosomal RNA (rRNA) sequences, widely used for taxonomic classification of microbial communities. Data from both the small subunit (SSU) and large subunit (LSU) ribosomal RNA sequences were included, along with corresponding NCBI Taxon IDs. The SILVA database provides valuable insights into the composition of microbiomes, supporting research into microbial diversity and ecology.

##### Greengenes^55^

Greengenes is a database of 16S ribosomal RNA gene sequences used for the identification of microbial species. Data was sourced from RNAcentral, and RNAcentral IDs, Greengenes IDs, and NCBI Taxon IDs were retained to ensure consistent taxonomic classification of microbiome-related data. The inclusion of Greengenes allows for the accurate classification of bacterial species, supporting research into microbiomes and their impact on human health.

##### RDP^56^ (Ribosomal Database Project)

The Ribosomal Database Project (RDP) provides quality-controlled ribosomal RNA gene sequence data. Similar to Greengenes, RDP data was sourced via RNAcentral, and mapping relationships between RNAcentral IDs, RDP IDs, and NCBI Taxon IDs were preserved. This allows for the consistent classification of microbial entities, supporting microbiome research and analysis.

##### GTDB^57^ (Genome Taxonomy Database)

GTDB is a comprehensive resource for the classification of Archaea and Bacteria. Data from GTDB was retrieved for both archaeal and bacterial entities, with GTDB IDs and NCBI Taxon IDs retained to ensure accurate taxonomic classification. By verifying the uniqueness of NCBI Taxon IDs, the dataset provides reliable support for microbiome research, enabling the exploration of microbial diversity across various environments.

##### CTD^58^ (The Comparative Toxicogenomics Database)

CTD is a pivotal resource for integrating chemical, gene, disease, and exposure data, facilitating the study of toxicogenomics and environmental health. CTD serves as an entity-centric database where chemicals, genes, and diseases are interconnected through curated interaction data. For chemical entities, CTD uses standardized identifiers such as Chemical Abstracts Service (CAS) numbers to ensure consistent representation and integration with other chemical databases like PubChem.

##### HPO^59^ (Human Phenotype Ontology)

The Human Phenotype Ontology (HPO) provides a standardized vocabulary for phenotypic abnormalities encountered in human disease. Data was imported from the HPO database, version 2024-8-13, and relevant phenotype labels were extracted. After filtering and cleaning unwanted descriptive expressions, mapping relationships between HPO IDs were retained, ensuring that phenotypic data could be integrated with other disease and genomic datasets. This integration facilitates research into genotype-phenotype correlations, a key area in genetic and clinical research.

##### ICD^61,62^ (International Classification of Diseases)

The International Classification of Diseases (ICD), maintained by the World Health Organization, is the global standard for the coding and classification of diseases. Both ICD-10 and ICD-11 codes were included to ensure that the dataset could be used in various research and clinical contexts. Mapping relationships between ICD versions were retained, allowing for compatibility across different healthcare systems and facilitating research on disease epidemiology and outcomes.

##### Disease Ontology^63^ (DO)

The Disease Ontology (DO) provides a standardized ontology for the classification of human diseases. DO includes cross-references to other medical ontologies, such as UMLS, MeSH, and ICD-10, which were retained in this study to ensure consistent disease classification across databases. The inclusion of DO enabled the dataset to capture detailed and structured information on diseases, supporting research in medical informatics and bioinformatics.

##### MeSH^64^ (Medical Subject Headings)

MeSH is a comprehensive controlled vocabulary for the purpose of indexing journal articles and books in the life sciences. It is widely used in medical and biomedical research for categorizing diseases, drugs, and other entities. In this study, MeSH terms were retrieved from the MeSH XML files, focusing on records under the Diseases category. Mapping relationships with UMLS, ICD-10, and other disease ontologies were preserved to ensure consistency in terminology across datasets. This facilitated the integration of disease data and enabled the dataset to support detailed disease-related analyses.

##### UMLS^60^ (Unified Medical Language System)

The Unified Medical Language System (UMLS), developed by the National Library of Medicine (NLM), integrates multiple biomedical terminologies into a single framework. The Disease or Syndrome category of UMLS was selected for this study, with an emphasis on “Preferred” terms defined in English. Mapping relationships between UMLS, MeSH, SNOMED-CT, and ICD-10 were maintained to ensure accurate classification and cross-referencing of disease entities. The inclusion of UMLS data ensures that disease-related data can be consistently linked across multiple terminological systems, facilitating research in clinical informatics and biomedical research.

##### SNOMED-CT^65^ (Systematized Nomenclature of Medicine Clinical Terms)

SNOMED-CT is an international clinical terminology that is used to code the entire scope of human medical practice, including diseases, symptoms, diagnoses, and treatments. Data from the Snapshot version of SNOMED-CT was used to extract active entries from the “Disorder” category, preserving mapping relationships with ICD-10. This allowed for the integration of clinical disease information with other ontologies, enhancing the utility of the dataset for both clinical and research applications.

##### Mondo^66^

The Mondo Disease Ontology integrates multiple disease ontologies and databases, offering comprehensive cross-references to UMLS, MeSH, and other classification systems. Data from Mondo was included in this study, with a focus on preserving mapping relationships between Mondo IDs, UMLS, and MeSH. This integration enabled the consistent classification of disease entities, ensuring that disease-related data from different sources could be accurately linked.

##### PubChem^67^

PubChem is a large database of chemical molecules and their biological activities, maintained by the National Center for Biotechnology Information (NCBI). It is widely used for retrieving chemical information related to small molecules, including drugs, metabolites, and other compounds. For this study, data from the Drug and Medication Information and Pharmacology and Biochemistry categories within the PubChem compound catalog was extracted. Key chemical descriptors, such as InChI, SMILES, InChIKey, and IUPAC names, were selected to provide detailed chemical structure information. The PubChem CID (Compound Identifier) was used as a unique identifier to facilitate consistent cross-referencing of chemical compounds across datasets.

##### NDC^68^ (National Drug Code)

The National Drug Code (NDC) is a unique identifier for medications in the United States, maintained by the U.S. Food and Drug Administration (FDA). It is an essential resource for drug-related data integration. Data from the NDC was selected for inclusion, focusing on the NDC code and substance names, which correspond to the UNII (Unique Ingredient Identifier) code’s preferred term. By ensuring the uniqueness of each substance name, accurate integration with UNII data was facilitated, allowing for comprehensive drug-related analyses.

##### UNII^69^ (Unique Ingredient Identifier)

The Unique Ingredient Identifier (UNII) system, maintained by the FDA, assigns unique identifiers to chemical substances, including active ingredients in drugs. UNII data was sourced from both the PubChem website and the FDA, with mapping relationships between UNII codes, PubChem CIDs, and CAS numbers being preserved. Additionally, structural descriptors such as SMILES and InChIKeys were included, providing a detailed representation of the chemical substances. This ensures that UNII data can be integrated seamlessly with other chemical and pharmacological databases.

##### DrugBank^70^

DrugBank is a unique bioinformatics and cheminformatics resource that combines detailed drug data with comprehensive drug-target information. Data from DrugBank was included in this study to retain mapping relationships between DrugBank IDs and other chemical identifiers, such as PubChem CID, SID (Substance ID), and CAS numbers. DrugBank’s extensive annotation of drug targets and mechanisms of action made it a valuable resource for cross-referencing drugs with their molecular and clinical effects, enabling more in-depth pharmacological studies.

#### A.2 Data Resources for Relations

##### Ensembl^40^

Ensembl is a comprehensive genome browser and database that provides a wealth of information on gene sequences, annotations, and relationships across multiple species. It supports the analysis of gene-transcript interactions by linking genes to their corresponding transcripts. Ensembl also provides transcript-protein interaction, providing detailed annotations of how transcripts give rise to protein products. The dataset is essential for understanding gene structure, function, and the consequences of gene expression.

##### NCBI - RefSeq^44^ (Reference Sequence Database)

RefSeq is a well-curated collection of gene, transcript, and protein sequences, offering high-quality data for gene-transcript and transcript-protein relationships. It provides standardized and curated sequences that ensure consistency in gene annotations. RefSeq is crucial for researchers needing reliable reference sequences for various biological analyses, particularly in understanding the relationships between transcripts and their encoded proteins.

##### UniProt^46^ (Universal Protein Resource)

UniProt is a leading repository of protein sequence and functional information. It plays a dual role by linking transcripts to their corresponding protein products. In addition to capturing transcript-protein interactions, UniProt also includes annotations of protein-disease relationships, making it essential for understanding how protein dysfunctions can lead to disease.

##### BioGrid^72^

BioGrid is a key resource for protein-protein interaction data, curated from both high-throughput and small-scale experimental studies. This database is essential for exploring how proteins interact within cellular networks, facilitating the study of complex biological processes such as signaling pathways, metabolic networks, and structural assemblies. BioGrid data on protein-protein interactions supports a wide range of applications, from basic research to drug discovery.

##### STRING^74^

STRING is a database of known and predicted protein-protein interactions, integrating data from various sources such as experimental studies, computational predictions, and publicly available text collections. It is essential for understanding the functional interactions between proteins and mapping protein interaction networks. STRING helps to identify potential interactions that play critical roles in biological processes and disease states, making it a valuable tool for systems biology research.

##### KEGG^48^ (Kyoto Encyclopedia of Genes and Genomes)

KEGG is a comprehensive database that integrates genomic, chemical, and systemic functional information, offering valuable insights into various biological interactions. It is essential for studying protein-protein interactions, illustrating how proteins cooperate in cellular processes, as well as gene-pathway interactions, showing how genes function within specific biological pathways. Furthermore, KEGG explores drug-pathway interactions, revealing how drugs influence these pathways, and facilitates the study of pathway-gene and pathway-drug interactions, providing a clear understanding of how pathways are regulated by genes and targeted by drugs.

##### HPO^59^ (Human Phenotype Ontology)

HPO provides a standardized vocabulary of phenotypic abnormalities associated with human diseases. It is invaluable for connecting genes to phenotypes (gene-phenotype interaction), linking diseases to their phenotypic presentations (disease-phenotype interaction), and mapping genes to diseases (gene-disease interaction). HPO also facilitates the study of phenotype-phenotype relationships, enabling researchers to compare phenotypic similarities and differences across genetic conditions.

##### DisGeNet^75^

DisGeNet is a comprehensive platform that integrates data on gene-disease associations from multiple sources, including expert-curated databases, scientific literature, and publicly available repositories. It plays a critical role in identifying gene-disease interactions, helping to elucidate the genetic basis of various diseases. DisGeNet supports research into disease mechanisms by providing insights into the complex genetic networks that underlie disease phenotypes.

##### DISEASES^76^

The DISEASES database provides information on protein-disease associations, integrating data from literature mining and manually curated sources. It links proteins to the diseases they are associated with, offering a detailed view of how protein dysfunctions contribute to disease phenotypes. DISEASES is especially useful for identifying molecular mechanisms underlying diseases and for exploring potential therapeutic targets.

##### HMDB^52^ (Human Metabolome Database)

HMDB is an extensive resource that provides detailed information on human metabolites, including drugs, drug metabolites, and endogenous small molecules. It captures a wide range of interactions, including drug-metabolome, metabolome-disease, and metabolome-protein relationships. HMDB supports research in metabolomics, systems biology, and pharmacology, providing data on metabolic pathways, metabolite-protein interactions, and the role of metabolites in health and disease.

##### MetaNetX^77^

It is a comprehensive resource developed by the SIB Swiss Institute of Bioinformatics to facilitate the standardization, integration, and analysis of genome-scale metabolic networks (GSMNs) and biochemical pathways. MetaNetX allows users to construct, modify, and analyze metabolic models through tools for flux balance analysis (FBA), reaction knockout simulations, and network comparison. By integrating data from diverse sources and providing a standardized framework, MetaNetX is a valuable tool for researchers in systems biology and bioinformatics, enabling a deeper understanding of complex metabolic processes.

##### DisBiome^78^

DisBiome is a database that focuses on the relationships between microbiomes and diseases. It captures microbiome-disease interactions, providing insights into how microbial taxa are associated with health and disease. DisBiome supports research into the role of the human microbiome in various disease conditions, facilitating the exploration of microbial communities as potential biomarkers or therapeutic targets.

##### MDAD^79^ (Microbe-Drug Association Database)

MDAD is a comprehensive resource that compiles clinically and experimentally validated associations between microbes and drugs. It contains 5,055 entries, encompassing 1,388 drugs and 180 microbes, sourced from multiple drug databases and scientific publications. Each record in MDAD includes detailed annotations, such as molecular forms of drugs, links to DrugBank, microbe target information from UniProt, and original reference citations. This database serves as a valuable tool for researchers aiming to understand microbe-drug interactions, facilitating advancements in drug discovery, disease therapy, and personalized medicine.

##### PharmacoMicrobiomics^80^

It is a field that examines the interactions between the human microbiome and drugs, focusing on how microbial communities influence drug metabolism, efficacy, and toxicity. This bidirectional relationship involves microbes activating, inactivating, or transforming drugs into metabolites with altered effects, while drugs, in turn, can reshape the composition and function of the microbiome. These interactions have profound implications for personalized medicine, as variations in the microbiome can affect individual drug responses, side effects, and therapeutic outcomes. By understanding these dynamics, PharmacoMicrobiomics aims to optimize drug therapies, reduce adverse effects, and pave the way for microbiome-targeted medical interventions.

##### CTD (The Comparative Toxicogenomics Database)^58^

CTD is a publicly available, manually curated resource that provides insights into the complex relationships between chemicals, genes, and diseases, with a specific emphasis on environmental exposures. CTD integrates data on chemical-gene interactions, chemical-disease associations, and gene-disease relationships, offering researchers a unique platform to explore the molecular mechanisms underlying toxicological effects and exposure-related health outcomes. By including exposure-related information, CTD helps bridge the gap between environmental science and molecular biology, enabling studies on how environmental factors influence gene function and contribute to disease etiology. This resource is particularly valuable for advancing research in toxicogenomics, precision medicine, and environmental health.

##### DO (Disease Ontology)^63^

DO is a standardized biomedical ontology that provides a structured vocabulary and hierarchical classification for human diseases, enabling consistent annotation and integration of disease-related data across research and clinical domains. Each disease entry is assigned a unique identifier and is cross-referenced with external resources such as OMIM, ICD, SNOMED CT, and MeSH, ensuring interoperability and facilitating data harmonization. By linking diseases to their etiology, molecular mechanisms, and clinical manifestations, DO supports applications in translational medicine, computational biology, and precision medicine. Its integration with genomic and phenotypic datasets makes it a critical tool for advancing disease research, biomarker discovery, and therapeutic development.

##### DrugBank^70^

DrugBank is a comprehensive resource that integrates detailed information on drugs and their targets. It captures multiple types of interactions, including protein-drug, drug-drug relationships. DrugBank provides data on drug mechanisms, drug interactions, and the diseases they are used to treat, making it an essential tool for pharmacological research and drug development. It also supports studies on how drugs interact with biological systems at the molecular level.

##### BindingDB^81^

BindingDB is a public repository of measured binding affinities between proteins (mainly drug targets) and small, drug-like molecules. It supports research into protein-drug interactions by providing experimental data on the binding affinities of drugs to their target proteins. BindingDB is a valuable resource for drug discovery and pharmacology, helping researchers identify potential drug candidates and understand the molecular mechanisms of drug action.

**DrugCentral**^82^ DrugCentral is a centralized portal for drug information, offering data on drug-protein, drug-disease, interactions. It integrates information on drug indications, targets, and mechanisms of action, supporting the study of therapeutic interventions and pharmacodynamics. DrugCentral is an important resource for researchers exploring drug repurposing, drug development, and clinical applications.

##### SIDER^83^ (Side Effect Resource)

SIDER provides comprehensive data on the adverse effects of drugs, linking pharmaceutical compounds to their phenotypic side effects. This resource is essential for studying drug-phenotype interactions, helping researchers understand the unintended consequences of drug use. SIDER supports pharmacovigilance efforts and aids in optimizing drug safety profiles by highlighting potential risks associated with pharmaceutical compounds.

**Table S1.**
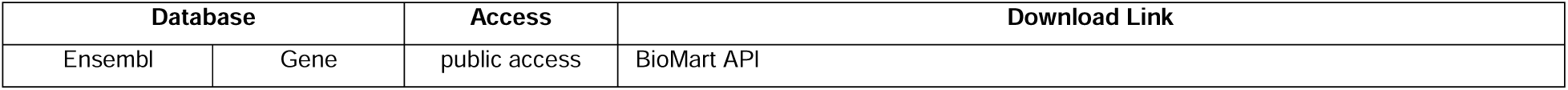
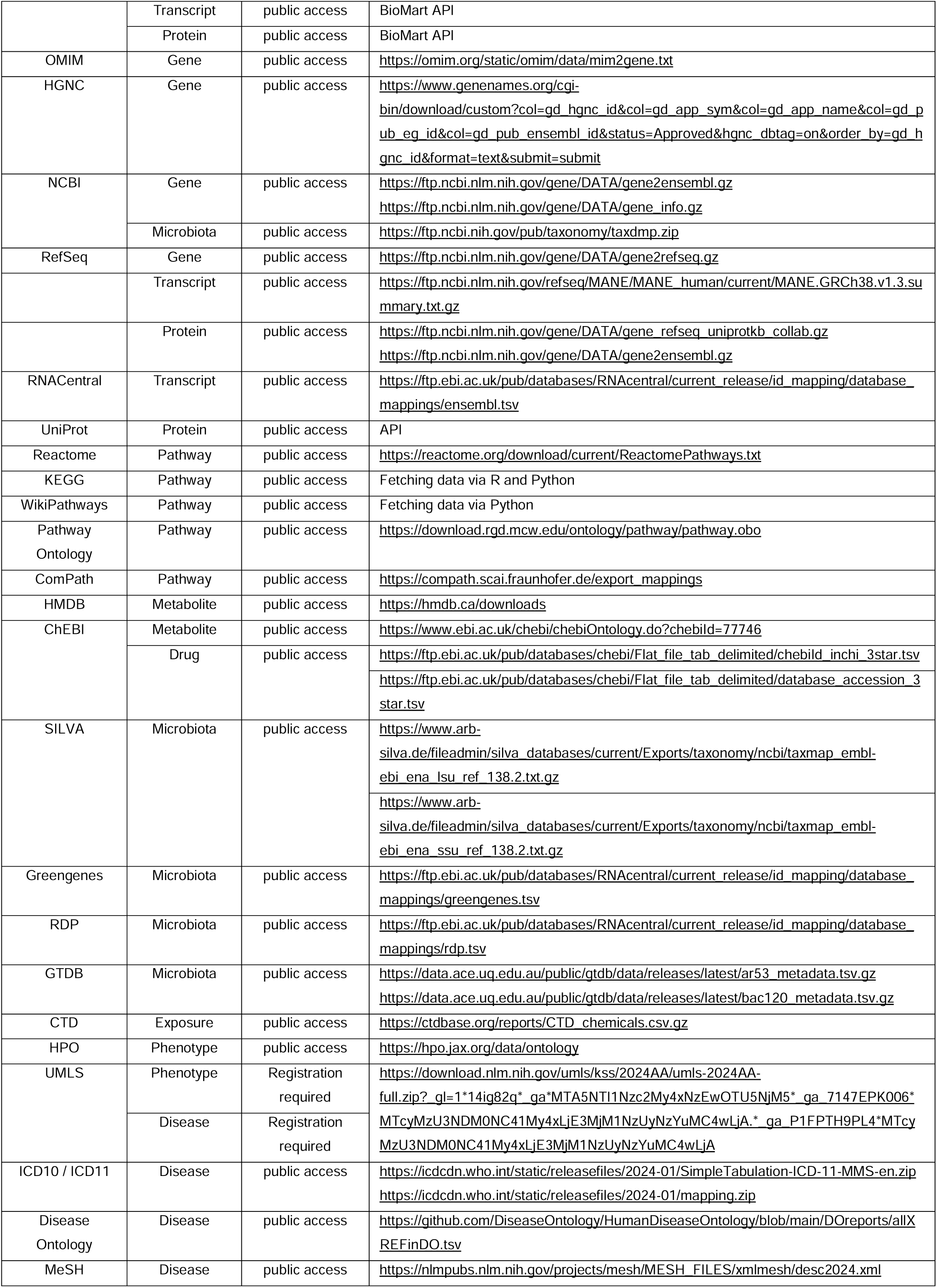
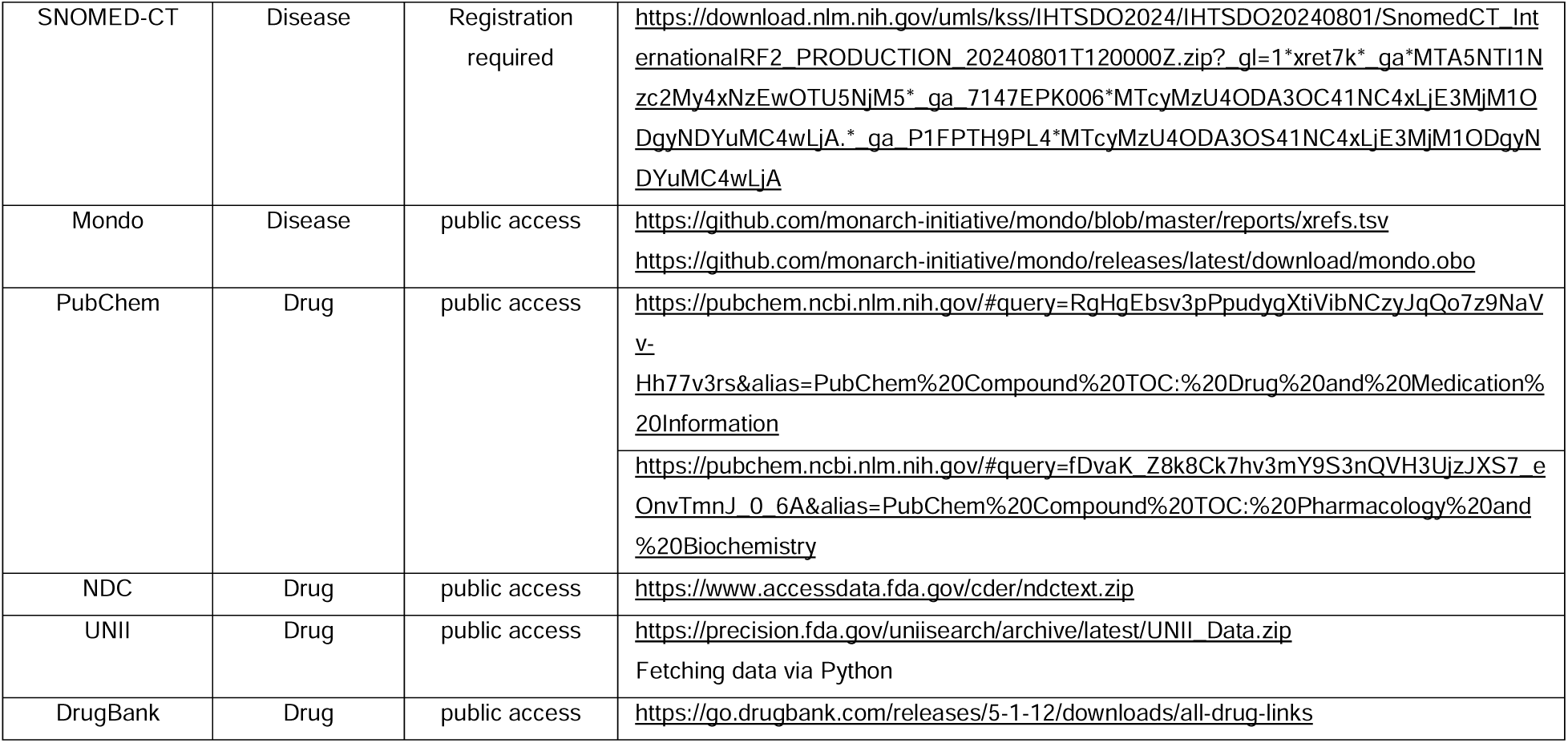
Download Links and Access Control for Entity Databases.

**Table S2.**
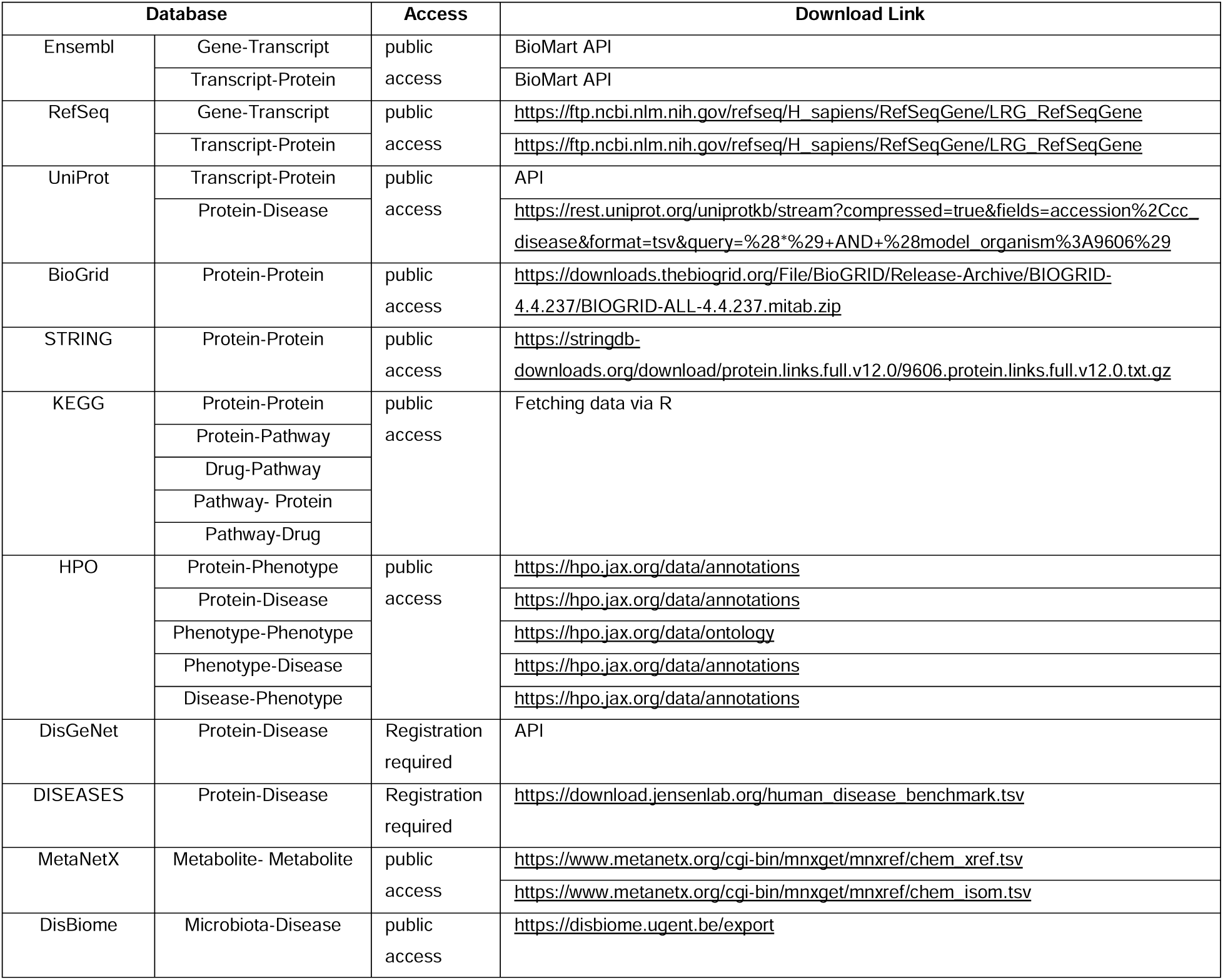
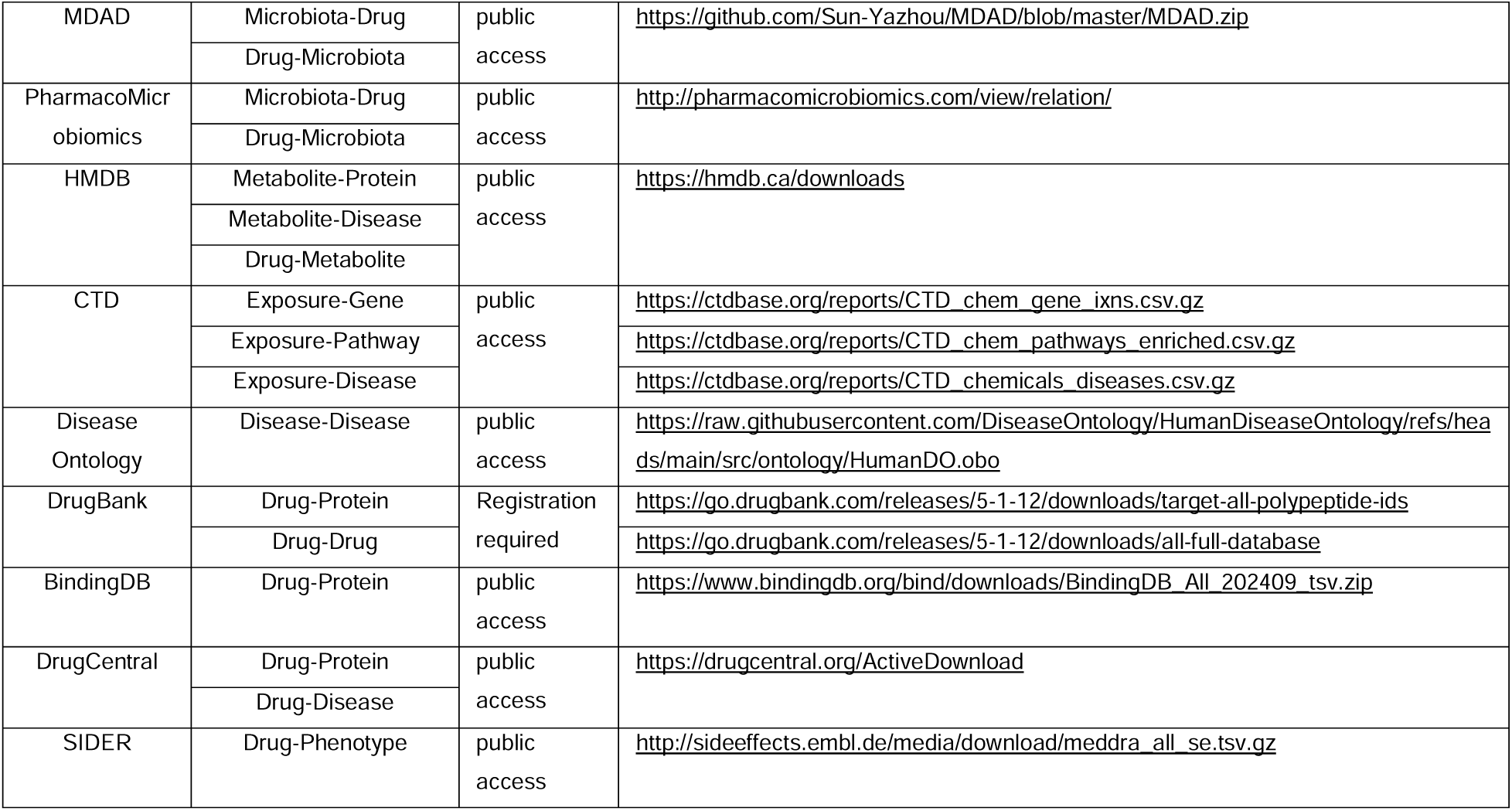
Download Links and Access Control for Relation Databases.

## Section B. Details of Entity and Relation Integration

### B.1 Entity Integration

#### Gene Entity Merging

The Ensembl database was utilized as the primary basis for data integration. Initially, data from Ensembl, HGNC, and NCBI were merged based on matching Ensembl IDs. Subsequently, data from RefSeq and OMIM were incorporated, with NCBI IDs serving as the common identifier. The NCBI ID was chosen as the minimal unit for unifying the data, and a final integration of gene information was conducted according to NCBI IDs (refer to Table S3 and Figure S1 in the supplementary section for details). The columns highlighted in bold within the table denote those used for merging across databases, with the IDs in these columns being unique. Additionally, a textual description of each gene entity was appended. Using the free Perl script geneDocSum.pl provided by NCBI (download link: https://ftp.ncbi.nih.gov/gene/tools/geneDocSum.pl), all human records marked as current (alive) and containing summaries were retrieved. By mapping NCBI Gene IDs to corresponding entries in BioMedGraphica_Gene, the descriptions associated with BioMedGraphica_Gene IDs (BMG_GN) were obtained.

**Table S3.**
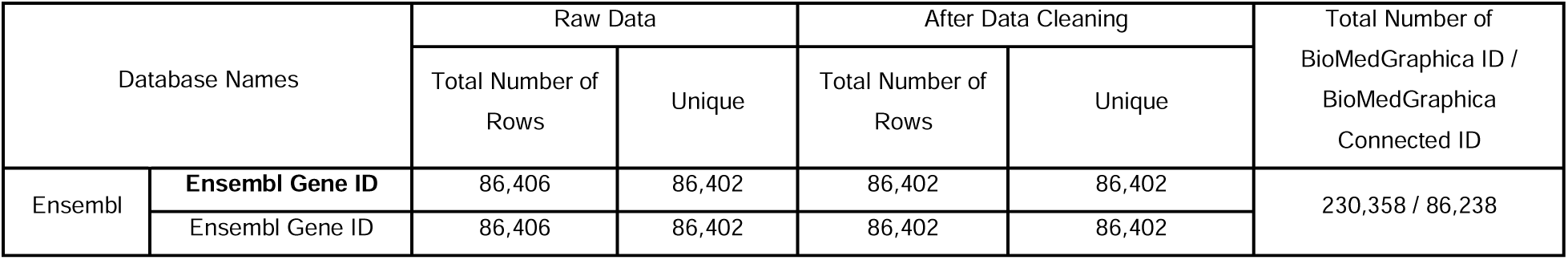
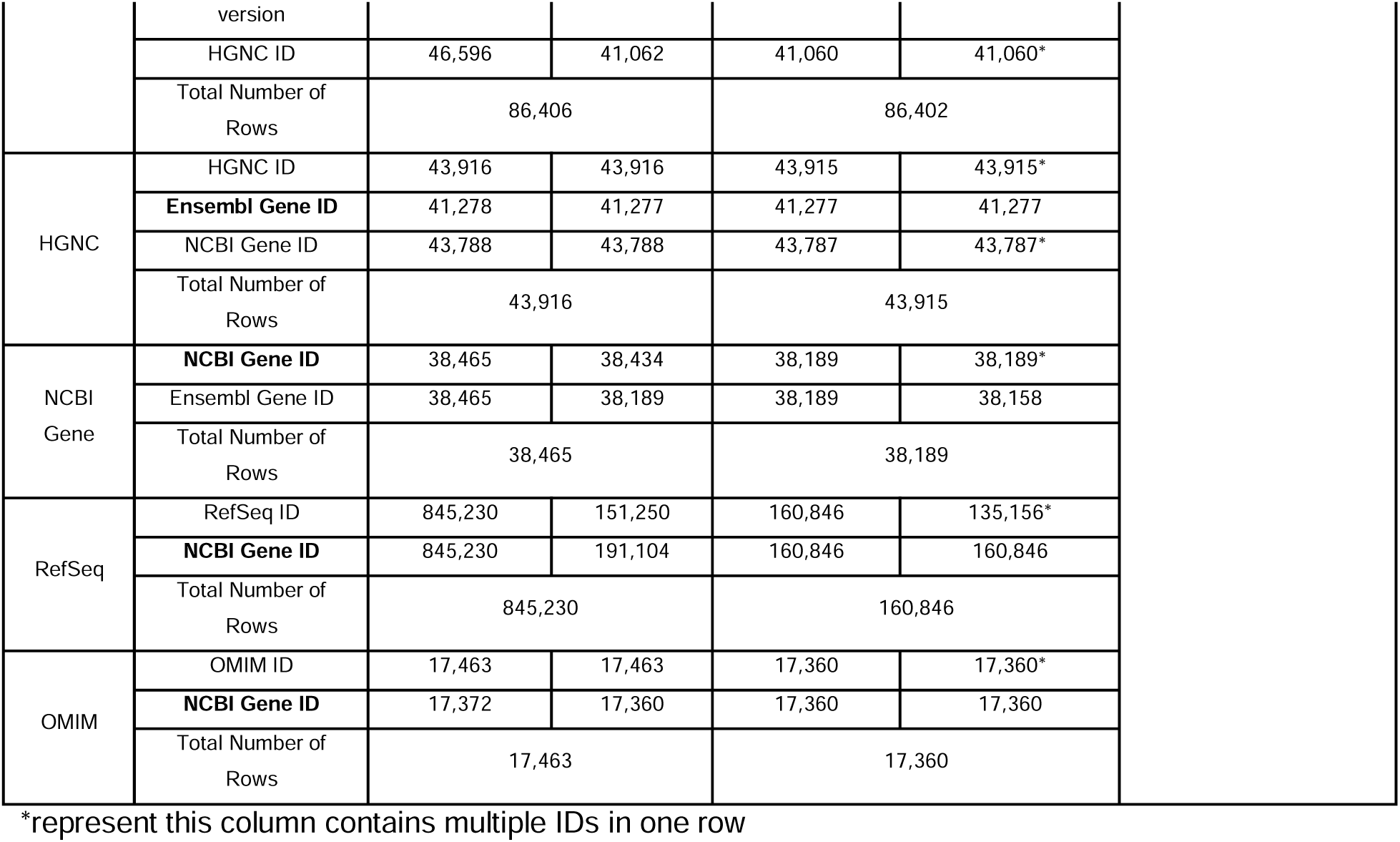
Gene Entity Information.

| Database Names |  | Raw Data |  | After Data Cleaning |  | Total Number of BioMedGraphica ID / BioMedGraphica Connected ID |
| --- | --- | --- | --- | --- | --- | --- |
|  |  | Total Number of Rows | Unique | Total Number of Rows | Unique |  |
| Ensembl | **Ensembl Gene ID** | 86,406 | 86,402 | 86,402 | 86,402 | 230,358 / 86,238 |
|  | Ensembl Gene ID | 86,406 | 86,402 | 86,402 | 86,402 |  |

|  |  |  |  |  |  |
| --- | --- | --- | --- | --- | --- |
|  | version |  |  |  |  |
|  | HGNC ID | 46,596 | 41,062 | 41,060 | 41,060* |
|  | Total Number of Rows | 86,406 |  | 86,402 |  |
| HGNC | HGNC ID | 43,916 | 43,916 | 43,915 | 43,915* |
|  | Ensembl Gene ID | 41,278 | 41,277 | 41,277 | 41,277 |
|  | NCBI Gene ID | 43,788 | 43,788 | 43,787 | 43,787* |
|  | Total Number of Rows | 43,916 |  | 43,915 |  |
| NCBI Gene | NCBI Gene ID | 38,465 | 38,434 | 38,189 | 38,189* |
|  | Ensembl Gene ID | 38,465 | 38,189 | 38,189 | 38,158 |
|  | Total Number of Rows | 38,465 |  | 38,189 |  |
| RefSeq | RefSeq ID | 845,230 | 151,250 | 160,846 | 135,156* |
|  | NCBI Gene ID | 845,230 | 191,104 | 160,846 | 160,846 |
|  | Total Number of Rows | 845,230 |  | 160,846 |  |
| OMIM | OMIM ID | 17,463 | 17,463 | 17,360 | 17,360* |
|  | NCBI Gene ID | 17,372 | 17,360 | 17,360 | 17,360 |
|  | Total Number of Rows | 17,463 |  | 17,360 |  |

\*represent this column contains multiple IDs in one row

**Figure S1.**
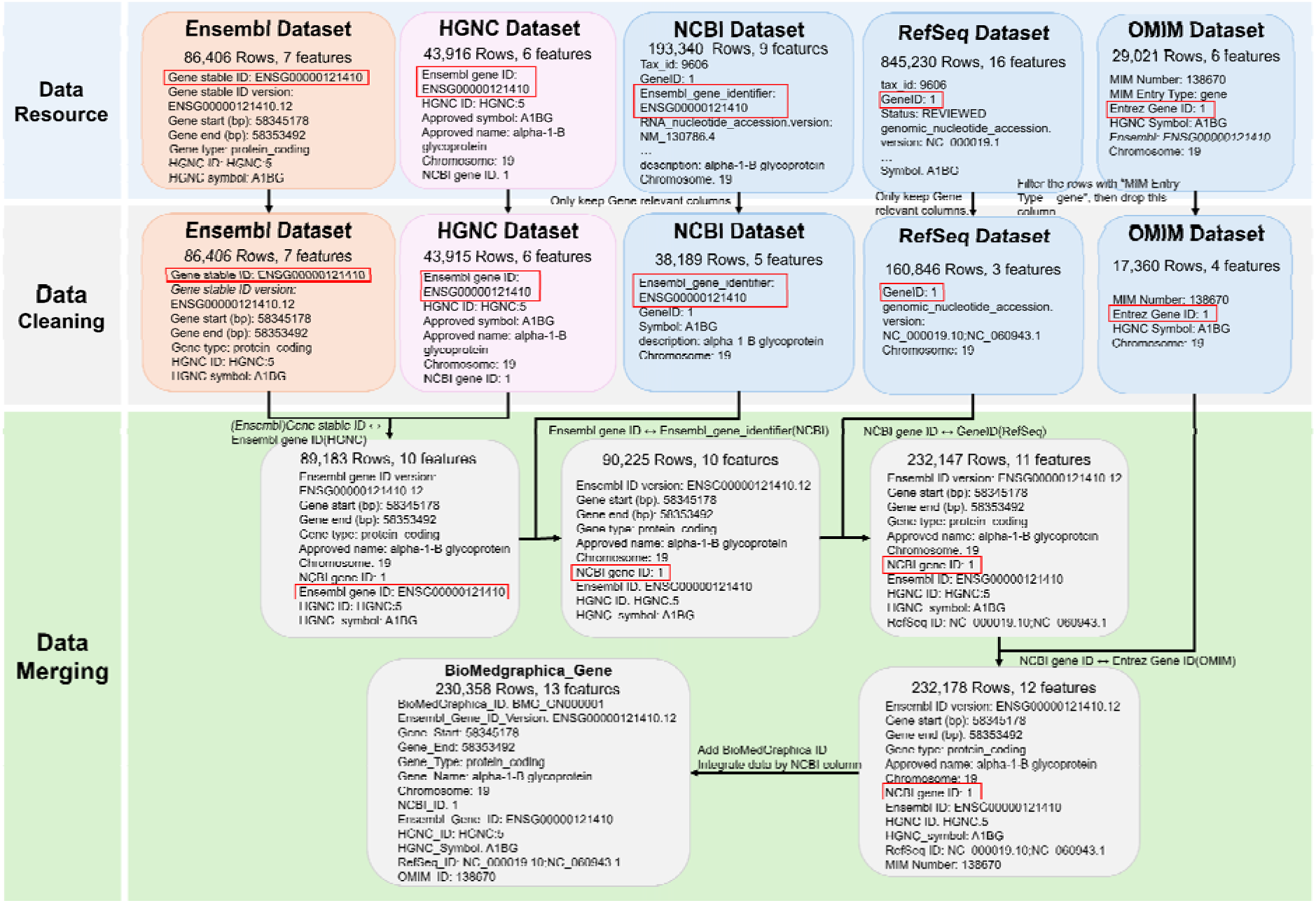
Details of Gene Entity Merging Process.

**Figure S1** provides a detailed overview of the integration process for BioMedGraphica Gene, using A1BG as an example. The “Data Resource” section depicts the original datasets sourced from various databases for gene entity integration. The “Data Cleaning” section presents the cleaned data format prepared for integration. Columns highlighted in red boxes indicate the key matching fields used during the merging process. In the “Data Merging” section, the gray boxes showcase the data format at each step of database integration. The overall gene entity integration employs an outer join approach: Ensembl and HGNC databases are merged first, followed by integration with the NCBI Gene database. Subsequently, RefSeq and OMIM are incorporated sequentially. The final unification is based on the NCBI Gene ID, ensuring that all entries with the same ID are consolidated.

#### Transcript Entity Merging

The integration of the three databases utilized the Ensembl ID as the standard reference. For transcript entities, the Ensembl Transcript Stable ID was adopted as the smallest unit of data granularity. The integration process is illustrated in **Figure S2** of the supplementary section, and the merged results are detailed in **Table S4**. Bolded entries in the table identify the columns used for database merging, where the IDs in these columns are unique. Transcript descriptions were extracted from the Ensembl database using the BioMart API. By mapping the Transcript Stable ID to corresponding transcripts in BioMedGraphica, transcript descriptions were successfully assigned to the majority of BioMedGraphica transcripts.

**Table S4.** Transcript Entity Information.

| Database |  | Raw Data |  | After Data Cleaning |  | Total Number of BioMedGraphica ID / BioMedGraphica Connected ID |
| --- | --- | --- | --- | --- | --- | --- |
|  |  | Total Number of Rows | Unique | Total Number of Rows | Unique |  |
| Ensembl | Ensembl Transcript ID | 451,959 | 412,034 | 451,959 | 412,034 | 412,326 / 412,039 |
|  | Ensembl Transcript ID version | 451,959 | 412,034 | 451,959 | 412,034 |  |
|  | Ensembl Gene ID | 451,959 | 86,402 | 451,959 | 86,402 |  |
|  | RefSeq mRNA ID | 85,642 | 66,939 | 85,642 | 66,939 |  |
|  | RefSeq ncRNA ID | 35,946 | 20,199 | 35,946 | 20,199 |  |
|  | RefSeq MANE Select ID | 38,501 | 19,287 | 38,501 | 19,287 |  |
|  | Total Number of Rows | 451,959 |  | 451,959 |  |  |
| RefSeq | Ensembl Transcript ID version | 19,404 | 19,404 | 19,404 | 19,404 |  |
|  | RefSeq ID | 19,404 | 19,404 | 19,404 | 19,404 |  |
|  | Total Number of Rows | 19,404 |  | 19,404 |  |  |
| RNAcentral | RNAcentral ID | 66,789 | 62,925 | 66,789 | 62,925 |  |
|  | Ensembl Transcript ID | 66,789 | 66,789 | 66,789 | 66,789 |  |
|  | Total Number of Rows | 66,789 |  | 66,789 |  |  |

\*represent this column contains multiple IDs in one row

**Figure S2.**
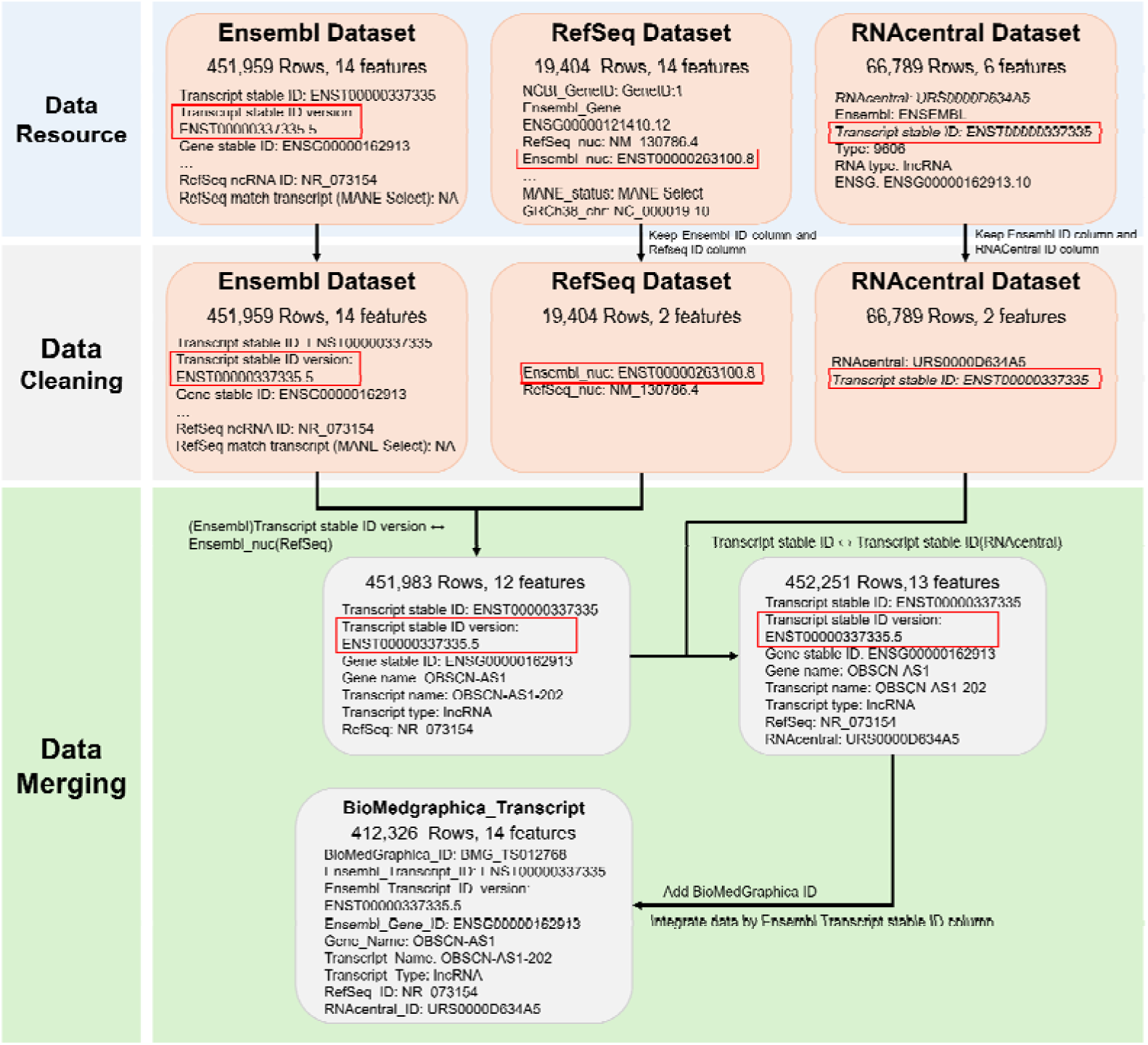
Details of Transcript Entity Merging Process.

A detailed depiction of the integration process for BioMedGraphica transcript has been provided **in Figure S2**, using ENST00000337335.5 as an example. The “Data Resource” section shows the raw data sourced from databases used in transcript entity integration. Since the original RefSeq dataset lacked a corresponding RefSeq ID for ENST00000337335.5, an alternative transcript was selected as a supplementary example. The “Data Cleaning” section presents the cleaned data format prepared for integration. Columns highlighted in red boxes indicate the key matching fields used during the merging process. In the “Data Merging” section, gray boxes illustrate the data format after each step of database integration. The integration process for transcript entities employs an outer join approach: first, the Ensembl and RefSeq databases are merged, followed by integration with the RNAcentral database. The Ensembl stable ID serves as the primary unit for final data unification, consolidating all entries with the same Ensembl stable ID.

#### Protein Entity Merging

The integration process began by merging data from Ensembl and UniProt based on the Protein Stable ID Version. Subsequently, RefSeq data was incorporated by leveraging mapping relationships between RefSeq and the two databases. The Ensembl Protein ID Version was established as the minimal unit of data granularity for protein entities (refer to **Figure S3** in the supplementary section for the merging workflow and **Table S5** for detailed results). Bolded entries in the table highlight the columns used for cross-database merging, where the IDs are uniquely assigned. Protein descriptions were retrieved from the UniProt database using the UniProt API. By mapping UniProt IDs to corresponding proteins in BioMedGraphica, descriptive information was successfully provided for BioMedGraphica proteins.

**Table S5.**
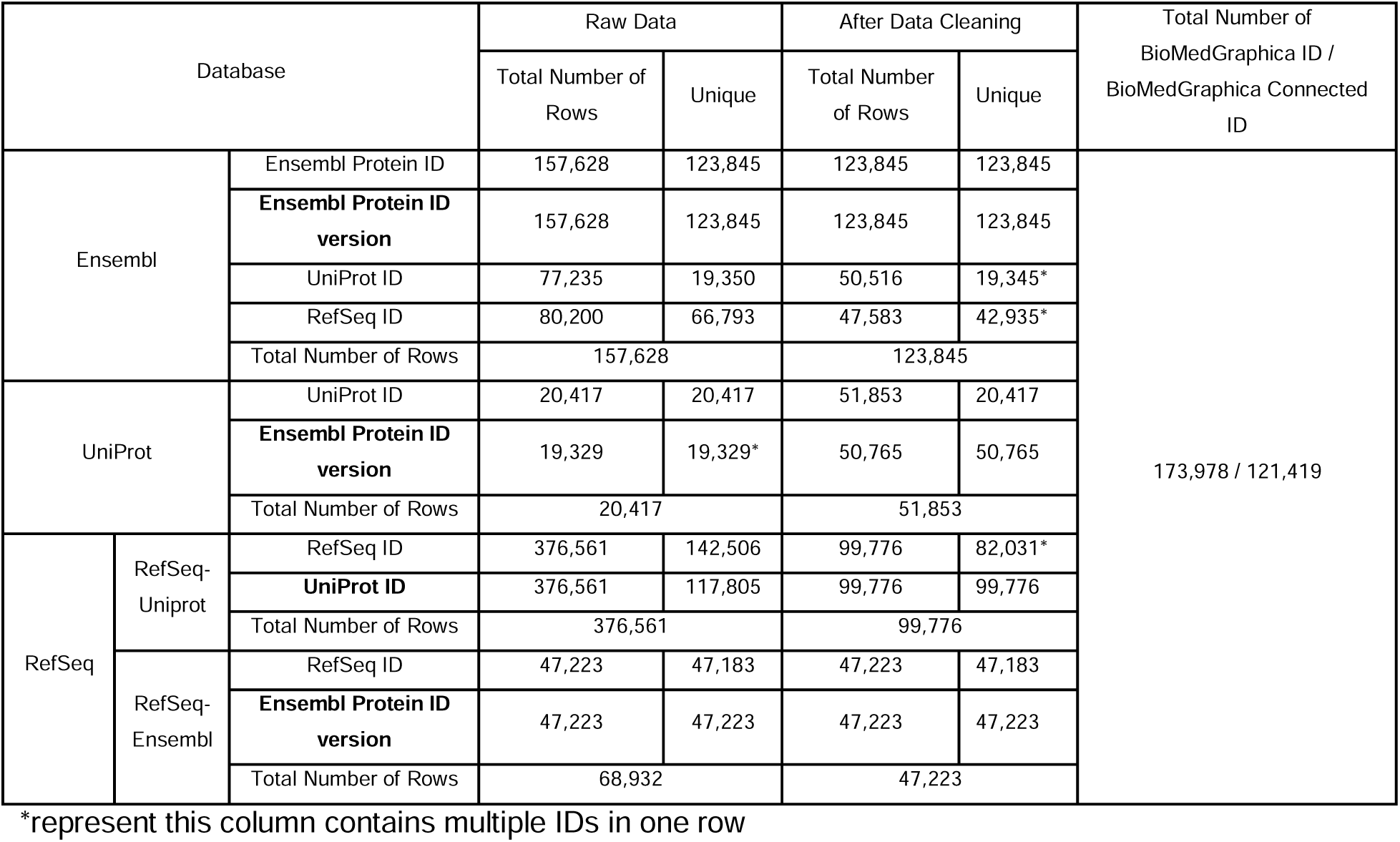
Protein Entity Information.

**Figure S3.**
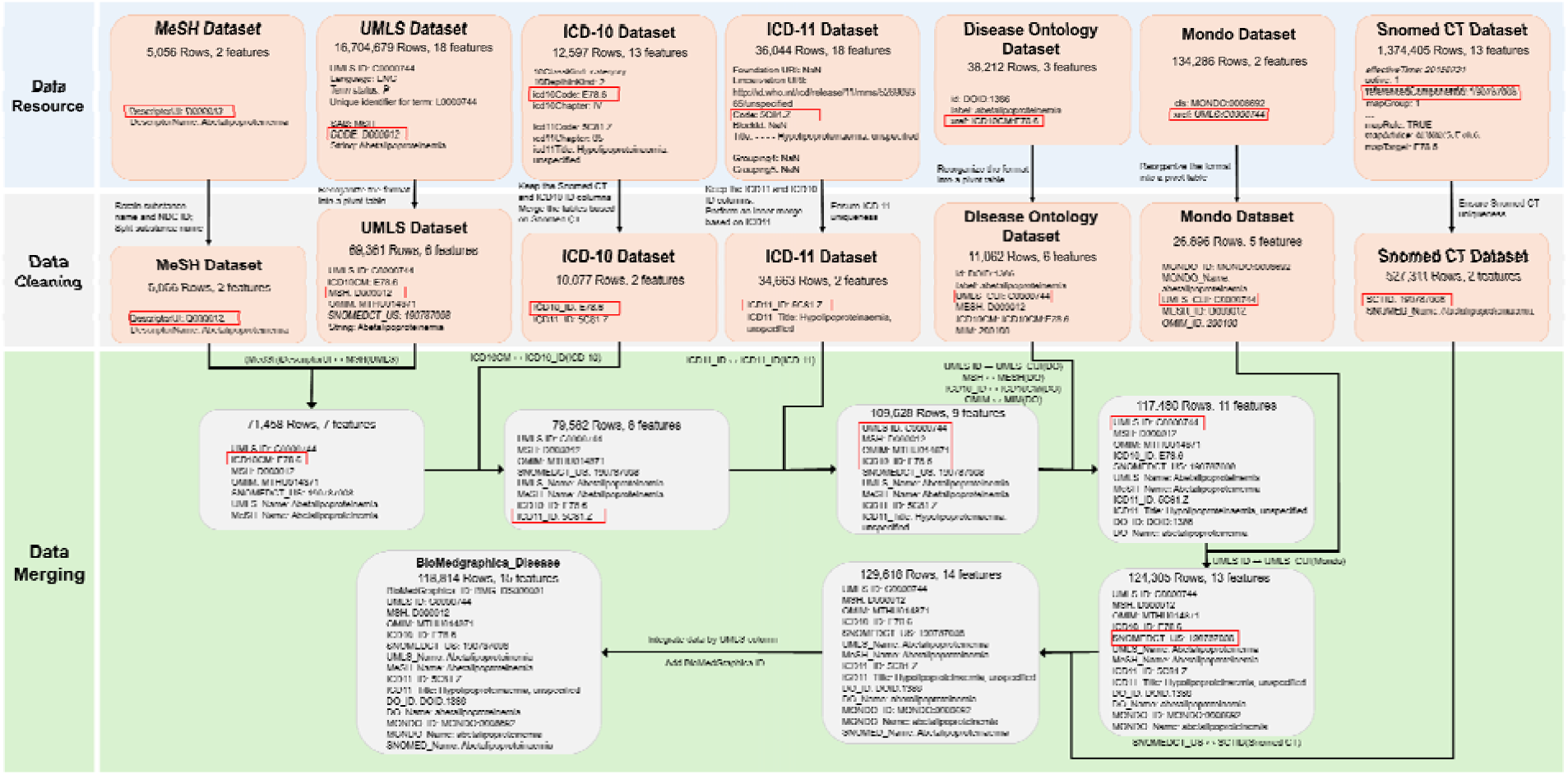
Details of Protein Entity Merging Process.

**Figure S3** illustrates the integration process for BioMedGraphica Protein, using ENSP00000000233.5 a a representative example. The “Data Resource” section outlines the raw datasets obtained from various databases utilized in protein entity integration. The “Data Cleaning” section highlights the standardized format of the data after preparation for integration. Key matching columns, marked in red boxes, were used to align data across sources. The “Data Merging” section visualizes the transformation of data formats through successive integration steps, represented by gray boxes. The integration process employs an outer join methodology, starting with the merging of Ensembl and UniProt databases. This combined dataset is then integrated with RefSeq. The final step uses the Ensembl stable ID version as the primary key to unify entries, ensuring that all records associated with the same Ensembl stable ID version are consolidated.

#### Pathway Entities Integration

The data integration process began with Pathway Ontology (PO) as the foundational framework, merging datasets from PO, KEGG, and Reactome. Missing data was subsequently addressed through equivalent mapping relationships between KEGG and Reactome, as provided by ComPath. Finally, human pathway data from WikiPathway was integrated using equivalent mappings between KEGG and WikiPathway also facilitated by ComPath. Bolded columns in the table represent the fields used for merging with other databases, where the IDs in these columns are uniquely assigned (refer to **Figure S4** in the supplementary section for the detailed integration workflow and **Table S6** for results).

**Table S6.** Pathway Entity Information.

| Database |  | Raw Data |  | After Data Cleaning |  | Total Number of BioMedGraphica ID / BioMedGraphica Connected ID |
| --- | --- | --- | --- | --- | --- | --- |
|  |  | Total Number of Rows | Unique | Total Number of Rows | Unique |  |
| Pathway Ontology | PO ID | 2677 | 2677 | 2677 | 2677 | 6,793 / 1,930 |
|  | KEGG ID | 237 | 211 | 201 | 179 |  |
|  | Reactome ID | 327 | 321 | 327 | 321 |  |
|  | Total Number of Rows | 2677 |  | 2677 |  |  |
| KEGG | KEGG ID | 365 | 365 | 365 | 365 |  |
|  | Total Number of Rows | 365 |  | 365 |  |  |
| Reactome | Reactome ID | 2751 | 2751 | 2751 | 2751 |  |
|  | Total Number of Rows | 2751 |  | 2751 |  |  |
| ComPath | KEGG ID | 953 | 859 | 113 | 89 |  |
|  | Reactome ID | 1,274 | 1,055 | 58 | 57 |  |
|  | WikiPathway ID | 940 | 724 | 55 | 55 |  |
|  | Total Number of Rows | 1,592 |  | 113 |  |  |
| WikiPathways | WikiPathways ID | 1,534 | 1,534 | 1,534 | 1,534 |  |
|  | Total Number of Rows | 1,534 |  | 1,534 |  |  |

\*represent this column contains multiple IDs in one row
Values separated by semicolons indicate data from multiple files

**Figure S4.**
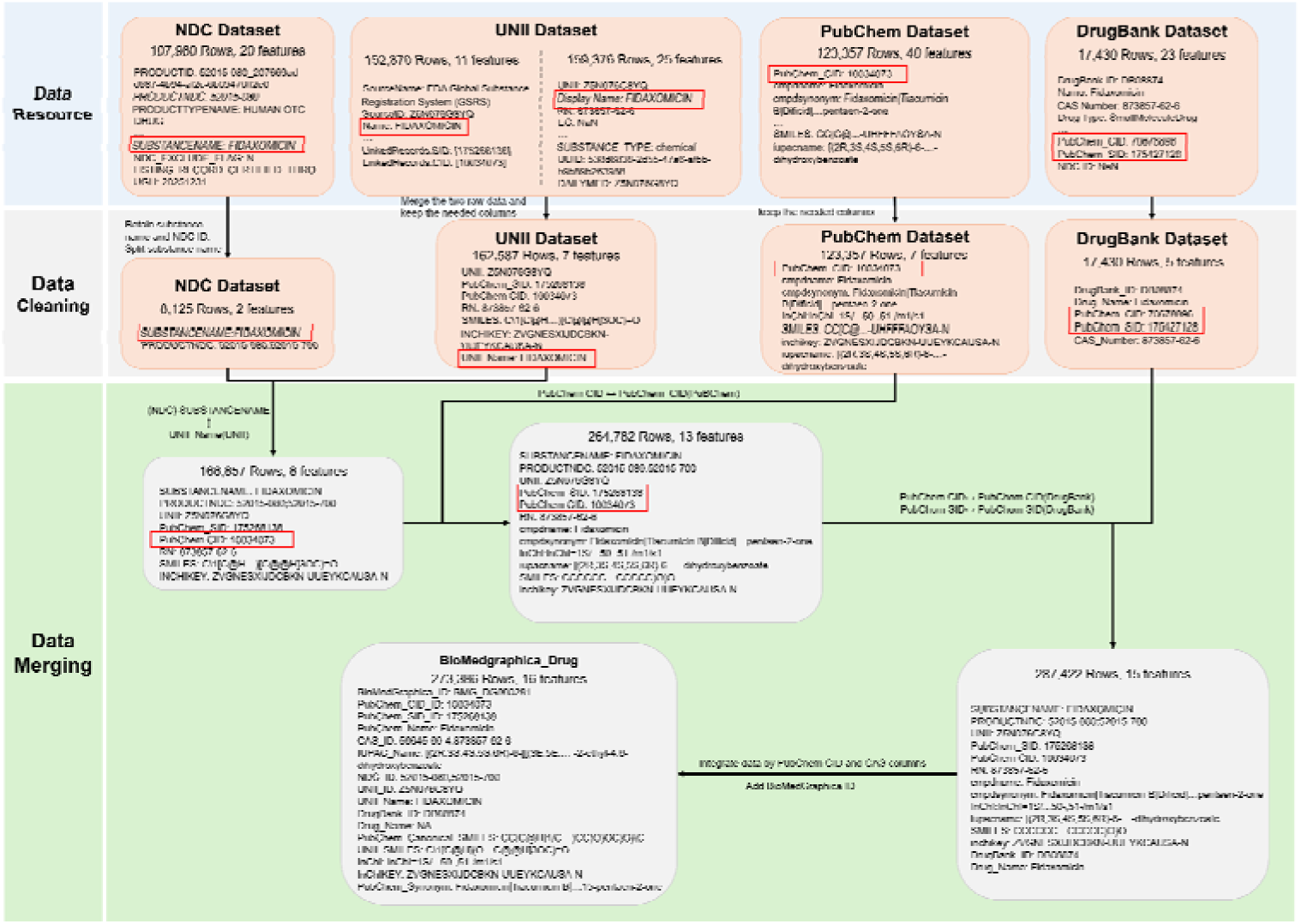
Details of Pathway Entity Merging Process.

**Figure S4** provides a detailed illustration of the integration process for BioMedGraphica Pathway, using PW:0000009 as an example. The “Data Resource” section represents the raw data from the databases used in the integration of the pathway entity. The “Data Cleaning” section displays the format of the cleaned data prepared for integration. The data highlighted in the red boxes indicates the key matching columns used for merging. In the “Data Merging” section, the gray boxes show the format of the data after each step of database integration. The pathway entity integration process follows an outer join method. First, the Pathway Ontology and KEGG databases are merged, followed by the integration of Reactome data into the combined dataset, and subsequently ComPath and WikiPathway are integrated in sequence.

#### Metabolite Entities Integration

The integration of the two databases utilized the ChEBI ID as the primary linking key. Subsequently, entries with identical HMDB IDs were consolidated, establishing the HMDB ID as the smallest unit of data granularity. Columns highlighted in bold within the table denote those used for database merging, ensuring the uniqueness of IDs in these columns (see **Figure S5** in the supplementary section for details on the merging process and **Table S7** for the results).

**Table S7.**
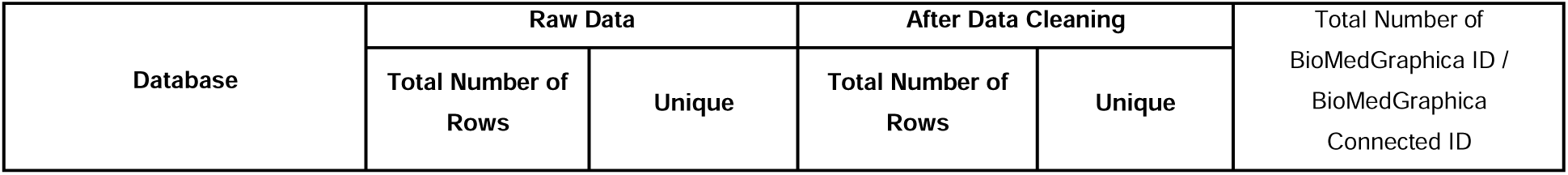
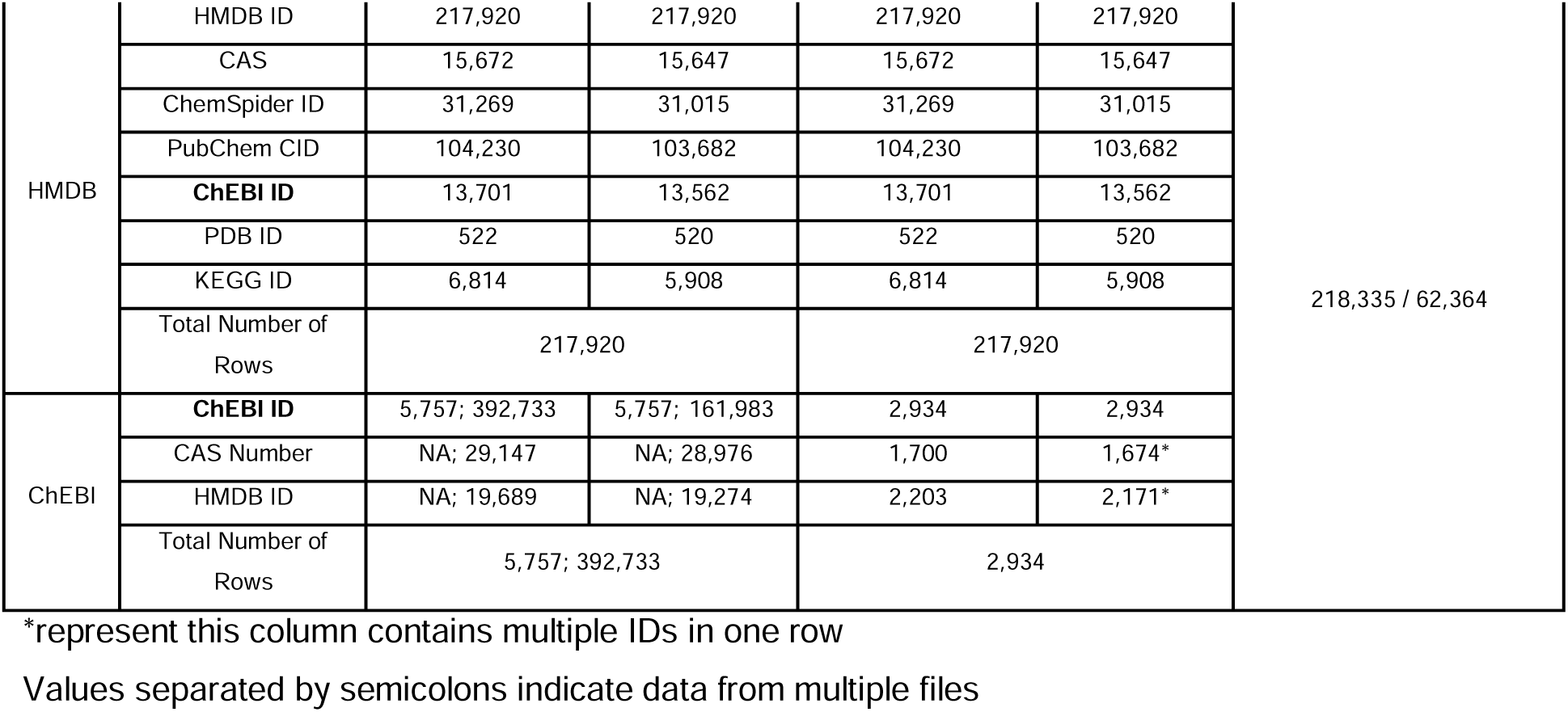
Metabolite Entity Information.

**Figure S5.**
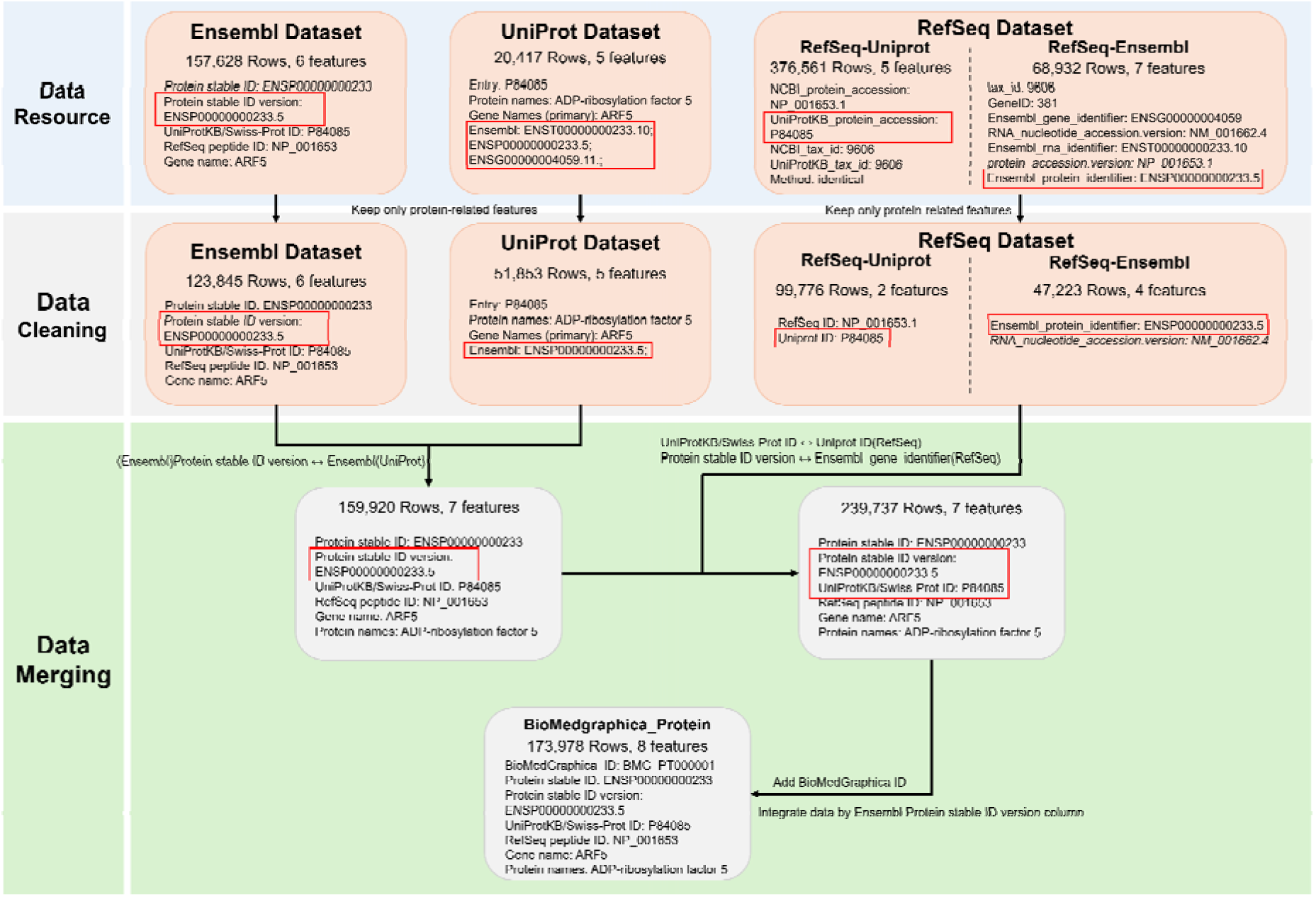
Details of Metabolite Entity Merging Process.

**Figure S5** provides a detailed illustration of the integration process for BioMedGraphica Metabolite, using ChEBI: 18019 as an example. The “Data Resource” section represents the raw data from the database used in the integration of the metabolite entity. The “Data Cleaning” section displays the format of the cleaned data prepared for integration. The data highlighted in the red boxes indicates the key matching columns used for merging. In the “Data Merging” section, the gray boxes show the data format after each step of database integration. The metabolite entity integration process follows an outer join method, merging data from the HMDB and ChEBI databases. Finally, the HMDB ID is used as the minimal unit for data unification, consolidating all entries with the same HMDB ID.

#### Microbiota Entities Integration

The NCBI Taxon ID was employed as the standard key for harmonizing data across all microbiota datasets. This identifier was chosen due to its widespread presence in the included databases, enabling data merging. Columns highlighted in bold within the accompanying table indicate those used for cross-database integration, ensuring the uniqueness of IDs in these fields. For a detailed explanation of the integration methodology, refer to **Figure S6** in the supplementary section, with comprehensive results presented in **Table S8**.

**Table S8.**
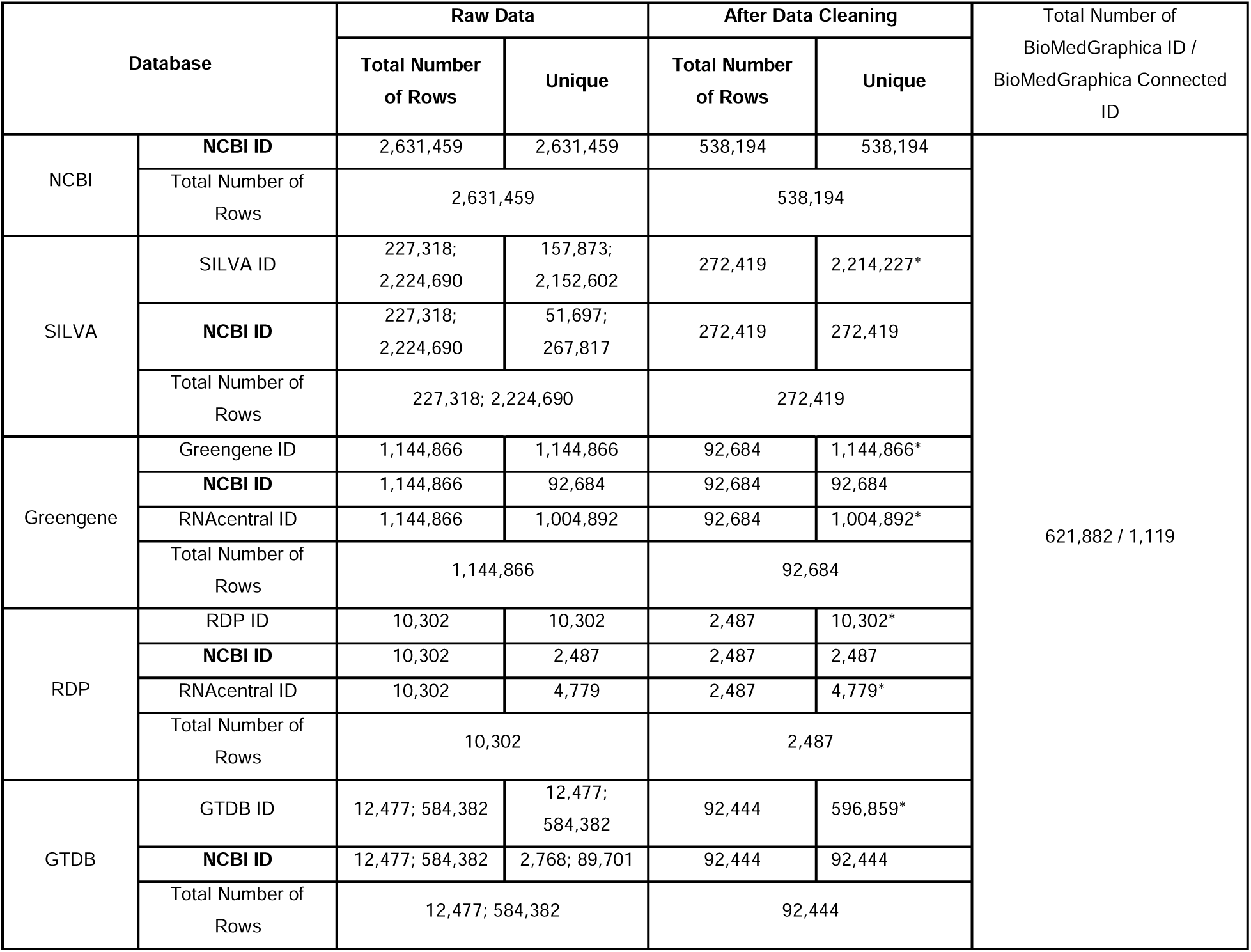
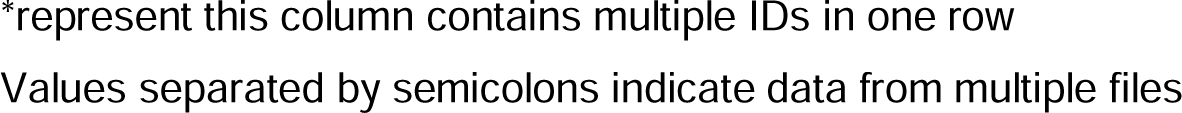
Microbiota Entities Information.

| Database |  | Raw Data |  | After Data Cleaning |  | Total Number of<br>BioMedGraphica ID /<br>BioMedGraphica Connected<br>ID |
| --- | --- | --- | --- | --- | --- | --- |
|  |  | Total Number<br>of Rows | Unique | Total Number<br>of Rows | Unique |  |
| NCBI | NCBI ID | 2,631,459 | 2,631,459 | 538,194 | 538,194 | 621,882 / 1,119 |
|  | Total Number of<br>Rows | 2,631,459 |  | 538,194 |  |  |
| SILVA | SILVA ID | 227,318;<br>2,224,690 | 157,873;<br>2,152,602 | 272,419 | 2,214,227* |  |
|  | NCBI ID | 227,318;<br>2,224,690 | 51,697;<br>267,817 | 272,419 | 272,419 |  |
|  | Total Number of<br>Rows | 227,318; 2,224,690 |  | 272,419 |  |  |
| Greengene | Greengene ID | 1,144,866 | 1,144,866 | 92,684 | 1,144,866* |  |
|  | NCBI ID | 1,144,866 | 92,684 | 92,684 | 92,684 |  |
|  | RNAcentral ID | 1,144,866 | 1,004,892 | 92,684 | 1,004,892* |  |
|  | Total Number of<br>Rows | 1,144,866 |  | 92,684 |  |  |
| RDP | RDP ID | 10,302 | 10,302 | 2,487 | 10,302* |  |
|  | NCBI ID | 10,302 | 2,487 | 2,487 | 2,487 |  |
|  | RNAcentral ID | 10,302 | 4,779 | 2,487 | 4,779* |  |
|  | Total Number of<br>Rows | 10,302 |  | 2,487 |  |  |
| GTDB | GTDB ID | 12,477; 584,382 | 12,477;<br>584,382 | 92,444 | 596,859* |  |
|  | NCBI ID | 12,477; 584,382 | 2,768; 89,701 | 92,444 | 92,444 |  |
|  | Total Number of<br>Rows | 12,477; 584,382 |  | 92,444 |  |  |

\*represent this column contains multiple IDs in one row Values separated by semicolons indicate data from multiple files

**Figure S6.**
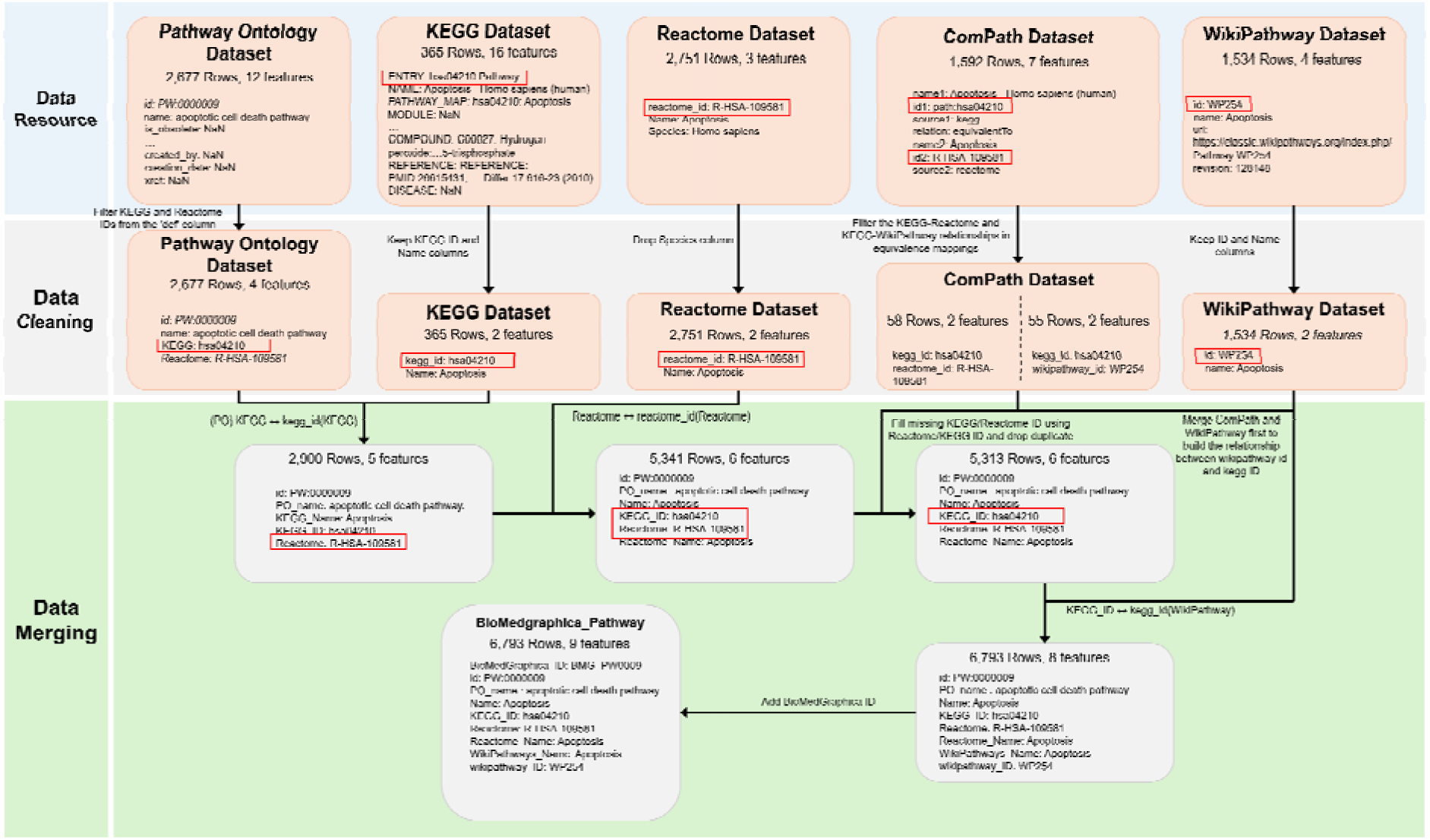
Details of Microbiota Entity Merging Process.

**Figure S6** provides a detailed illustration of the integration process for BioMedGraphica Microbiota, using NCBI Taxon ID: 1105100 as an example. The “Data Resource” section represents the raw data from the databases used in the integration of the microbiota entity. The “Data Cleaning” section shows the format of the cleaned data prepared for integration. The data highlighted in the red boxes indicates the key matching columns used for merging. In the “Data Merging” section, the gray boxes display the data format after each step of database integration. The microbiota entity integration process follows an outer join strategy, first merging data from the NCBI Taxonomy and SILVA databases, followed by integration with Greengenes, RDP, and GTDB in sequence. Finally, the NCBI Taxon ID is used as the primary unit for data unification, consolidating all entries with the same NCBI Taxon ID.

#### Exposure Entity Integration

The data integration for this entity was based on the CTD database. The construction of the exposure entity primarily depends on the Exposure-Study and Exposure-Event associations available in the Comparative Toxicogenomics Database (CTD). By extracting and integrating records based on the ExposureStressorID (corresponding to MeSH identifiers) from both datasets, an initial version of the exposure entity was assembled. Subsequently, each MeSH ID was annotated with it corresponding CSA RN, resulting in the finalized version of the entity. (check **Figure S7** in supplementar section for merging process and results in **Table S9**).

**Table S9.** Exposure Entities Information.

| Database |  | Raw Data |  | After Data Cleaning |  | Total Number of BioMedGraphica ID / BioMedGraphica Connected ID |
| --- | --- | --- | --- | --- | --- | --- |
|  |  | Total Number of Rows | Unique | Total Number of Rows | Unique |  |
| CTD | MeSH ID | 3,539; 224,304; 179,336 | 1,523; 1,159; 179,336 | 1,159 | 1,159 | 1,159 / 1,037 |
|  | CAS ID | NA; NA; 179,336 | NA; NA; 56,642 | 748 | 748 |  |
|  | Total Number of Rows | 3,539; 224,304; 179,336 |  | 1,159 |  |  |

\*represent this column contains multiple IDs in one row
Values separated by semicolons indicate data from multiple files

**Figure S7.**
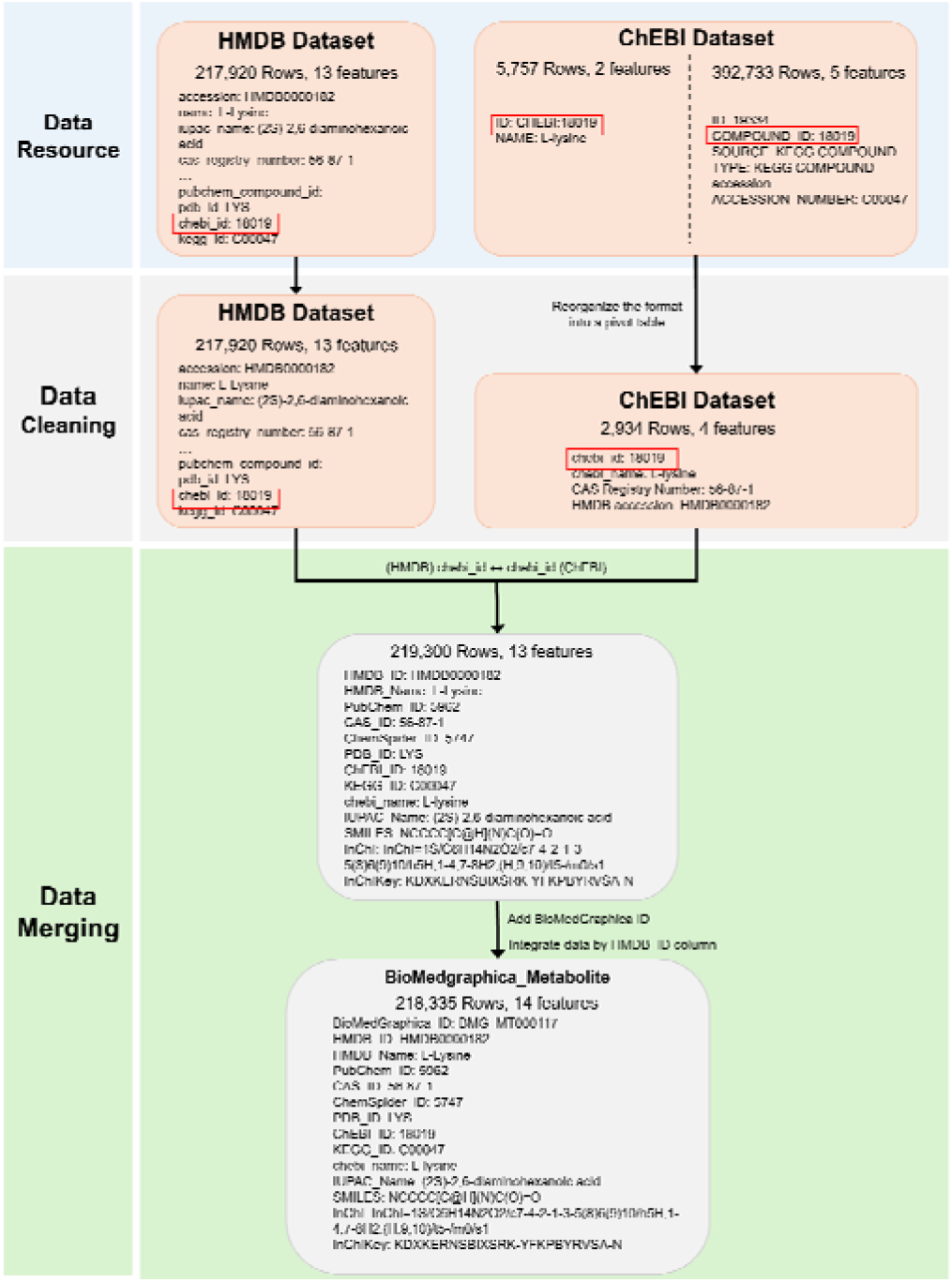
Details of Exposure Entity Merging Process.

**Figure S7** uses MeSH ID D002219 as an example to illustrate the integration process for BioMedGraphica Exposure. The “Data Resource” section displays the raw data from databases used for the exposure entity. The “Data Cleaning” section shows the cleaned data format prepared for integration. The data highlighted in the red boxes indicates the key matching columns used for merging. In the “Data Merging” section, the gray boxes display the data format after each step of database integration.

#### Phenotype Entity Merging

The integration of phenotype entities was performed using data from two primary sources: the Human Phenotype Ontology (HPO) and the Unified Medical Language System (UMLS). From the HPO dataset, we retained the HPO identifiers along with their cross-references to UMLS concepts; similarly, UMLS data were processed to extract corresponding mappings. The datasets were then merged using a left join based on the HPO identifiers, with HPO serving as the primary source. To ensure data consistency and avoid duplication, we further validated the uniqueness of each HPO ID in the resulting entity set. Columns highlighted in bold within the table denote those used for database merging, ensuring the uniqueness of IDs in these columns. (see **Figure S8** in the supplementary section for details on the merging workflow and **Table S10** for results).

**Table S10.** Phenotype entities database description.

| Database |  | Raw Data |  | After Data Cleaning |  | Total Number of BioMedGraphica ID / BioMedGraphica Connected ID |
| --- | --- | --- | --- | --- | --- | --- |
|  |  | Total Number of Rows | Unique | Total Number of Rows | Unique |  |
| HPO | HPO ID | 19,533 | 19,533 | 19,532 | 19,532 | 19,532 / 19,078 |
|  | UMLS ID | 19,532 | 12,786 | 19,532 | 12,786* |  |
|  | Total Number of Rows | 19,533 |  | 19,532 |  |  |
| UMLS | UMLS ID | 14,036,386 | 3,211,875 | 14,914 | 15,958* |  |
|  | HPO ID | 40,197 | 17,664 | 14,914 | 14,914 |  |
|  | Total Number of Rows | 14,036,386 |  | 14,914 |  |  |

\*represent this column contains multiple IDs in one row

**Figure S8.**
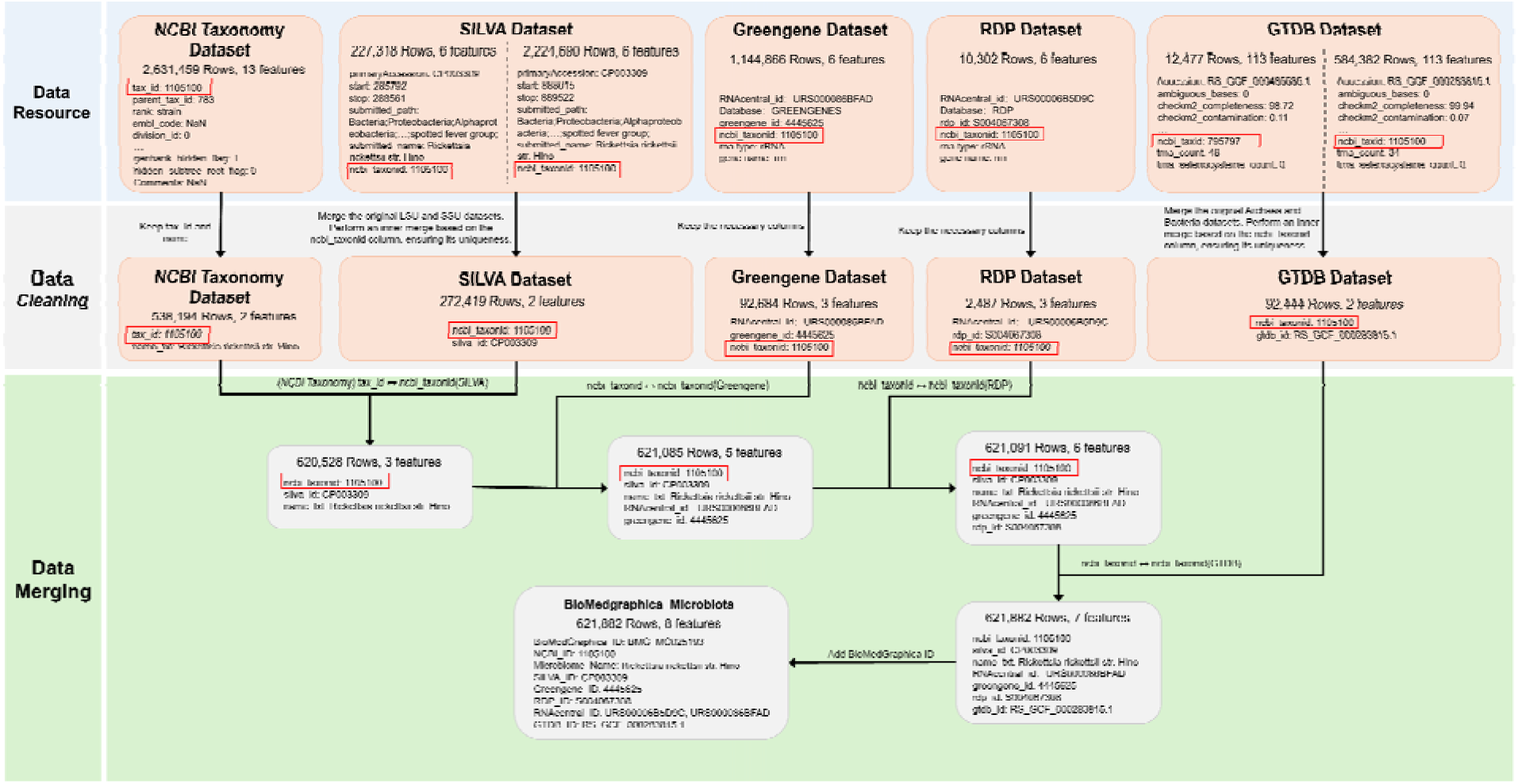
Details of Phenotype Entity Merging Process.

**Figure S8** provides a detailed illustration of the integration process for BioMedGraphica Phenotype, using HP: 0200101 as an example. The “Data Resource” section represents the raw data from the databases used in the integration of the phenotype entity. The “Data Cleaning” section displays the format of the cleaned data prepared for integration. For HPO, descriptive terms in the original names were removed. The data highlighted in the red boxes indicates the key matching columns used for merging. In the “Data Merging” section, the gray boxes show the format of the data after each step of database integration. The phenotype entity integration process follows an outer join approach, merging data from the HPO and UMLS databases.

#### Disease Entity Integration

The integration of disease entities began with the alignment of datasets from UMLS and MeSH. This was followed by the incorporation of mappings to ICD-10 and ICD-11, enabling further consolidation of disease codes. Using relationships provided by the Disease Ontology, identifiers from UMLS, MeSH, and ICD-10 were mapped to corresponding Disease Ontology (DO) terms to enrich the dataset with DO IDs. Missing UMLS data were supplemented via cross-references to SNOMED-CT. Finally, SNOMED-CT data were integrated to provide descriptive labels (names) for all previously collected SNOMED-CT identifiers; both SNOMED-CT IDs and their associated names were retained. Additionally, Mondo Disease Ontology data were merged using its mappings to UMLS and MeSH. Throughout the integration process, the UMLS ID served as the primary unit of granularity, ensuring unique identification across the dataset. Bolded columns in the accompanying table indicate fields used as primary keys during data merging; all identifiers in these fields are uniquely assigned. (refer to **Figure S9** in the supplementary section for a detailed workflow and **Table S11** for results).

**Table S11.**
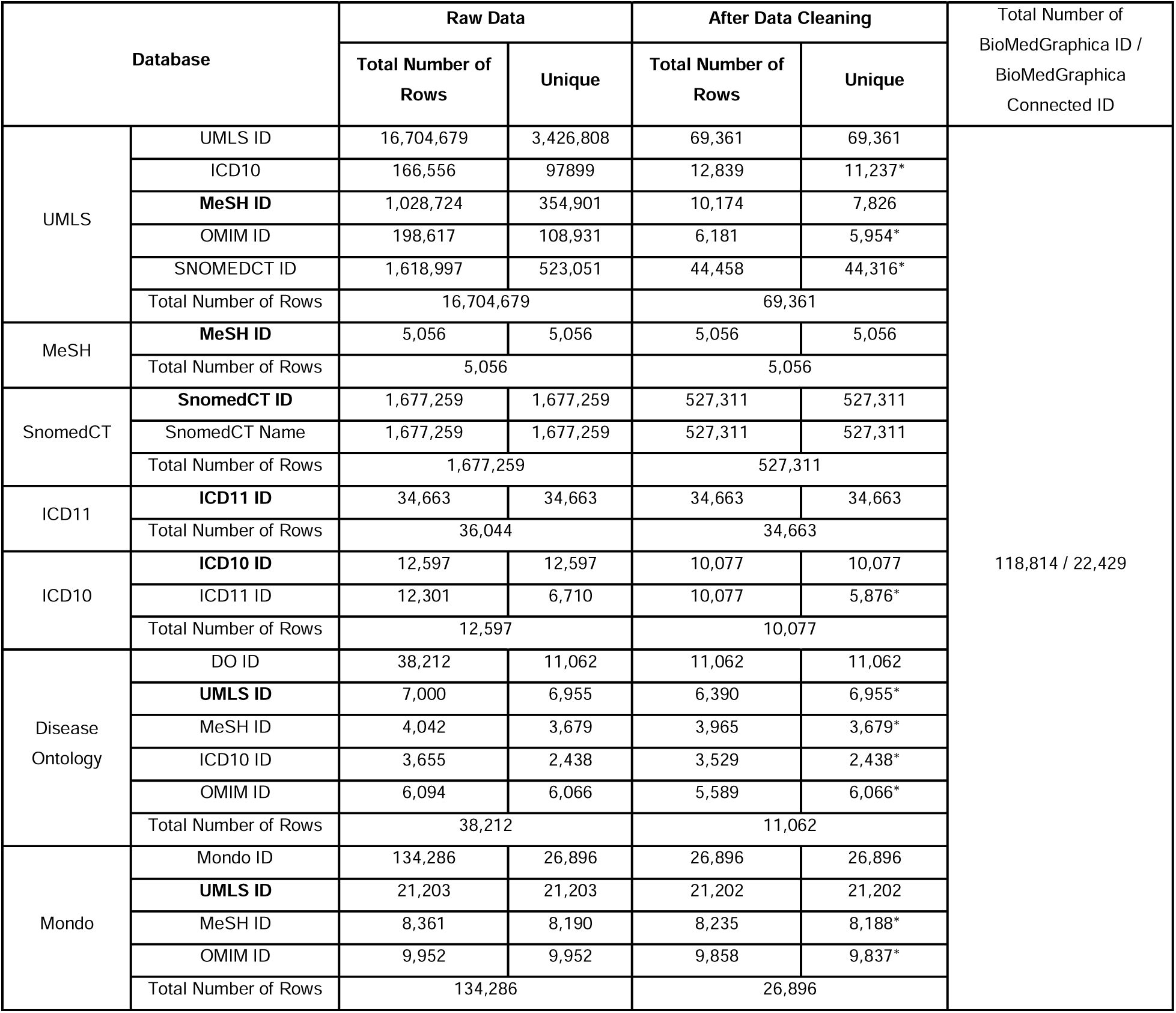
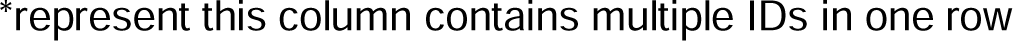
Disease Entity Information.

| Database |  | Raw Data |  | After Data Cleaning |  | Total Number of BioMedGraphica ID / BioMedGraphica Connected ID |
| --- | --- | --- | --- | --- | --- | --- |
|  |  | Total Number of Rows | Unique | Total Number of Rows | Unique |  |
| UMLS | UMLS ID | 16,704,679 | 3,426,808 | 69,361 | 69,361 | 118,814 / 22,429 |
|  | ICD10 | 166,556 | 97899 | 12,839 | 11,237* |  |
|  | MeSH ID | 1,028,724 | 354,901 | 10,174 | 7,826 |  |
|  | OMIM ID | 198,617 | 108,931 | 6,181 | 5,954* |  |
|  | SNOMEDCT ID | 1,618,997 | 523,051 | 44,458 | 44,316* |  |
|  | Total Number of Rows | 16,704,679 |  | 69,361 |  |  |
| MeSH | MeSH ID | 5,056 | 5,056 | 5,056 | 5,056 |  |
|  | Total Number of Rows | 5,056 |  | 5,056 |  |  |
| SnomedCT | SnomedCT ID | 1,677,259 | 1,677,259 | 527,311 | 527,311 |  |
|  | SnomedCT Name | 1,677,259 | 1,677,259 | 527,311 | 527,311 |  |
|  | Total Number of Rows | 1,677,259 |  | 527,311 |  |  |
| ICD11 | ICD11 ID | 34,663 | 34,663 | 34,663 | 34,663 |  |
|  | Total Number of Rows | 36,044 |  | 34,663 |  |  |
| ICD10 | ICD10 ID | 12,597 | 12,597 | 10,077 | 10,077 |  |
|  | ICD11 ID | 12,301 | 6,710 | 10,077 | 5,876* |  |
|  | Total Number of Rows | 12,597 |  | 10,077 |  |  |
| Disease Ontology | DO ID | 38,212 | 11,062 | 11,062 | 11,062 |  |
|  | UMLS ID | 7,000 | 6,955 | 6,390 | 6,955* |  |
|  | MeSH ID | 4,042 | 3,679 | 3,965 | 3,679* |  |
|  | ICD10 ID | 3,655 | 2,438 | 3,529 | 2,438* |  |
|  | OMIM ID | 6,094 | 6,066 | 5,589 | 6,066* |  |
|  | Total Number of Rows | 38,212 |  | 11,062 |  |  |
| Mondo | Mondo ID | 134,286 | 26,896 | 26,896 | 26,896 |  |
|  | UMLS ID | 21,203 | 21,203 | 21,202 | 21,202 |  |
|  | MeSH ID | 8,361 | 8,190 | 8,235 | 8,188* |  |
|  | OMIM ID | 9,952 | 9,952 | 9,858 | 9,837* |  |
|  | Total Number of Rows | 134,286 |  | 26,896 |  |  |

\*represent this column contains multiple IDs in one row

**Figure S9.**
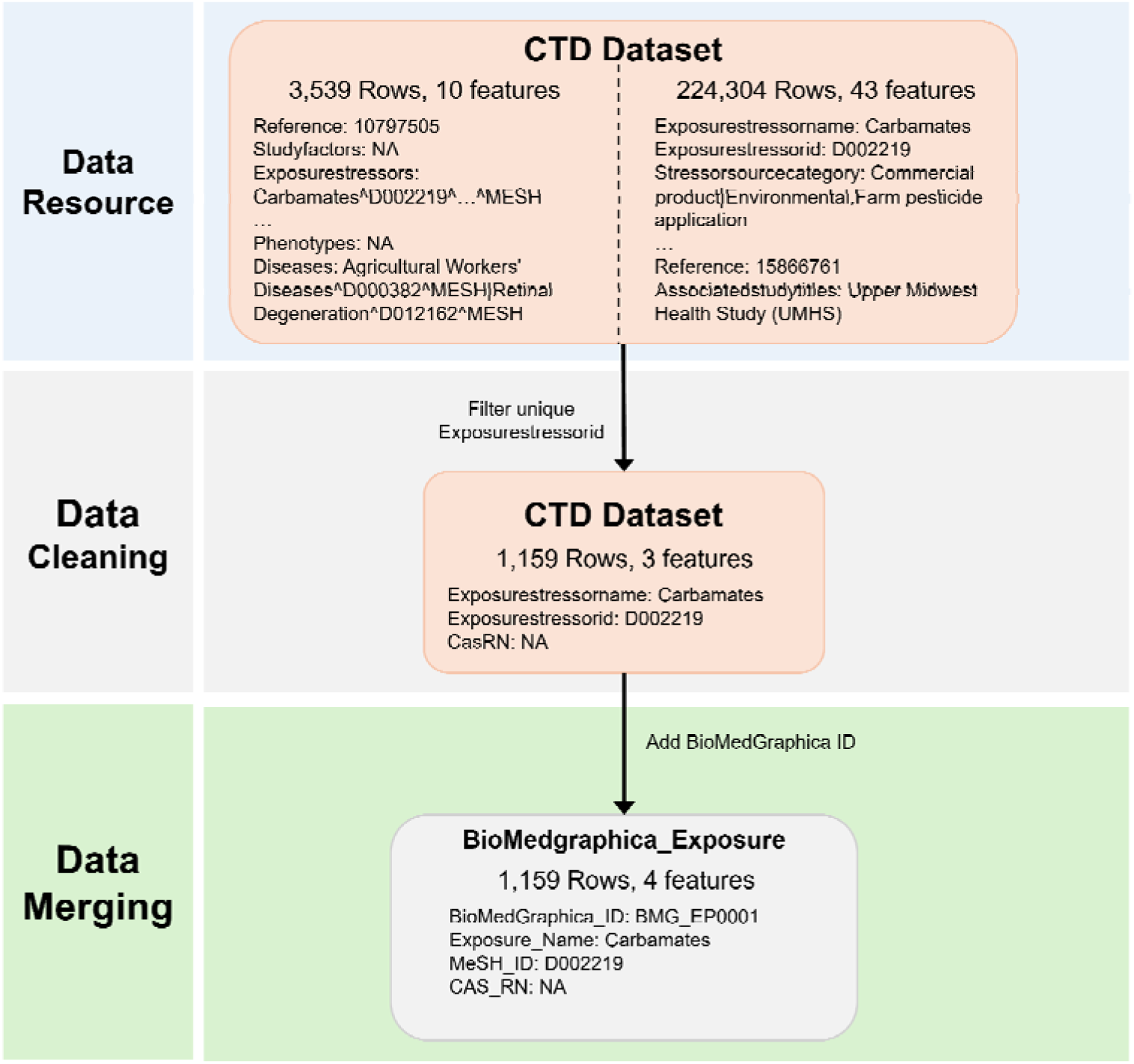
Details of Disease Entity Merging Process.

**Figure S9** provides a detailed illustration of the integration process for BioMedGraphica Disease, using C0000744 as an example. The “Data Resource” section represents the raw data from the databases used in the integration of the disease entity. The “Data Cleaning” section shows the format of the cleaned data prepared for integration. The data highlighted in the red boxes indicates the key matching columns used for merging. In the “Data Merging” section, the gray boxes display the data format after each step of database integration. The disease entity integration process follows an outer join methodology. Initially, MeSH and UMLS databases were merged to construct the foundational dataset. This was followed by the sequential integration of ICD-10, ICD-11, Disease Ontology (DO), and Mondo, each contributing additional mappings and identifiers. In the final step, SNOMED CT data were incorporated to supplement the dataset with SNOMED CT identifiers and their corresponding concept names. Finally, the UMLS ID serves as the primary identifier for data unification, consolidating all entries with the same UMLS ID.

#### Drug Entity Merging

The integration process commenced by merging NDC and UNII datasets using the SUBSTANCENAME as the key identifier. PubChem data was then incorporated through its mapping with PubChem CIDs. DrugBank data was integrated next, utilizing mappings between DrugBank IDs, CAS numbers, and SIDs. Finally, any missing data within the same row was supplemented using synonyms from both PubChem and DrugBank. After integration, internal deduplication was performed using PubChem CIDs and CAS RNs in sequence to ensure consistency. The CAS number was designated a the minimal unit of data granularity for this entity. Bolded entries in the table indicate the columns used for merging across databases, where IDs in these columns are uniquely assigned (refer to **Figure S10** in the supplementary section for details on the merging process and **Table S12** for the results).

**Table S12.**
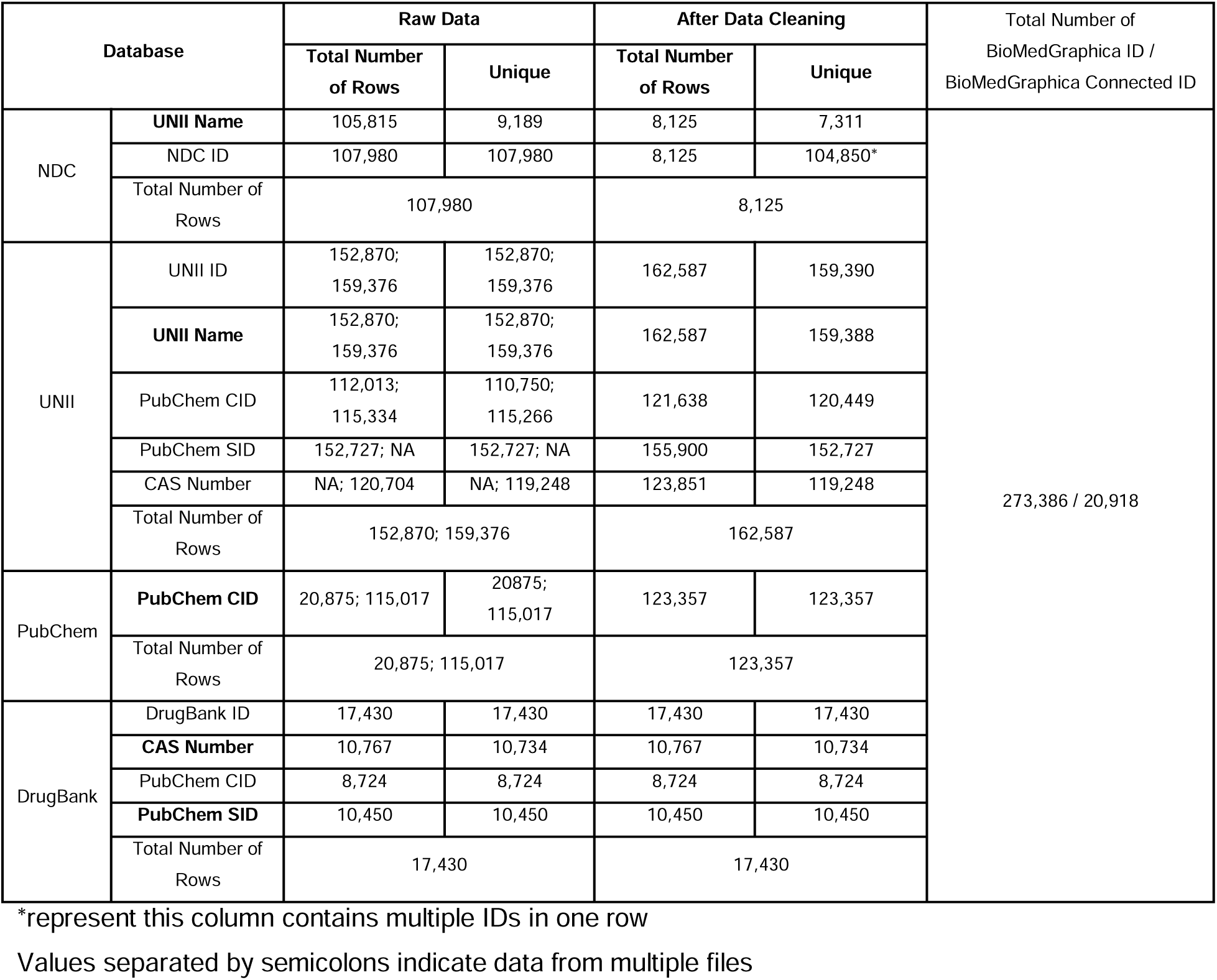
Drug Entity Information.

**Figure S10.**
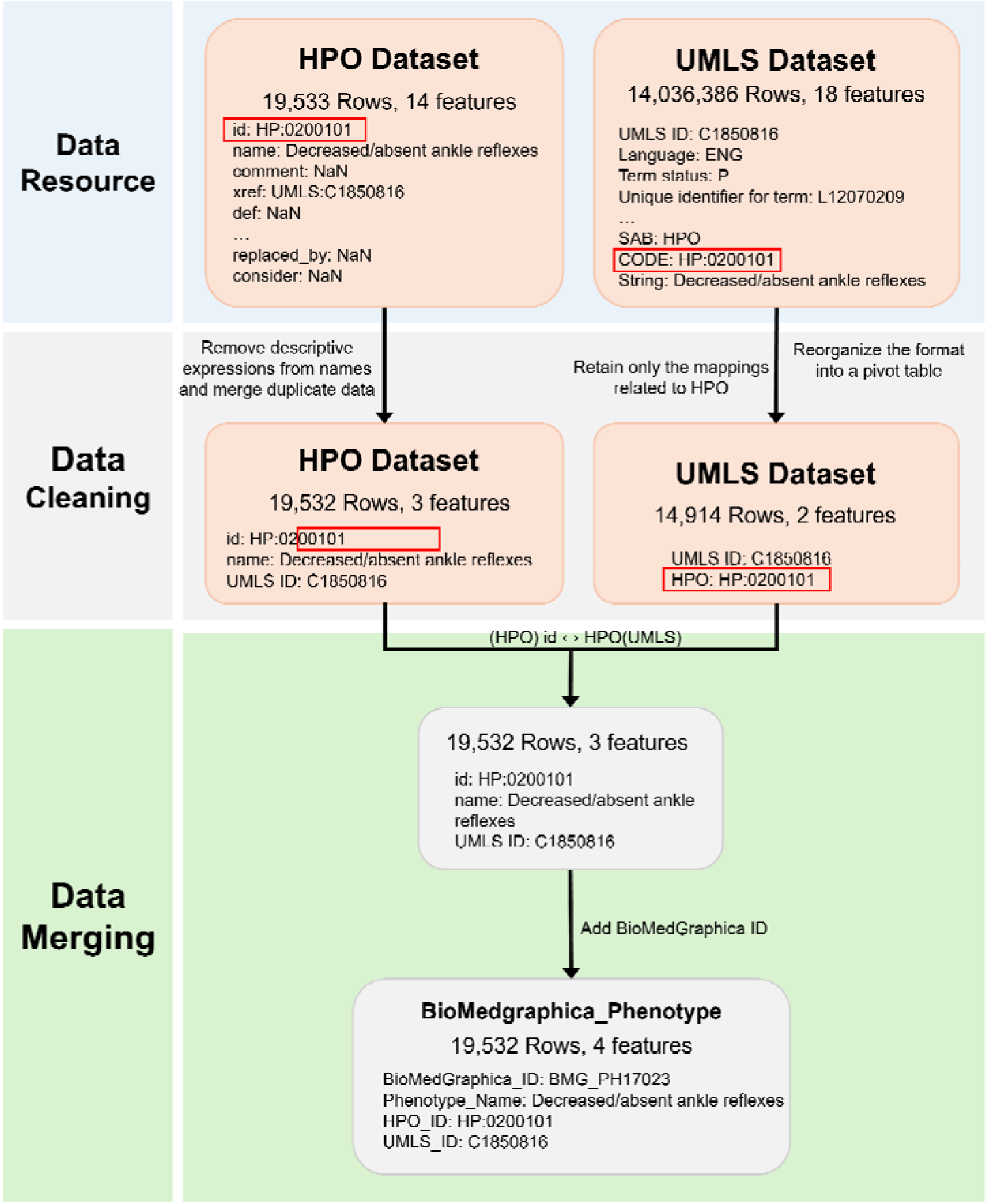
Details of Drug Entity Merging Process.

**Figure S10** provides a detailed illustration of the integration process for BioMedGraphica Drug, using PubChem CID: 10034073 as an example. The “Data Resource” section represents the raw data from the databases used in the integration of the drug entity. The “Data Cleaning” section displays the format of the cleaned data prepared for integration. The data highlighted in the red boxes indicate the key matching columns used for merging. In the “Data Merging” section, the gray boxes show the format of the data after each database integration step. The drug entity integration process follows an outer join approach. First, the NDC and UNII databases are merged, followed by integrating the combined data with the PubChem database, and then DrugBank. Finally, the CAS number is used as the primary identifier for data unification, ensuring all entries with the same CAS number are consolidated.

## References

1. TCGA. https://www.cancer.gov/about-nci/organization/ccg/research/structural-genomics/tcga.

2. Barretina J, Caponigro G, Stransky N, et al. The Cancer Cell Line Encyclopedia enables predictive modelling of anticancer drug sensitivity. Nature. 2012;483(7391):603-607. doi:10.1038/nature11003

3. Yang W, Soares J, Greninger P, et al. Genomics of Drug Sensitivity in Cancer (GDSC): A resource for therapeutic biomarker discovery in cancer cells. Nucleic Acids Res. 2013;41(D1):D955–D961. doi:10.1093/nar/gks1111

4. Ghandi M, Huang FW, Jané-Valbuena J, et al. Next-generation characterization of the Cancer Cell Line Encyclopedia. Nature. 2019;569(7757):503-508. doi:10.1038/s41586-019-1186-3

5. Bennett DA, Buchman AS, Boyle PA, Barnes LL, Wilson RS, Schneider JA. Religious Orders Study and Rush Memory and Aging Project. J Alzheimers Dis. 2018;64(s1):S161-S189. doi:10.3233/JAD-179939

6. Deelen J, Evans DS, Arking DE, et al. A meta-analysis of genome-wide association studies identifies multiple longevity genes. Nat Commun. 2019;10(1):3669. doi:10.1038/s41467-019-11558-2

7. Zhang H, Xu T, Cao D, et al. OmniCellTOSG: The First Cell Text-Omic Signaling Graphs Dataset for Joint LLM and GNN Modeling. arXiv preprint arXiv:250402148. Published online 2025.

8. Program CZICS, Abdulla S, Aevermann B, et al. CZ CELLxGENE Discover: a single-cell data platform for scalable exploration, analysis and modeling of aggregated data. Nucleic Acids Res. 2025;53(D1):D886–D900. doi:10.1093/nar/gkae1142

9. Rood JE, Wynne S, Robson L, et al. The Human Cell Atlas from a cell census to a unified foundation model. Nature. Published online 2024. doi:10.1038/s41586-024-08338-4

10. Heimberg G, Kuo T, DePianto DJ, et al. A cell atlas foundation model for scalable search of similar human cells. Nature. Published online 2024. doi:10.1038/s41586-024-08411-y

11. Cui H, Wang C, Maan H, et al. scGPT: toward building a foundation model for single-cell multi-omics using generative AI. Nat Methods. 2024;21(8):1470–1480. doi:10.1038/s41592-024-02201-0

12. Gottweis J, Weng WH, Daryin A, et al. Towards an AI co-scientist. arXiv preprint arXiv:250218864. Published online 2025.

13. Huang D, Li H, Li W, et al. OmniCellAgent: Towards AI Co-Scientists for Scientific Discovery in Precision Medicine. bioRxiv. Published online 2025:2025–2027.

14. Sapkota R, Roumeliotis KI, Karkee M. Ai agents vs. agentic ai: A conceptual taxonomy, applications and challenges. arXiv preprint arXiv:250510468. Published online 2025.

15. Huang Y, Chen Y, Zhang H, et al. Deep Research Agents: A Systematic Examination And Roadmap. arXiv preprint arXiv:250618096. Published online 2025.

16. Wang H, He Y, Coelho PP, et al. SpatialAgent: An Autonomous AI Agent for Spatial Biology. bioRxiv. Published online 2025:2024-2025.

17. Wei J, Wang X, Schuurmans D, et al. Chain-of-thought prompting elicits reasoning in large language models. Adv Neural Inf Process Syst. 2022;35:24824–24837.

18. Tan X, Wang H, Qiu X, et al. Struct-X: Enhancing Large Language Models Reasoning with Structured Data. arXiv preprint arXiv:240712522. Published online 2024.

19. Jiang J, Zhou K, Dong Z, Ye K, Zhao WX, Wen JR. Structgpt: A general framework for large language model to reason over structured data. arXiv preprint arXiv:230509645. Published online 2023.

20. Hulsen T, Jamuar SS, Moody AR, et al. From big data to precision medicine. Front Med (Lausanne).Frontiers Media S.A. 2019;6(MAR). doi:10.3389/fmed.2019.00034

21. Kendall TJ, Jimenez-Ramos M, Turner F, et al. An integrated gene-to-outcome multimodal database for metabolic dysfunction-associated steatotic liver disease. Nat Med. 2023;29(11):2939–2953. doi:10.1038/s41591-023-02602-2

22. Fernández-Torras A, Duran-Frigola M, Bertoni M, Locatelli M, Aloy P. Integrating and formatting biomedical data as pre-calculated knowledge graph embeddings in the Bioteque. Nat Commun. 2022;13(1). doi:10.1038/s41467-022-33026-0

23. Königs C, Friedrichs M, Dietrich T. The heterogeneous pharmacological medical biochemical network PharMeBINet. Sci Data. 2022;9(1). doi:10.1038/s41597-022-01510-3

24. Ma T, Zhang A. Integrate multi-omics data with biological interaction networks using Multi-view Factorization AutoEncoder (MAE). BMC Genomics. 2019;20. doi:10.1186/s12864-019-6285-x

25. Cavalleri E, Cabri A, Soto-Gomez M, et al. An ontology-based knowledge graph for representing interactions involving RNA molecules. Sci Data. 2024;11(1). doi:10.1038/s41597-024-03673-7

26. MacNamara A, Nakic N, Amin Al Olama A, et al. Network and pathway expansion of genetic disease associations identifies successful drug targets. Sci Rep. 2020;10(1). doi:10.1038/s41598-020-77847-9

27. Zhang H, Cao D, Xu T, et al. mosGraphFlow: a novel integrative graph AI model mining disease targets from multi-omic data. bioRxiv. Published online January 1, 2024:2024.08.01.606219. doi:10.1101/2024.08.01.606219

28. Zhang H, Huang D, Chen E, et al. mosGraphGPT: a foundation model for multi-omic signaling graphs using generative AI. bioRxiv. Published online January 1, 2024:2024.08.01.606222. doi:10.1101/2024.08.01.606222

29. Zhang H, Cao D, Chen Z, et al. mosGraphGen: a novel tool to generate multi-omics signaling graphs to facilitate integrative and interpretable graph AI model development. Bioinformatics Advances. 2024;4(1):vbae151. doi:10.1093/bioadv/vbae151

30. Zhang H, Goedegebuure SP, Ding L, et al. M3NetFlow: A multi-scale multi-hop graph AI model for integrative multi-omic data analysis. iScience. 2025;28(3):111920. 10.1016/j.isci.2025.111920

31. Feng J, Province M, Li G, Payne PRO, Chen Y, Li F. PathFinder: a novel graph transformer model to infer multi-cell intra- and inter-cellular signaling pathways and communications. Front Cell Neurosci. Published online January 1, 2024:2024.01.13.575534. doi:10.1101/2024.01.13.575534

32. Dong Z, Zhao Q, Payne PRO, et al. Highly accurate disease diagnosis and highly reproducible biomarker identification with PathFormer.. https://www.ncbi.nlm.nih.gov/pmc/articles/PMC10680938/. Published online 2023.

33. Zhang H, Chen Y, Payne P, Li F. Using DeepSignalingFlow to mine signaling flows interpreting mechanism of synergy of cocktails. NPJ Syst Biol Appl. 2024;10(1):92. doi:10.1038/s41540-024-00421-w

34. Ma S, Zeng AGX, Haibe-Kains B, Goldenberg A, Dick JE, Wang B. Moving towards genome-wide data integration for patient stratification with Integrate Any Omics. Nat Mach Intell. 2025;7(1):29–42.

35. Zhang H, Cao D, Chen Z, et al. mosGraphGen: a novel tool to generate multi-omic signaling graphs to facilitate integrative and interpretable graph AI model development. doi:10.1101/2024.05.15.594360

36. Ma S, Zeng AG, Haibe-Kains B, Goldenberg A, Dick JE, Wang B. Integrate Any Omics: Towards Genome-Wide Data Integration for Patient Stratification.; 2024.

37. Lee J, Yoon W, Kim S, et al. BioBERT: A pre-trained biomedical language representation model for biomedical text mining. Bioinformatics. 2020;36(4):1234–1240. doi:10.1093/bioinformatics/btz682

38. Zhang H, Huang D, Chen Y, Li F. GraphSeqLM: A Unified Graph Language Framework for Omic Graph Learning. In: Companion Proceedings of the ACM on Web Conference 2025. 2025:1510–1513.

39. Park JS, O’Brien J, Cai CJ, Morris MR, Liang P, Bernstein MS. Generative agents: Interactive simulacra of human behavior. In: Proceedings of the 36th Annual Acm Symposium on User Interface Software and Technology. 2023:1–22.

40. Howe KL, Achuthan P, Allen J, et al. Ensembl 2021. Nucleic Acids Res. 2021;49(D1):D884–D891. doi:10.1093/nar/gkaa942

41. Amberger J, Bocchini CA, Scott AF, Hamosh A. McKusick’s Online Mendelian Inheritance in Man (OMIM®). Nucleic Acids Res. 2009;37(SUPPL. 1). doi:10.1093/nar/gkn665

42. Povey S, Lovering R, Bruford E, Wright M, Lush M, Wain H. The HUGO Gene Nomenclature Committee (HGNC). Hum Genet. 2001;109(6):678–680. doi:10.1007/s00439-001-0615-0

43. Schoch CL, Ciufo S, Domrachev M, et al. NCBI Taxonomy: A comprehensive update on curation, resources and tools. Database.Oxford University Press. 2020;2020. doi:10.1093/database/baaa062

44. O’Leary NA, Wright MW, Brister JR, et al. Reference sequence (RefSeq) database at NCBI: Current status, taxonomic expansion, and functional annotation. Nucleic Acids Res. 2016;44(D1):D733–D745. doi:10.1093/nar/gkv1189

45. Sweeney BA, Petrov AI, Burkov B, et al. RNAcentral: A hub of information for non-coding RNA sequences. Nucleic Acids Res. 2019;47(D1):D221–D229. doi:10.1093/nar/gky1034

46. Wu CH, Apweiler R, Bairoch A, et al. The Universal Protein Resource (UniProt): an expanding universe of protein information. Nucleic Acids Res. 2006;34(Database issue). doi:10.1093/nar/gkj161

47. Fabregat A, Jupe S, Matthews L, et al. The reactome pathway knowledgebase. Nucleic Acids Res. 2018;46(D1):D649–D655.

48. Kanehisa MGS, Goto S. KEGG: kyoto Encyclopedia of Genes and Genomes. Nucleic Acids Res. 2000;28:27–30. doi:10.1093/nar/28.1.27

49. Kelder T, Van Iersel MP, Hanspers K, et al. WikiPathways: building research communities on biological pathways. Nucleic Acids Res. 2012;40(D1):D1301–D1307.

50. Petri V, Jayaraman P, Tutaj M, et al. The pathway ontology–updates and applications. J Biomed Semantics. 2014;5:1–12.

51. Domingo-Fernández D, Hoyt CT, Bobis-Álvarez C, Marín-Llaó J, Hofmann-Apitius M. ComPath: an ecosystem for exploring, analyzing, and curating mappings across pathway databases. NPJ Syst Biol Appl. 2018;4(1):43.

52. Wishart DS, Guo A, Oler E, et al. HMDB 5.0: the human metabolome database for 2022. Nucleic Acids Res. 2022;50(D1):D622–D631.

53. Degtyarenko K, De matos P, Ennis M, et al. ChEBI: A database and ontology for chemical entities of biological interest. Nucleic Acids Res. 2008;36(SUPPL. 1). doi:10.1093/nar/gkm791

54. Quast C, Pruesse E, Yilmaz P, et al. The SILVA ribosomal RNA gene database project: improved data processing and web-based tools. Nucleic Acids Res. 2012;41(D1):D590–D596.

55. DeSantis TZ, Hugenholtz P, Larsen N, et al. Greengenes, a chimera-checked 16S rRNA gene database and workbench compatible with ARB. Appl Environ Microbiol. 2006;72(7):5069–5072.

56. Cole JR, Wang Q, Fish JA, et al. Ribosomal Database Project: data and tools for high throughput rRNA analysis. Nucleic Acids Res. 2014;42(D1):D633–D642.

57. Parks DH, Chuvochina M, Waite DW, et al. A standardized bacterial taxonomy based on genome phylogeny substantially revises the tree of life. Nat Biotechnol. 2018;36(10):996–1004.

58. Davis AP, Grondin CJ, Johnson RJ, et al. Comparative Toxicogenomics Database (CTD): Update 2021. Nucleic Acids Res. 2021;49(D1):D1138–D1143. doi:10.1093/nar/gkaa891

59. Köhler S, Gargano M, Matentzoglu N, et al. The human phenotype ontology in 2021. Nucleic Acids Res. 2021;49(D1):D1207–D1217.

60. Bodenreider O. The unified medical language system (UMLS): integrating biomedical terminology. Nucleic Acids Res. 2004;32(suppl_1):D267–D270.

61. Organization WH. International Statistical Classification of Diseases and Related Health Problems: Alphabetical Index. Vol 3. World Health Organization; 2004.

62. Organization WH. International classification of diseases for mortality and morbidity statistics (11th Revision). Preprint posted online 2018.

63. Schriml LM, Arze C, Nadendla S, et al. Disease ontology: A backbone for disease semantic integration. Nucleic Acids Res. 2012;40(D1). doi:10.1093/nar/gkr972

64. Lipscomb CE. Medical subject headings (MeSH). Bull Med Libr Assoc. 2000;88(3):265.

65. Donnelly K. SNOMED-CT: The advanced terminology and coding system for eHealth. Stud Health Technol Inform. 2006;121:279.

66. Vasilevsky N, Essaid S, Matentzoglu N, et al. Mondo Disease Ontology: harmonizing disease concepts across the world. In: CEUR Workshop Proceedings, CEUR-WS. Vol 2807. 2020.

67. Wang Y, Xiao J, Suzek TO, Zhang J, Wang J, Bryant SH. PubChem: A public information system for analyzing bioactivities of small molecules. Nucleic Acids Res. 2009;37(SUPPL. 2). doi:10.1093/nar/gkp456

68. Tribble DA. The National Drug Code explained. American Journal of Health-System Pharmacy. Published online 2024:zxae274.

69. Weisgerber DW. Chemical abstracts service chemical registry system: history, scope, and impacts. Journal of the American Society for Information Science. 1997;48(4):349–360.

70. Knox C, Wilson M, Klinger CM, et al. DrugBank 6.0: the DrugBank Knowledgebase for 2024. Nucleic Acids Res. 2024;52(D1):D1265–D1275. doi:10.1093/nar/gkad976

71. Stark C, Breitkreutz BJ, Reguly T, Boucher L, Breitkreutz A, Tyers M. BioGRID: a general repository for interaction datasets. Nucleic Acids Res. 2006;34(suppl_1):D535–D539.

72. Oughtred R, Stark C, Breitkreutz BJ, et al. The BioGRID interaction database: 2019 update. Nucleic Acids Res. 2019;47(D1):D529–D541.

73. Szklarczyk D, Franceschini A, Wyder S, et al. STRING v10: protein–protein interaction networks, integrated over the tree of life. Nucleic Acids Res. 2015;43(D1):D447–D452.

74. Szklarczyk D, Gable AL, Lyon D, et al. STRING v11: protein–protein association networks with increased coverage, supporting functional discovery in genome-wide experimental datasets. Nucleic Acids Res. 2019;47(D1):D607–D613.

75. Piñero J, Bravo À, Queralt-Rosinach N, et al. DisGeNET: a comprehensive platform integrating information on human disease-associated genes and variants. Nucleic Acids Res. Published online 2016:gkw943.

76. Pletscher-Frankild S, Pallejà A, Tsafou K, Binder JX, Jensen LJ. DISEASES: Text mining and data integration of disease-gene associations. Methods. 2015;74:83–89. doi:10.1016/j.ymeth.2014.11.020

77. Moretti S, Tran VDT, Mehl F, Ibberson M, Pagni M. MetaNetX/MNXref: unified namespace for metabolites and biochemical reactions in the context of metabolic models. Nucleic Acids Res. 2021;49(D1):D570–D574.

78. Janssens Y, Nielandt J, Bronselaer A, et al. Disbiome database: linking the microbiome to disease. BMC Microbiol. 2018;18:1–6.

79. Sun YZ, Zhang DH, Cai SB, Ming Z, Li JQ, Chen X. MDAD: a special resource for microbe-drug associations. Front Cell Infect Microbiol. 2018;8:424.

80. Doestzada M, Vila AV, Zhernakova A, et al. Pharmacomicrobiomics: a novel route towards personalized medicine? Protein Cell. 2018;9(5):432–445.

81. Gilson MK, Liu T, Baitaluk M, Nicola G, Hwang L, Chong J. BindingDB in 2015: a public database for medicinal chemistry, computational chemistry and systems pharmacology. Nucleic Acids Res. 2016;44(D1):D1045-D1053.

82. Ursu O, Holmes J, Knockel J, et al. DrugCentral: online drug compendium. Nucleic Acids Res. Published online 2016:gkw993.

83. Kuhn M, Campillos M, Letunic I, Jensen LJ, Bork P. A side effect resource to capture phenotypic effects of drugs. Mol Syst Biol. 2010;6(1):343.

84. Guo T, Yang Q, Wang C, et al. Knowledgenavigator: Leveraging large language models for enhanced reasoning over knowledge graph. Complex & Intelligent Systems. 2024;10(5):7063–7076.

85. Ma T, Tao W, Li M, et al. KGExplainer: Towards Exploring Connected Subgraph Explanations for Knowledge Graph Completion. arXiv preprint arXiv:240403893. Published online 2024.

86. Rajabi E, Etminani K. Knowledge-graph-based explainable AI: A systematic review. J Inf Sci. 2024;50(4):1019–1029.

87. 2022 Alzheimer’s disease facts and figures. Alzheimer’s & Dementia. 2022;18(4):700–789. 10.1002/alz.12638

88. 2018 Alzheimer’s disease facts and figures. Alzheimer’s & Dementia. 2018;14(3):367–429. 10.1016/j.jalz.2018.02.001

89. Cummings J, Lee G, Ritter A, Zhong K. Alzheimer’s disease drug development pipeline: 2018. Alzheimer’s and Dementia: Translational Research and Clinical Interventions. 2018;4:195–214. doi:10.1016/j.trci.2018.03.009

90. AD facts. https://www.alz.org/alzheimers-dementia/facts-figures.

91. Verheijen J, Sleegers K. Understanding Alzheimer Disease at the Interface between Genetics and Transcriptomics. Vol 34.; 2018. doi:10.1016/j.tig.2018.02.007

92. Daichao X, Jin T, Zhu H, et al. TBK1 Suppresses RIPK1-Driven Apoptosis and Inflammation during Development and in Aging. Vol 174.; 2018. doi:10.1016/j.cell.2018.07.041

93. Li F, Eteleeb AM, Buchser W, et al. Weakly activated core neuroinflammation pathways were identified as a central signaling mechanism contributing to the chronic neurodegeneration in Alzheimer’s disease. Front Aging Neurosci. 2022;14. doi:10.3389/fnagi.2022.935279

94. Li F, Oh I, Kumar S, et al. Loss of estrogen unleashing neuro-inflammation increases the risk of Alzheimer&#039;s disease in women. bioRxiv. Published online January 1, 2022:2022.09.19.508592. doi:10.1101/2022.09.19.508592

95. Mizuno S, Iijima R, Ogishima S, et al. AlzPathway: a comprehensive map of signaling pathways of Alzheimer’s disease. BMC Syst Biol. 2012;6:52. doi:10.1186/1752-0509-6-52

96. Godoy JA, Rios JA, Zolezzi JM, Braidy N, Inestrosa NC. Signaling pathway cross talk in Alzheimer’s disease. Cell Commun Signal. 2014;12:23. doi:10.1186/1478-811X-12-23

97. Lazic D, Sagare AP, Nikolakopoulou AM, Griffin JH, Vassar R, Zlokovic B V. 3K3A-activated protein C blocks amyloidogenic BACE1 pathway and improves functional outcome in mice. J Exp Med. Published online January 15, 2019:jem.20181035. doi:10.1084/jem.20181035

98. Ofengeim D, Mazzitelli S, Ito Y, et al. RIPK1 mediates a disease-associated microglial response in Alzheimer’s disease. Proceedings of the National Academy of Sciences. 2017;114(41):E8788 LP–E8797. doi:10.1073/pnas.1714175114

99. Sims R, van der Lee SJ, Naj AC, et al. Rare coding variants in PLCG2, ABI3, and TREM2 implicate microglial-mediated innate immunity in Alzheimer&#39;s disease. Nat Genet. 2017;49:1373.

100. Kunkle BW, Grenier-Boley B, Sims R, et al. Genetic meta-analysis of diagnosed Alzheimer’s disease identifies new risk loci and implicates Aβ, tau, immunity and lipid processing. Nat Genet. 2019;51(3):414–430. doi:10.1038/s41588-019-0358-2

